# A conserved Guided Entry of Tail-anchored pathway is involved in the trafficking of tail-anchored membrane proteins in *Plasmodium falciparum*

**DOI:** 10.1101/2021.05.03.442402

**Authors:** Tarkeshwar Kumar, Satarupa Maitra, Abdur Rahman, Souvik Bhattacharjee

## Abstract

Tail-anchored (TA) proteins are defined by the absence of N-terminus signal sequence and the presence of a single transmembrane domain (TMD) proximal to their extreme C-terminus. They play fundamental roles in cellular processes including vesicular trafficking, protein translocation and quality control. Accordingly, TA proteins are post-translationally integrated by the Guided Entry of TA (GET) pathway to the cellular membranes; with their N-terminus oriented towards the cytosol and C-terminus facing the organellar lumen. The TA repertoire and the GET machinery have been extensively characterized in the yeast and mammalian systems, however, they remain elusive in the human malaria parasite *Plasmodium falciparum.* In this study, we bioinformatically predicted a total of 63 TA proteins in the *P. falciparum* proteome and revealed the association of their subset with the *P. falciparum* homolog of Get3 (PfGet3). In addition, our proximity labelling studies either definitively identified or shortlisted the other eligible GET constituents, and our *in vitro* association studies validated associations between PfGet3 and the corresponding homologs of Get4 and Get2 in *P. falciparum*. Collectively, this study reveals the presence of proteins with hallmark TA signatures and the involvement of evolutionary conserved GET trafficking pathway for their targeted delivery within the parasite.

**Synopsis:** Tail-anchored (TA) proteins, characterized by an absence of N-terminal signal sequence and the presence of a transmembrane domain near the C-terminus, are post-translationally inserted at organellar membranes by the conserved multi-component Guided Entry of TA (GET) pathway. Here, we identified the putative homologs of GET machinery in the human malaria parasite *Plasmodium falciparum* and revealed their association with a subset of bioinformatically predicted 63 putative TA proteins, thereby validating the functional existence of this trafficking pathway within the apicomplexan parasite.

**Graphical Abstract:** 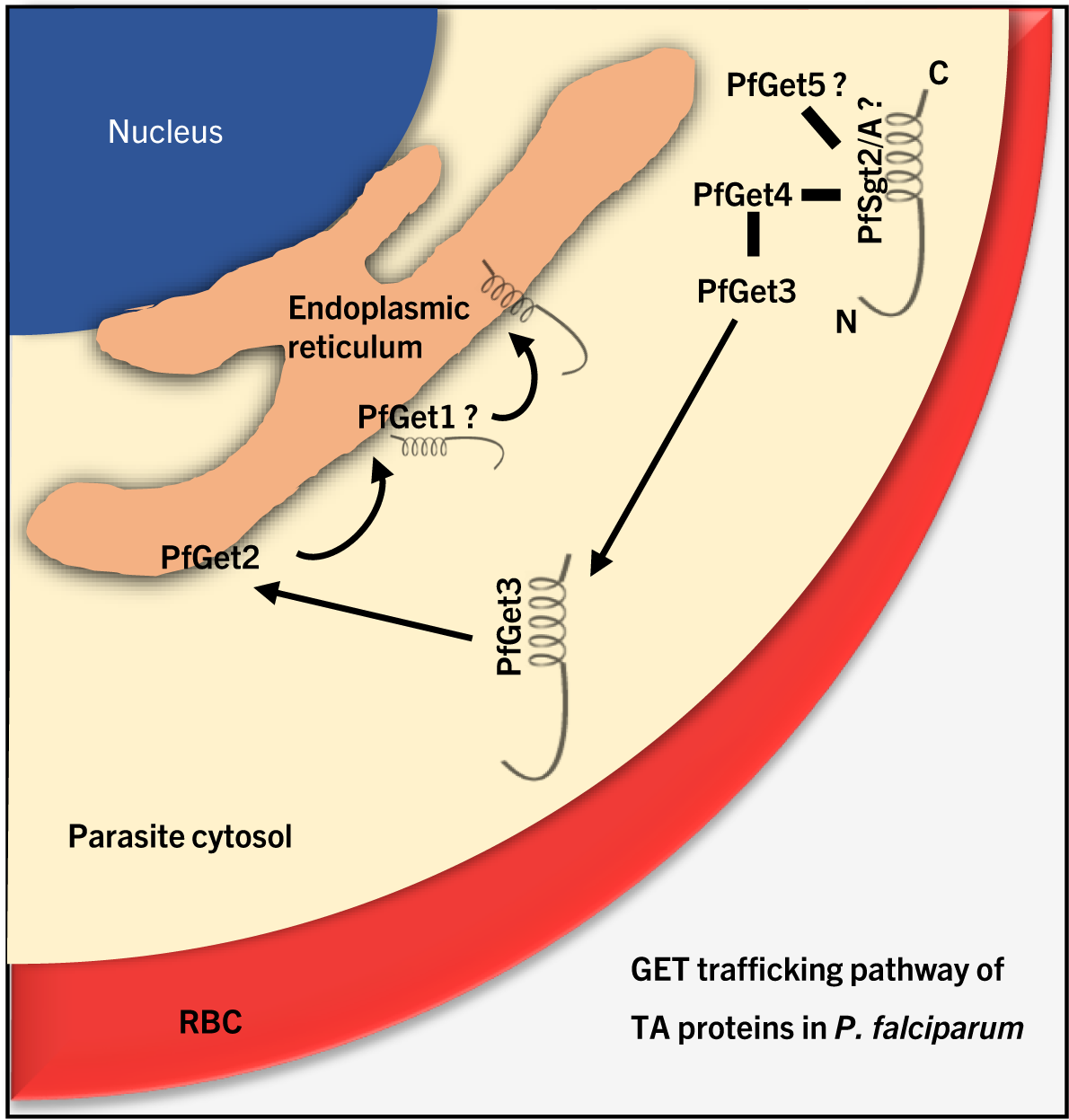

## 1 INTRODUCTION

Integral membrane proteins constitute ∼20-30% of the total eukaryotic proteome where they serve essential cellular functions including vesicular sorting, solute transport, protein homeostasis and organelle biosynthesis. Thus, precise targeting of membrane proteins to their respective subcellular destinations is often dictated by the evolutionary conserved and sophisticated trafficking mechanisms. Most of the membrane proteins are inserted through the chaperone-assisted and co-translational pathway involving recognition of ribosome-associated nascent chains (RNC) by the signal recognition particle (SRP), targeting to the SRP-receptor at the ER membrane [1, 2], and their release to the Sec61 translocon [2–4]. The Sec61 complex subsequently facilitates TMD integration into the lipid bilayer as they emerge out from the ribosomes [5–9]. A major advantage of the co-translational targeting is the tightly coordinated relay of events between protein synthesis, targeting and membrane insertion to ensure efficient shielding of the hydrophobic TMDs from the bulk hydrophilic cytosolic milieu. However, not all membrane proteins recruit the SRP/Sec61 route for insertion. Tail-anchored (TA) proteins are one such unique class of integral membrane proteins characterized by the absence of any N-terminus signal sequence (SS) and the presence of a single helical transmembrane domain (TMD) at or near their C-terminus (CTS) [10]. This close proximity of the TMDs in TA proteins places it within the ribosomal tunnel, thus precluding SRP/Sec61-mediated co-translational insertion, and consequently TA proteins must target in a strictly post-translational manner [11–13]. Notable examples include proteins of the vesicular trafficking pathway (the SNAREs, Soluble NSF Attachment protein REceptors) [14, 15]; ER and mitochondrial subunit translocation machinery [16, 17]; mitochondrial electron carrier (cytochrome b5/Cb5; [18–20]) and outer mitochondrial membrane proteins that regulate apoptosis (Bcl family; [21, 22]) or mitochondrial dynamics (*e.g.*, Fission 1/FIS1; [23]). TA proteins have been identified across evolutionary diverse organisms, including *Saccharomyces cerevisiae*, bacteria, *Homo sapiens*, *Arabidopsis thaliana*, and more recently, in the apicomplexan parasite *Toxoplasma gondii* [15, 24–27]. The TA biogenesis is well-characterized for proteins localized specifically to the ER and then transported from their ER-integrated state to the other cellular compartments (such as plasma membrane, nuclear envelope, Golgi complex, endosomes, lysosomes and peroxisomes) *via* the network of secretory vesicles (reviewed in [28, 29]). Multiple pathways are implicated in the targeting of TA proteins destined for the ER, including a promiscuous ‘moonlighting function’ by the SRP/Sec61 translocon (also involved in the co-translation translocation) [30–32], spontaneous insertion by the SRP-independent targeting (SND) components [31, 33, 34], and the components of the ER membrane complex (EMC) [35, 36]. Recently, it has also been suggested that TA proteins with more hydrophobic TMDs (compared to the mildly hydrophobic TMD of Cb5, which inserts in an unassisted manner) require the assistance of chaperones from the Hsp40/Hsc70 family [37, 38]. Thus, the targeting information for sorting of the nascent TA proteins is not defined by the presence of specific sequence motif within their TMD, rather it relies on physicochemical properties such as the length, hydrophobicity index and charge of the TMDs, as well as the length and the net charge of the CTS [39, 40]. For example, as compared to ER-localized TA proteins, mitochondrial-destined TA proteins tend to possess TMDs that are shorter and less hydrophobic, and their CTSs are enriched in positively charged amino acid residues [40].

In the yeast and mammalian cells, TA proteins destined for ER (or post-ER compartments) rely on the proteinaceous machinery known as the Guided Entry of TA (GET) or transmembrane domain recognition complex (TRC) pathway, respectively (reviewed in [41, 42]). However, intersectional competition between the SRP and the GET targeting pathways for the same RNCs and their convergence at the Sec61 translocon has also been observed [43–46]. In the GET pathway, the ER-destined TA proteins are first recognized by a small glutamine-rich tetratricopeptide repeat (TPR)-containing protein Sgt2 through the C-terminus hydrophobic binding domains and then transferred to ER-targeting cytosolic ATPase Get3, facilitated by the interactions with Get4/5 complex [47]. TA proteins destined to the mitochondria do not bind Sgt2 directly, instead they interact with other TPR-domain associated chaperones. Molecular chaperones of the Hsp70 family are also implicated to function upstream of Sgt2 in some studies [48, 49]. The TA-loaded Get3 subsequently interacts with the ER-resident Get1/2 transmembrane receptors that culminates in active membrane integration of the TA cargo in an ATP-dependent manner. In contrast, the TA insertion events in mammalian cells are more complex; in addition to homologues of Sgt2 (small glutamine-rich tetratricopeptide repeat-containing α; SGTA), Get4 (TRC35), Get5 (ubiquitin-like protein 4A; UBL4A), Get3 (TRC40) and Get1/2 (tryptophan rich basic protein/calcium-modulating cyclophilin ligand; WRB/CAML), higher eukaryotes also involve mediation by Bag6 (also known as BAT3/Scythe) in a complex together with TRC35 and UBL4A (constituting the Bag6 complex) that facilitates substrate transfer from SGTA to TRC40 [50]. Since Bag6 is primarily involved in the ubiquitination of any defective ribosomal proteins or mislocalized secretory and membrane proteins, its presence likely confers a protein quality control aspect to the TRC pathway [51, 52]. Interestingly, binding of substrate proteins to SGTA has been shown to revert Bag6-mediated ubiquitination and promote deubiquitination of mislocalized proteins, essentially shunting away TA clients for their delivery to the ER and not for their degradation [53–55].

In a recent study, 59 novel TA proteins were identified in a related apicomplexan parasite *Toxoplasma gondii* [27]. Domain swapping experiments further suggested an interplay between the sequence of the TMD and charge distribution in the CTS as deciding factors for the TA distribution across the ER, mitochondria, and Golgi apparatus. However, neither TA proteins nor the homologs of the GET/TRC pathway have ever been theoretically identified or experimentally validated in the human malaria parasite *Plasmodium falciparum*. *P. falciparum* causes the most severe form of human malaria [56]. During its asexual stages, the parasite invades and proliferates within human erythrocytes from early ring stages to mature trophozoites, and finally schizonts undergoing schizogony to form differentiated segmenters, which then liberate daughter merozoites in the blood stream after the erythrocytic rupture. Although the terminally differentiated red cells ensures a safe haven for the parasite away from the host immune system, they lack optimal infrastructure conducive for its nutrient and secretory needs. So, *P. falciparum* develops a complex network of membranes and vesicles for efficient protein trafficking [57]. These events not only help the parasite in its *in vivo* survival, they also ensure that the virulence determinants are trafficked specifically to their destined organelles. Protein trafficking in malaria-infected erythrocytes requires an additional level of refinement since the parasite resides within a parasitophorous vacuole (PV) of its host and can target proteins not only to intra-parasite organelles but further beyond; to the parasitophorous vacuolar membrane (PVM) and the infection-induced modified host erythrocytic structures (such as Maurer’s clefts, erythrocyte cytosol and red cell membrane knobs) [58], reviewed in [59]. The early secretory system in *P. falciparum* necessarily recruits the function of the SS similar to higher eukaryotes [60]. However, host-exported proteins require the presence of an additional leader sequence, known as *Plasmodium* export element (PEXEL) or host-targeting (HT) motif with a consensus RxLxE/D/Q (where x is any amino acid) [61, 62], and their export involves a multiprotein translocon complex [63]. Nonetheless, HT-independent trafficking has since been reported [64] and PEXEL-negative exported proteins (PNEPs) are found in the infected erythrocyte membranes lacking any discernible export motif [57]. In *P. falciparum*, the parasite ER functions as an obligatory destination for proteins during the initial stages of the secretory pathway. At the ER, important sorting decisions are made and exported proteins reportedly bind phosphatidylinositol 3-phosphate-enriched domains in the parasite ER through their HT/PEXEL motif [65] prior to their cleavage by the ER-resident aspartic protease plasmepsin V [66, 67]. However, the export pathway (via ER-recruitment) need not necessarily involve SS since PNEPs without an N-terminus signal peptide are efficiently recruited and exported to the Maurer’s clefts and the host erythrocyte [68].

Irrespective of the route of trafficking or the final locale within an infected erythrocyte, the roles of vesicular transport intermediates as carriers of virulence determinants are undisputed in *P. falciparum* [69]; reviewed in [70]. Model TA proteins, validated in other systems such as Syntaxins, SNAREs and NSF, are implicated in directing the budding and fusion of secretory vesicles in *P. falciparum*. Thus, there is an urgent need to identify the TA proteins and the underlying GET/TRC pathway in *P. falciparum*. In this study, we *firstly* identified a *P. falciparum* homolog of Get3 (PfGet3) based on the sequence, structure, and functional similarity to the Get3s of prokaryotic and eukaryotic origins. *Secondly*, we showed the association of PfGet3 with a subset of the total predicted 63 putative TA proteins in the *P. falciparum* proteome. Our proximity labelling experiments further revealed the identities of the two other homologs of the GET machinery in *P. falciparum*, such as PfGet4 and PfGet2, and we validated their association with PfGet3 by *in vitro* binding studies. We were, however, unable to definitively identify the plasmodial homologs of other GET components, such SGTA/Sgt2, Get5/UBL4A and Bag6 from our shortlisted proteins since all of them featured either the tetratricopeptide repeat motif (in the case of Sgt2/A homologs) or the characteristic ubiquitin-like domain (UBL; for Get5/UBL4A and Bag6), both of which are ubiquitously present across multiple proteins enriched from our proximity labelling studies and LC-MS/MS analyses. We were also unable to identify the plasmodial homolog of Get1/WRB due to extreme sequence divergence. Nonetheless, we believe this is the first study that provides bioinformatic and biochemical evidence for the presence of TA proteins and the partners of the GET machinery in the human malaria parasite *P. falciparum* and has the potential to lead the future investigations into elucidating the dynamics of this pathway and its development for therapeutic interventions.

## 2 RESULTS AND DISCUSSION

### 2.1 A repertoire of tail-anchored (TA) proteins in the *P. falciparum* proteome

To date, no annotation as designated ‘TA protein’ exist in the *P. falciparum* database. We thus used sequential selection criteria in the PlasmoDB (www.plasmodb.org) to identify putative TA proteins in the *P. falciparum* 3D7 strain, based on the following classical TA features: (i) absence of an ER-type secretory SS as predicted by SignalP (http://www.cbs.dtu.dk/services/SignalP/), (ii) presence of a single TMD within the first 50 amino acids from the C-terminal end of the protein, and (iii) orientation of the protein with respect to the TMD, *i.e.,* we favoured N-terminus_cytosolic_-TMD-C-terminus_luminal_ orientation (using the TMHMM server at http://www.cbs.dtu.dk/services/TMHMM/) (the schematic flow chart shown in **Figure 1A**). The output revealed a total of 130 proteins with these signatures (**Supplementary Table 1**), including homologs previously validated as TA proteins in other systems, such as Sec61β (PF3D7_0821800), guanine-nucleotide exchange factor Sec12 (PF3D7_1116400), SNARE proteins like Bos1 (PF3D7_1111300), VAMP7 (PF3D7_0826600), VAMP8 (PF3D7_1303200), mitochondrial Fission1 protein FIS1 (PF3D7_1325600) (**Table 1**). We manually removed the 67 misrepresented members from the list of outputs based on the results from earlier studies. These included 64 members of the repeated interspersed multigene family (RIFINs), mainly the subgroup RIFIN-A implying them as putative TA proteins. RIFINs constitute the largest known family of variant antigens in *P. falciparum* (∼150 copies per haploid 3D7 genome) and encode for 27-45-kDa proteins [71, 72]. Previous studies support the absence of a SS and the presence of a single TMD in RIFIN-As, while RIFIN-Bs are assumed to possess SS and two TMDs [73]. Members of RIFIN-A possess the host-targeting (HT) motif/*Plasmodium* export element (PEXEL) and are trafficked to the infected erythrocyte membrane via the parasite’s induced structures called Maurer’s clefts [74], thus they were excluded. Similarly, the other two parasite proteins, *i.e.,* PF3D7_0410000 (erythrocyte vesicle protein-1) and PF3D7_0936000 (ring-exported protein-2) are already confirmed to be exported to the host erythrocytes in previous studies [75, 76] and were eliminated since TA proteins are perceived to be confined within the cellular boundary. In addition, PF3D7_0206900 (merozoite surface protein 5), a GPI anchored protein [77], was also removed from the TA list (see **Supplementary Table 2** for the full list of eliminated proteins). Finally, a total of 63 TA proteins were predicted in the *P. falciparum* 3D7 (**Table 1**; the full list is in **Supplementary Table 3**).

**Figure 1.**
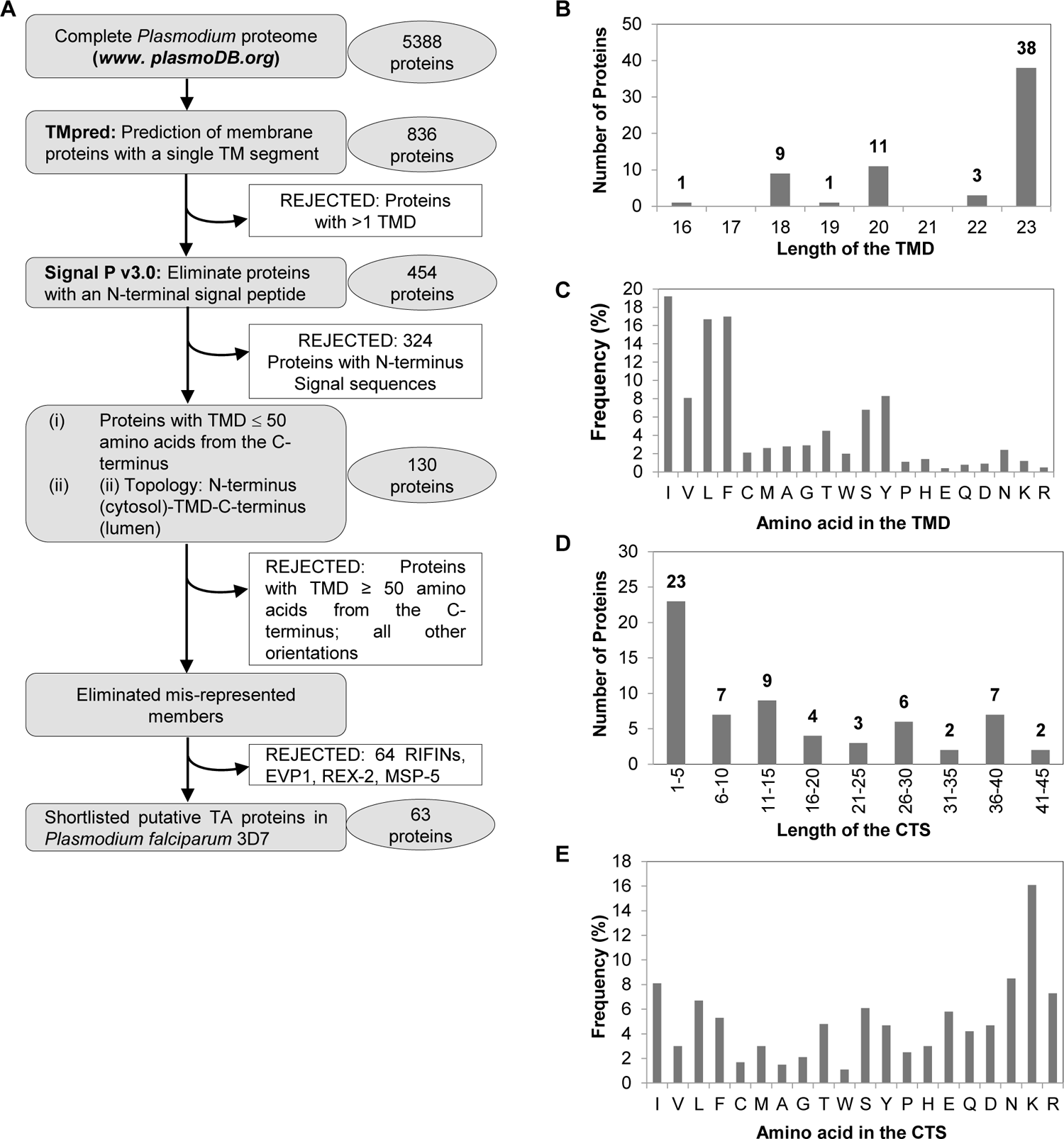
Identification of the putative Tail-Anchored (TA) proteins in *P. falciparum* 3D7. **A.** Flow diagram showing the successive bioinformatic steps performed, the corresponding retention/elimination criteria and the final outcome of the 63 predicted TA proteins in *P. falciparum* 3D7. **B-C.** Lengths of the predicted TMDs as according to TMHMM (www.cbs.dtu.dk/services/TMHMM) (B) and their amino acid compositions (C) for the 63 putative TA proteins. **D-E.** Lengths of the CTS (D) and the amino acid composition (E) were also calculated. The total number of predicted TA proteins are shown above each range of the TMD (B) and CTS (D) lengths. Amino acids listed (in C and E) are in decreasing order of hydrophobicity according to the Kyte and Doolittle scale [84].

**Table 1.**
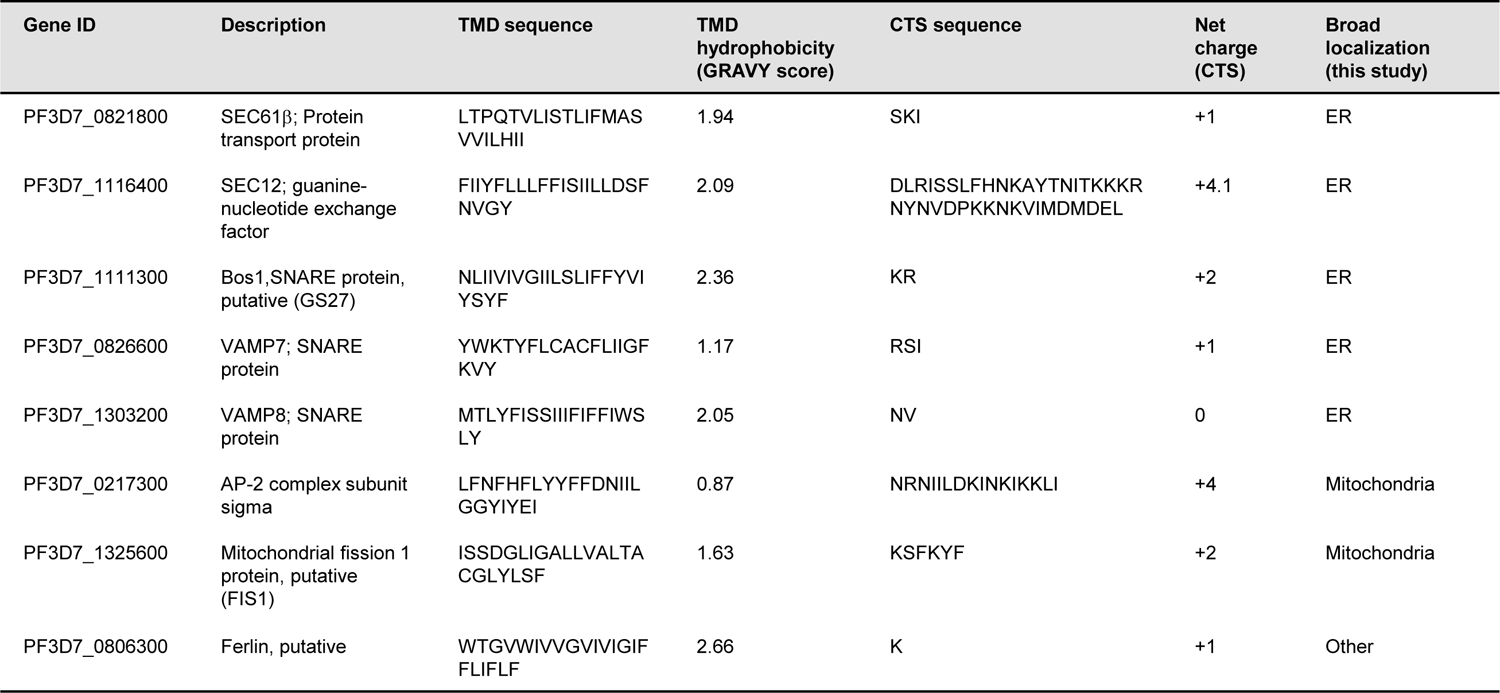
Representative list of the predicted TA proteins in P. falciparum 3D7 proteome. Proteins are listed according to their PlasmoDB ID (https://plasmodb.org/) and descriptions. Attributes of each protein include the transmembrane domain (TMD) sequence and its calculated hydrophobicity score (according to GRAVY calculator; http://www.gravy-calculator.de), the C-terminus sequence (CTS) and the net charge of the CTS. Broad organellar localization, as designated from this study, is also indicated. For the complete list, please refer *Supplementary Table 1*.

**Table 2.**
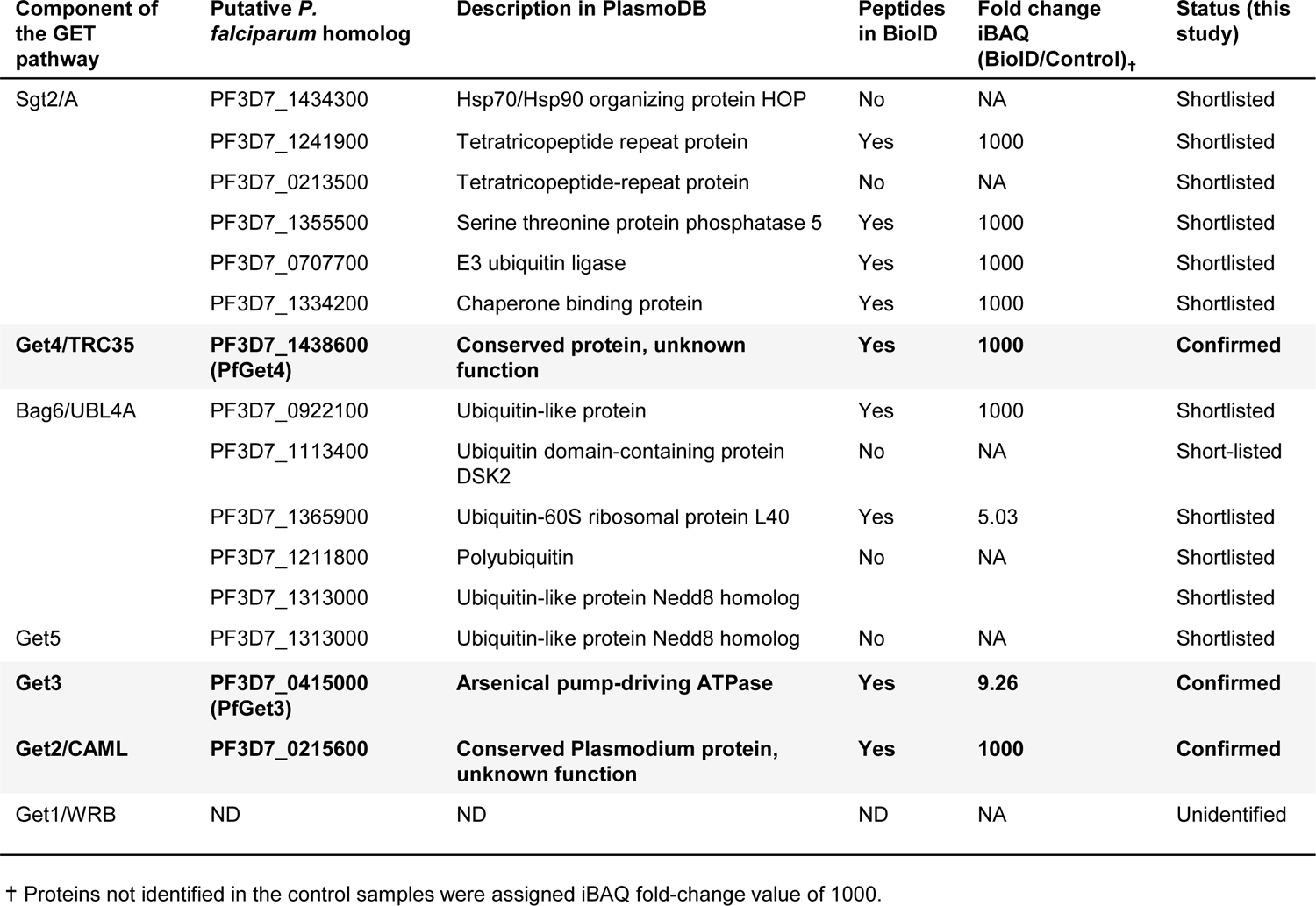
List of the confirmed and shortlisted homologs of the GET pathways in P. falciparum 3D7 identified in this study. Proteins are listed according to their PlasmoDB ID and descriptions. Rows showing the validated homologs of the GET machinery in *P. falciparum* are in bold and highlighted in grey, while shortlisted putative homologs without confirmed identities are in regular text. The fold change for each protein is measured as the ratio of iBAQ values between the BioID and control fractions (from *Supplementary Table 4*).

The TA proteins enter multiple cellular membranes including the endomembranous ER, Golgi, plasma membrane, mitochondria, and peroxisomes (and chloroplasts, in case of plants) (reviewed in [28, 29, 78]). The lengths of the TMDs and the amino acid composition of the CTS play important roles in the organellar localization of TA proteins across different species [19, 79–82]. Mitochondrial TA proteins often display significantly lower TMD hydrophobicity compared to the ER-destined pool [40]. A recent study carefully dissected the TMD and the CTS of multiple TA proteins in mammalian cells and implicated a delicate balance between the extent of TMD hydrophobicity and the net charge of the CTS in deciding the TA trafficking and/or their sharing between different organelles [83]. Peroxisomal TA proteins were shown to possess significantly higher net positively charged CTS as compared to those routed to the mitochondria, and the ER-destined TA proteins scored the least positively charged CTS. We thus investigated these features across the 63 putative TA proteins in *P. falciparum* 3D7. The TMD lengths varied between 18 to 23 amino acids among the TA proteins (TMHMM v.2.0; http://www.cbs.dtu.dk/services/TMHMM/), with an average length of 22 amino acids, correlating well within the helical length of 18-21 amino acids necessary to span the usual width of a lipid bilayer (**Figure 1B**). A bias towards the enrichment of amino acids with hydrophobic side chains was also observed across these TMDs (**Figure 1C**). In contrast, the length of the CTS was diverse, varying from 1-5 amino acid (23 putative TA proteins) to as high as 45 amino acids (in PF3D7_1422000) (**Figure 1D**). Also, there was no clear preference for any particular type of amino acids in the CTS (**Figure 1E**). We assigned the Grand Average of Hydropathy (GRAVY) (http://www.gravy-calculator.de) and Adagir (http://agadir.crg.es) scores to each TMDs of the 63 putative TA proteins in *P. falciparum* 3D7 (**Supplementary Table 3**) The GRAVY score assigns hydrophobicity indices to the TMDs based on Kyte and Doolittle [84], while the Adagir scores reflect the relative tendency to form helices in aqueous solution. In the GRAVY scale, the TMD hydrophobicity values ranged from the lowest 0.74 in PF3D7_1422000 to the highest 3.45 in PF3D7_1341700. PF3D7_1422000 is annotated as single pass putative cytochrome c oxidase assembly protein (COX14) that localizes to the mitochondria [85]. PF3D7_1341700, on the other hand, is a conserved plasmodial protein with unknown function. The other TA proteins with low hydrophobicity include PF3D7_0217300 (Adaptor protein complex-2 subunit σ; GRAVY score 0.87) playing an active role in vesicular membrane protein transport [86], and PF3D7_0306000 (putative cytochrome b-c1 complex subunit 8; GRAVY score 0.91) which is part of the mitochondrial electron transport chain driving oxidative phosphorylation [87]. Similarly, other annotated TA proteins with high hydrophobicity indices includes PF3D7_1332000 (Syntaxin 5; GRAVY score 3.04), PF3D7_0210700 (Syntaxin 17; GRAVY score 2.88) and PF3D7_0710800 (putative protein transport protein, USE1; GRAVY score 2.92) that constitute the SNARE complex and mediates anterograde or retrograde transport between ER and Golgi (reviewed in [88]). The ER-specific TA proteins identified in our list include PF3D7_0821800 (Protein transport protein Sec61 subunit β) and PF3D7_1116400 (Guanine nucleotide-exchange factor SEC12) and others, with an intermediate GRAVY score of 1.94 and 2.09, respectively. Similarly, lower Agadir scores are typically associated with TA proteins targeted to the mitochondria (mitochondrial fission 1 protein FIS1, PF3D7_1325600; Adagir score 1.39), while higher Agadir scores represent TA proteins targeted to the ER (syntaxin SYN11, PF3D7_1432000; Adagir score 10.52) [89].

We also manually grouped the 63 putative *P. falciparum* TA proteins into three broad sub-cellular localizations; namely the ER (and Golgi), the mitochondria and a collective of other destinations (including the endosomes/lysosomes, plasma membrane, peroxisomes, chloroplast etc.), based on the Gene Ontology (GO) annotations in the Uniprot database (http://uniprot.org) because, (i) there are no available machine learning tools to predict subcellular localization exclusively dedicated to apicomplexan parasites including *Plasmodium*, (ii) the three subcellular localization predicting algorithms [LOCTREE3 (https://rostlab.org/services/loctree3/), BUSCA (Bologna Unified Subcellular Component Annotator) (http://busca.biocomp.unibo.it/) and DeepLoc version 1.0 (http://www.cbs.dtu.dk/services/DeepLoc/)] portrayed notable discrepancies in their outputs for the plasmodial proteins (**Supplementary Table 3**), (iii) anaerobic protists and some parasites including *Plasmodium* display absence of peroxins and peroxisomes due to evolutionary reduction [90], thus ruling out this organelle. Further, (iv) *Plasmodium* parasites have unique organelles such as the apicoplast, food vacuole, rhoptries and micronemes which are likely to contain TA proteins but not represented in any algorithm. For example, apicoplast is reminiscent of chloroplast in plants, and chloroplasts contain essential TA proteins like SECE1 and SECE2 [78]. As a result, our analyses revealed a total of 14 ER-specific TA proteins, 7 mitochondrial-specific TAs and 42 TAs with diverse predicted cellular destinations (**Supplementary Table 3**). Thus, the collective features exhibited by the TMD and CTS of the 63 predicted TA protein in *P. falciparum* were consistent with previously confirmed TA proteins in other systems thereby validating our bioinformatic selection procedures (a few representative TA proteins are shown in **Table 1**).

### 2.2 Identification of a putative homolog of Get3 in *P. falciparum*

In both prokaryotes and eukaryotes, the conserved GET/TRC pathway has been implicated in targeting TA proteins through a concerted relay of events. Therein, the cytosolic ATPase Get3/TRC40 is located at the junctional interface and communicates with the pool of soluble components such as Sgt2/SGTA (hereafter referred to as Sgt2/A to incorporate features of both *H. sapiens* SGTA and *S. cerevisiae* Sgt2), Get4/5 or Bag6 at the upstream and the downstream ER membrane bound receptors Get1/2 (or WRB/CAML) [91–93]. We thus focussed our efforts towards identifying a corresponding homolog of Get3 in *P. falciparum* and subsequently use it as a handle to reveal the identities of the homologous effectors of the GET machinery and the bound TA substrates in this parasite. Although there are >2,000 putative Get3s reported in KEGG and OrthoDB databases [94], no Get3 homolog has yet been reported in the malaria parasite *P. falciparum.* Thus, we retrieved the amino acid sequences of the Get3 from yeasts (*Saccharomyces cerevisiae*; Uniprot ID Q12154, ScGet3; *Schizosaccharomyces pombe*, Uniprot ID Q9P7F8; SpGet3), plant (*Arabidopsis thaliana*; Uniprot ID Q12154; AtGet3) and mammalian (*Homo sapiens*; Uniprot ID O43681; HsGet3) systems from Uniprot (www.uniprot.org) and queried against the *P. falciparum* 3D7 database by BLAST in PlasmoDB. Results retrieved PF3D7_0415000 as the top hit in all queries. PF3D7_0415000 is a 379 amino acid protein annotated as a putative arsenical-pump driving ATPase, owing to its apparent ∼17% identity and 32% similarity to the bacterial arsenite transporter ArsA which provides resistance to arsenite [95, 96] (**Figure 2A**). Within eukaryotes, PF3D7_0415000 shared 46.9% similarity and 35.9% identity with ScGet3; 54.4% similarity and 41.8% identity with SpGet3; 51% similarity and 34.1% identity with AtGet3 and 59.7% similarity and 40.8% identity with HsGet3, respectively (**Supplementary Figure 1**). Like other Get3 members, PF3D7_0415000 also exhibited canonical features of NTPase superfamily, including the presence of (i) a nucleotide hydrolase domain (NHD) containing a A-type motif called as Walker B with a conserved ‘P-loop’ (GGKGGVGKTT) that directly interacts with the phosphate moiety of the NTP substrates, (ii) a conserved motif B, composed of a hydrophobic β-strand and terminating in an Asp residue (SVIVFD) which is involved in interaction with Mg^2+^ for catalysis, (iii) the A-loop for adenosine recognition (QLKNEIR) and (iv) Switch I (DPAHN) and Switch II (DTAPTGHT) loops that are reported to undergo conformational rearrangements in the presence of γ-phosphate of ATP to form a ‘loaded spring’ that uncoils upon ATP hydrolysis [95, 97, 98]. However, the ‘CxxC motif’ characteristic of the eukaryotic Get3s (ScGet3, SpGet3 and HsGet3/TRC40) which is implicated in coordination of the zinc ion for homodimerization was found to be absent in PF3D7_0415000 [99]. Instead PF3D7_0415000 displayed a deviant LxxC motif in which the first cysteine was replaced with leucine. Nonetheless, PF3D7_0415000 possessed a ∼20 residue insertion called the Get3 motif or TRC40 insert that is deemed essential for the substrate TA binding [99] (**Supplementary Figure 2**). Thus, these conserved features prompted us to confidently annotate PF3D7_0415000 as PfGet3 throughout the course of this study. PfGet3 also showed a high level of conservation across the other members of *Plasmodium* species, including those that infect humans (*P. vivax* and *P. knowlesi*), as well as the other rodent-specific malaria parasites (*P. yoelii, P. chabaudi* and *P. berghei*) (**Figure 2B** and **Supplementary Figure 1**). Phylogenetic analyses suggested that Get3 orthologs in the rodent malaria parasites diverge early in evolution and form a clade separate from the human counterparts. Get3 homologues have also been identified in other apicomplexan parasites, including *T. gondii* (TGVEG_231190), *T. cruzi* (Tc00.1047053507763.30), *L. donovani* (LDBPK_110710) and *C. parvum* (CGD7_4070) and they share varied similarity and identity to PfGet3 (**Supplementary Figure 1**).

**Figure 2.**
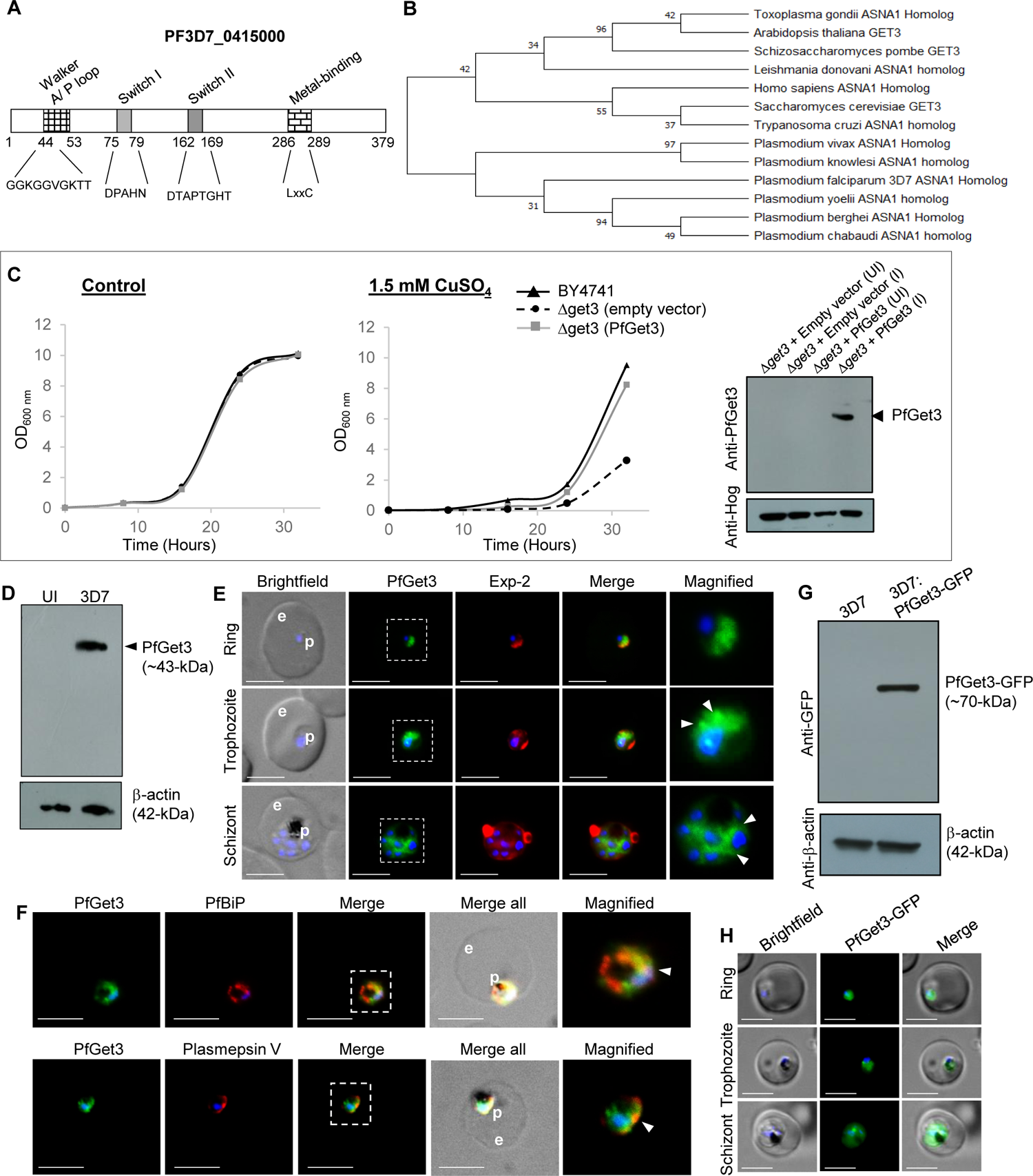
Identification and characterization of a *P. falciparum* homolog of Get3. **A.** Schematic representation of PFD_0415000 (PfGet3) showing the conserved motif characteristic of AAA-ATPase family such as the Walker B motif with conserved P loop, switch I and II motif for nucleotide binding, and the divalent cation binding motif. **B.** Phylogenetic reconstruction of putative Get3 homlogs in the plasmodial lineage thse corresponding to yeast (*S. cerevisiae* and *S. pombe*), related apicomplexan (*T. gondii*) and kinetoplastid parasites (*L. donovani* and *T. cruzi*), plant (*A. thaliana*) and human (*H. sapiens)*. The evolutionary history was inferred by using the Maximum Likelihood method [125] and branch annotation was done using bootstrapping. The percentage of replicate trees in which the associated taxa clustered together in the bootstrap test is shown next to the branches [148] and evolutionary analyses were conducted in MEGA X [149]. **C.** Growth curve of *S. cerevisiae* BY4741 wild-type (black lines) or *Δget3* cells transformed with either empty vector (dotted lines) or with sequences encoding for PfGet3 (grey lines) at 30°C in the absence (left) or presence (right) of 1.5 mM CuSO_4_. While no growth differences are apparent in the absence of CuSO_4_, the salt-sensitive growth-defect phenotype exhibited by *Δget3* cells transformed with the empty vector (dotted lines) is rescued by the expression of PfGet3. Western blot at the extreme right confirm the expression of PfGet3 (arrowhead) in transformed *Δget3* cells only under induced condition (I) as compared to uninduced (UI) control, using anti-PfGet3 antibodies (top). Anti-Hog antibodies detect *S. cerevisiae* Hog as a loading control (bottom). **D.** Western blot showing the expression of PfGet3 (arrowhead, top) at ∼43-kDa in infected erythrocytes (3D7, right lane) but not in uninfected (UI, left lane) red blood cells. Antibodies to the human β-actin (42-kDa) were used as a loading control (bottom). **E.** IFA images showing the expression and localization of PfGet3 (green) as diffused cytosolic fluorescence in ring (top panel), trophozoite (middle panel) and schizont (bottom panel) stages of infection. In trophozoites and schizonts, additional PfGet3-labeled structures as punctate spots or reticular pattens (white arrowheads) can be seen in the magnified versions of the dotted outlined regions. The parasite boundary is marked by antibodies to a parasitophorous membrane marker, PfExp-2 (red). Corresponding brightfield, single or merged fluorescent images with nuclear staining (blue) are shown. **F.** IFA images showing areas of colocalization between PfGet3 (green) and ER-resident proteins such as BiP (red, top panel) or Plasmepsin V (red, bottom panel) indicated by arrowheads in the magnified view of the dotted outlined regions. Corresponding brightfield, single or merged fluorescent images with nuclear staining (blue) are shown. **G.** Western blot showing the expression of PfGet3-GFP fusion protein (arrowhead, top) at ∼70-kDa in transgenic parasites (3D7:PfGet3-GFP, right lane) but not non-transfected controls (3D7, left lane) red blood cells. Antibodies to the human β-actin was used as a loading control (bottom). **H.** Live cell images showing the distribution of PfGet3-GFP (green) in transgenic parasites. Corresponding brightfield, fluorescence and merged images with the parasite nuclear staining (blue) are shown. For all live cell, brightfield and IFA images; p and e denote parasite and erythrocyte, respectively. Scale bar = 5 μm.

A null mutant in *S. cerevisiae* Get3 (*Δget3*) has been reported to show no growth defects in either synthetic complete media (or rich media) and exhibit no change in either cellular morphology, colony size, mating, sporulation, or germination [100]. Instead, *S. cerevisiae Δget3* strains are found to be more sensitive to As^3+^-, As^5+^-, Co^2+^-, Cr^3+^- and Cu^2+^-induced stress and high temperature-mediated stress than the parental wild-type strain [101]. A knockout of the Get3 homolog *asna1* in mice results in early embryonic lethality [102]. To investigate the comparable function between PfGet3 and ScGet3, we transfected *S. cerevisiae Δget3* strain with either empty vector or those containing sequences encoding for PfGet3 and assessed their growth at 30°C in the absence or presence of 1.5 mM CuSO_4_. The expression of PfGet3 at ∼43-kDa in transfected *Δget3* strain was confirmed by western blotting using custom-generated antibodies against PfGet3 (**Figure 2C**, right). Growth kinetics experiments revealed no apparent difference in growth between the wild-type BY4741 and *Δget3* strains in the absence of CuSO_4_. However, the *Δget3* strain transfected with the empty vector displayed a salt-sensitive growth phenotype that was successfully rescued by the *in trans* expression of PfGet3 to a level similar to the wild-type cells (**Figure 2C**). Thus, these results concluded that the functionality of PfGet3 is conserved across species.

### 2.3 PfGet3 shows cytosolic and ER distribution in the intraerythrocytic *P. falciparum* parasites

Transcriptomic data in PlasmoDB indicates two waves of *pfget3* transcriptions in *P. falciparum* 3D7 (refer to www.plasmodb.org for transcriptomic data): an initial high wave in ring stages within the first 10 hours post-invasion (hpi) and followed by a short shoulder at 20-36 hpi trophozoite stages. An abundance of *pfget3* transcripts occurs during the early ring stages with a peak at 5 hpi and followed by another peak with a smaller amplitude at 29 hpi trophozoites, thus suggesting the requirement of PfGet3 function in the early intraerythrocytic cycle during which the resident parasite actively modifies the host. To further investigate the expression of PfGet3 during the intraerythrocytic stages, polyclonal antibodies were custom generated against PfGet3 (see *materials and methods*) and validated in western blots with *P. falciparum* 3D7 parasites or with the purified recombinant PfGet3. Results indicated specific recognition at ∼43-kDa corresponding to the predicted molecular weight of PfGet3 as 43.3-kDa in *P. falciparum* parasites but not in uninfected erythrocytes (**Figure 2D**). Similarly, anti-PfGet3 antibodies also recognized the recombinant 6×his tagged PfGet3 expressed in *E. coli* cells (see **Supplementary Figure 3**).

Anti-PfGet3 antibodies were used in indirect immunofluorescence assay (IFA) to assess PfGet3 localization within the 3D7 parasites. IFA images predominantly showed a diffused PfGet3 fluorescence within the confines of the parasite boundary (and demarcated by antibodies to the parasitophorous vacuolar membrane protein Exp-2) and indicative of the parasite cytosolic staining of PfGet3 in the ring stage parasites (**Figure 2E**, top panel). However, in trophozoites and schizonts additional punctate/reticulate PfGet3 fluorescence were detected in the perinuclear region characteristic of the parasite ER compartments (**Figure 2E**, middle and bottom panels). These concentrated PfGet3 patterns could not be visualized in young rings owing to their small size and limitations in image resolution. In trophozoites and schizonts, the concentrated puncta of PfGet3 showed good colocalization with the ER chaperone PfBiP and the ER-resident parasite protease Plasmepsin V (**Figure 2F**), which suggested that a fraction of PfGet3 was indeed in the vicinity of the parasite ER and was consistent with the localization profile exhibited by the generic Get3 homologs in other systems. This further implied the existence of two different pools of PfGet3; a cytosolic soluble pool that possibly interacts with homologs other soluble cytosolic GET components (such as Sgt2/A, Get4 and Get5/UBL4A) for accessing the TA substrates; and an ER-membrane associated pool wherein the TA-bound Get3 interacts with the homologs of receptors like Get1/WRB and Get2/CAML for mediating TA-insertion into the ER. It is pertinent to mention that no homologs of the other GET components and neither the existence of a functional GET pathway has been identified in *P. falciparum* to date. Similarly, we also generated transgenic parasites expressing PfGet3-GFP, and western blots confirmed the expression of a fusion protein at ∼70-kDa (**Figure 2G**). Live cell microscopy of PfGet3-GFP transgenic parasites further showed the presence of the diffused cytosolic fluorescence as well as punctate Get3-GFP spots in the perinuclear region, thus confirming the distribution profiles seen in the IFA images (**Figure 2H**). The localization pattern of PfGet3 within infected erythrocytes strongly suggested the possibility of an active GET pathway in *P. falciparum*.

### 2.4 *In silico* and biochemical analyses of PfGet3 reveals architectures similar to the other Get3 homologs

Since there is no reported crystal structure of PfGet3, we submitted the amino acid sequence of PfGet3 to the Phyre2 server (www.sbg.bio.ic.ac.uk/phyre2) for *in silico* prediction of secondary structure and disordered region. Phyre2 output revealed the presence of 51% α-helices, 10% β-strands and 21% disordered regions (**Supplementary Figure 4A**; [103]). Submission of PfGet3 sequence to the TMHMM server (www.cbs.dtu.dk/services/TMHMM-2.0) predicted no transmembrane helix in PfGet3 (**Supplementary Figure 4B**) consistent with PfGet3 being a soluble cytosolic protein. The Phyre2 server also generated a 3D atomic model of PfGet3 based on remote homology and template-based assembly simulations to the existing Get3 crystal structures in the RCSB Protein Data Bank (PDB; **Figure 3A**). PfGet3 structure revealed 100% confidence with the various Get3 orthologs from both prokaryotic and eukaryotic origins (**Supplementary Figure 5**, showing the Phyre2 output). The PfGet3 3D structure showed highest identity (59%) to the Get3 from the yeast *Debaryomyces hansenii* (DhGet3; PDB ID: 3IO3; 1.8 Å resolution) followed by 51% identity to both nucleotide-free SpGet3 (PDB ID: 2WOO; 3.01 Å resolution) and ADP-AlF4 complexed ScGet3 (PDB ID: 2WOJ, resolved at 2.3 Å), respectively [99–104]. We preferred the PfGet3 model based on homology to the ScGet3 for our further analyses (**Figure 3A**). This is because: (i) the ADP-AlF4-ScGet3/2WOJ has 99.3% sequence coverage compared to 91.4% for DhGet3/3IO3 (the finger domain of DhGet3 was missing in the electron density map), (ii) PfGet3 rescued the CuSO_4_-sensitive phenotype in *S. cerevisiae Δget3* strain, and more importantly, (iii) the crystal structure of multi-subunit GET translocation complex (Get3-Get4-Get5) from *S. cerevisiae* was also available in the PDB at 6 Å resolution (PDB ID: 5BWK) [105], which we believe would aid in further characterizing the GET components and their interactions in *P. falciparum* in future studies. The distribution of hydrophobic and charged amino acids across PfGet3 surface also revealed the presence of a large hydrophobic patch, likely constituting part of the hydrophobic groove involved in binding the TMDs of TA substrates (**Figure 3B**). The residues forming the hydrophobic groove in PfGet3 were also found to be conserved among the other members of Get3 family ([104]; **Supplementary Figure 2**). We also submitted the PfGet3 sequence to the GalaxyWEB server (http://galaxy.seoklab.org) for the prediction of ligand and its binding sites [106]. Results indicated ADP as a possible ligand for PfGet3, similar to ScGet3 and the other Get3 homologs. Two regions of contacts via hydrogen bonds between PfGet3 and ADP were identified: the amino acid residues spanning the P-loop (47-53, GGVGKTT) and residues further downstream at the C-terminus (274Q, 347P, 349L, 353I, 354R and 362F) (**Supplementary Figure 6**). Interestingly, these residues of PfGet3 also show good conservation with the corresponding residues in ScGet3 thereby suggesting that the conserved functional features of Get3 extend across eukaryotes.

**Figure 3:**
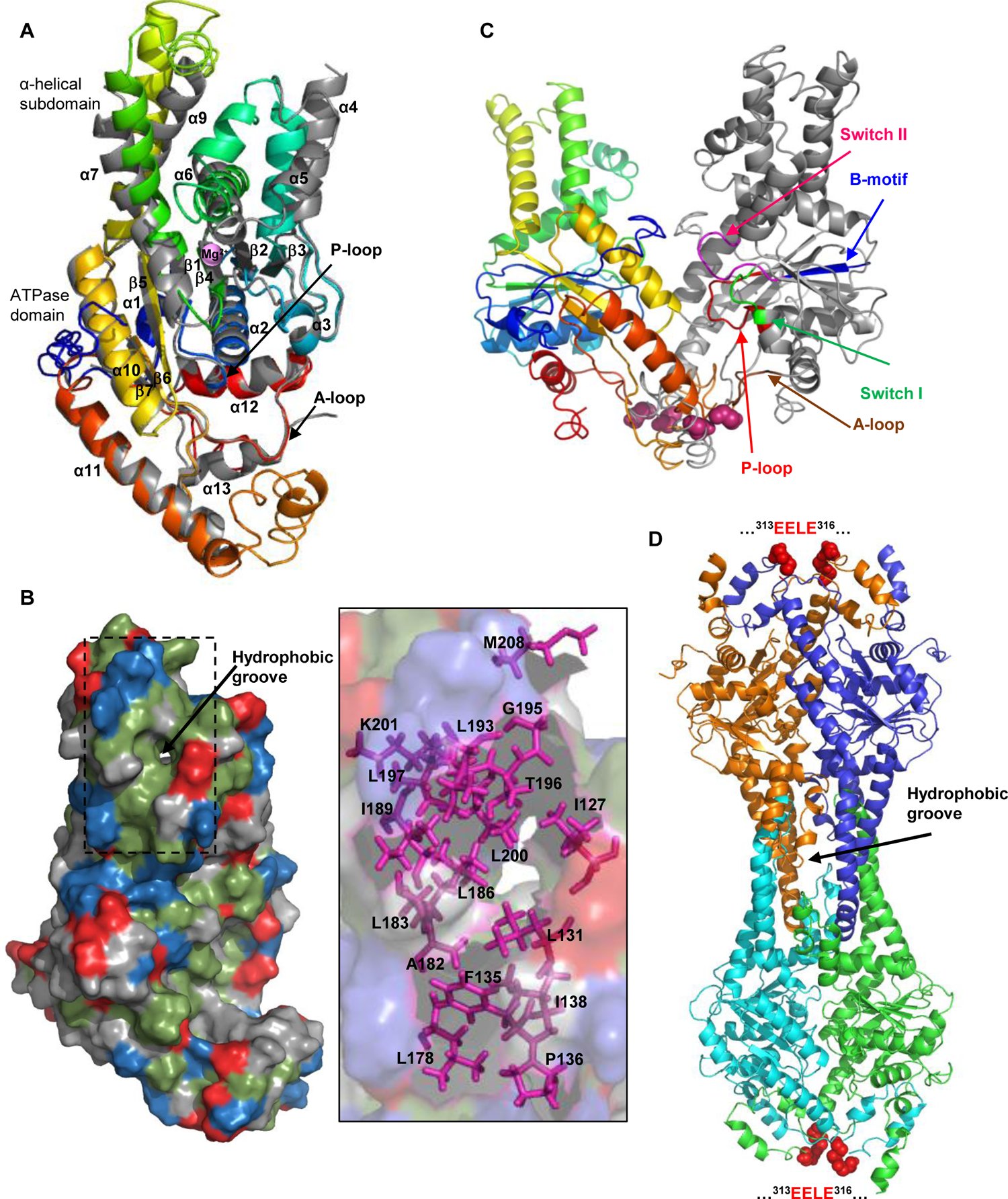
*In-silico* prediction of PfGet3 *s*tructural as compared to the other Get3 homologs. **A.** Alignment of the Phyre 2 (www.sbg.bio.ic.ac.uk/phyre2) predicted 3D structure of PfGet3 (rainbow colored) with the crystal structure of ScGet3 (PDB ID 2WOJ; grey) [99]. The α helices and β-sheets are numbered, and the core ATPase domain and α helical subdomains are labeled. The positions of Mg^2+^ is shown (pink sphere). The P- and A-loops are indicated by arrows. **B.** Surface representation of PfGet3 showing the putative TMD-binding hydrophobic groove (black arrow). The hydrophobic, acidic and basic amino acids are colored as green, red and blue, respectively. Magnified image of the hydrophobic pocket (dotted outline) is shown at the right in 50% surface transparency with conserved residues involved in forming the TMD-binding groove indicated by magenta sticks [104]. The orientation is similar to that in A. **C.** Predicted PfGet3 dimer. One monomer is shown as rainbow colored and the other is colored grey. P-loop (red), A-loop (brown), B-motif (blue), Switch I motif (green) and Switch II motif (magenta) are as indicated. The ELLE (ExxE) motif of each chain at the dimer interface are shown as pink spheres. **D.** Predicted PfGet3 tetramer based on MjGet3 (PDB ID 3UG7) template and showing the hydrophobic pocket (black arrow). The four subunits of PfGet4 are colored as blue, orange, cyan and green. The EELE motif of each chain at the dimer interface is shown as red spheres. All structural representations shown were generated using PyMOL software (www.pymol.org).

The deduced X-ray crystal structures of various Get3 homologs indicate diverse oligomerization assemblies. While ScGet3, SpGet3 and DhGet3 exhibits association of two monomers into a rotationally symmetrical homodimer analogous to the arrangement of tandem domains seen in ArsA, the archaeon *Metanocaldococcus jannaschii* Get3 (MjGet3), the human ortholog TRC40 and ScGet3 also display tetrameric assemblies under specific conditions [94, 95, 99, 104, 107–109]. Yet higher complex oligomerization assemblies were known to exist for some Get3 homologues, *for example,* the ADP-bound Get3 from thermophilic opportunistic human pathogen *Aspergillus fumigatus* (AfGet3; PDB ID code 31BG) has been reported to form a hexamer in the asymmetric unit with 3-fold symmetry of each monomer assembling into a dimer [95]. The apo-form of AfGet3 associates as a dimer. Similarly, trimeric forms of ArsA have also been visualized by electron microscopy and chromatographic studies [110]. The 3D structure of PfGet3 was thus submitted to GalaxyWEB server for homomer prediction [111, 112]. The top 4 out of 5 outputs suggested dimeric association (with the 2WOO/SpGet3, 3IQW/*Chaetomium thermophilum* Get3, 3IBG/ADP-AfGet3 or 3SJA/ScGet3 as templates, respectively), while the 5^th^ output predicted a tetrameric association (with the 3UG7/MjGet3 as template) (**Supplementary Figure 7A**). In the absence of any resolved crystal structure, we preferred an unbiased representation for our *in silico*-based oligomerization propensity of PfGet3. Accordingly, both the 3SJA template-based dimer (representing the open state crystal structure of ScGet3 in complex with Get1 cytosolic domain) and 3UG7/MjGet3-template based tetramer were selected to represent the possible oligomerization status of PfGet3 (**Figures 3C** and **3D**). Prior reports have also indicated that the formation of homodimer by most of the Get3 members is a prerequisite for both its ATPase activity and the formation of TMD-binding hydrophobic groove. In fungi and mammals, this is brought about by the CxxC motif in each subunit which aligns to coordinate a zinc ion [99, 113]. Surprisingly, PfGet3 lacks the corresponding CxxC motif and instead show a deviant LxxC motif by sequence alignment (**Supplementary Figure 2**). However, downstream sequences in PfGet3 reveal the presence of an ExxE motif in the vicinity of the dimerization interface (**Figures 3C** and **3D**). The ExxE motifs are known to coordinate iron ions and are characteristic of certain land plant homologs that are devoid of the CxxC motif [94, 114, 115]. Thus, the ExxE motif may contribute towards the homodimerization of PfGet3, which may further assemble into higher oligomers (**Figure 3C-D** and **Supplementary Figure 7B-C**).

In summary, our *in-silico* comparison highlighting the structural and ligand-binding similarities between PfGet3 to other Get3 homologs, in conjunction with its complementation capability in rescuing the CuSO_4_-sensitive phenotype of *S. cerevisiae Δget3* strain, strongly suggested the presence of a functional GET machinery in *P. falciparum*.

### 2.5 Proximity labelling BioID approach indicates association between PfGet3 and multiple TA-proteins

BioID harnesses the potential of biotin ligase BirA^*^ (BirA with a R118G mutation) to promote promiscuous biotinylation. When fused to a ‘bait’ protein, the ligase catalyses a two-step reaction in the presence of biotin to generate reactive bioAMP that covalently biotinylates adjacent primary amines of the proximate ‘prey’ proteins [116]. These biotinylation events are then effectively enriched using streptavidin affinity matrices and permit the identification of dynamic or weak interactions largely lost during the complicated (immuno)-purification protocols (see schematic in **Figure 4A**). Since the interactions of TA proteins with Get3 are deemed transient; thus, to experimentally validate our bioinformatic predictions of TA proteins in *P. falciparum* 3D7 and their association with PfGet3, we generated transgenic parasites expressing PfGet3 fused to BirA*-HA at the C-terminus (see *materials and methods*). Expression was confirmed in western blots using antibodies to the HA tag, which detected PfGet3-BirA*-HA fusion protein of 76-kDa (**Figure 4B**). IFA of the PfGet3-BirA*-HA-expressing parasites showed distribution within the parasite cytosol as well as in some concentrated puncta, a profile similar to that of the native PfGet3 and suggesting no aberrant effect of BirA^*^ tagging on PfGet3 localization (**Figure 4C**). Equal volumes of the total extracts from PfGet3-BirA*-HA cells grown either in the absence (control fraction) or in presence of D-Biotin (named as BioID fraction, hereafter) (**Figure 4D**) were then used to affinity purify the proximal biotinylated interactors using StrepTactin XT beads and identified either by western blotting or LC-MS/MS. In western blots using antibodies to both HA tag and PfGet3, the PfGet3-BirA*-HA was enriched only in the BioID fraction (**Figure 4E**) although it could be detected in the total lysates of both the control and BioID fractions (**Figure 4D**). Interestingly, antibodies to PfGet3 also detected the native untagged PfGet3 on longer exposure of the membrane thereby implying the formation of a ‘heteromer’ (and possibly dimer, tetramer or higher oligomerization states) between untagged PfGet3 and PfGet3-BirA*-HA (as expected from our *in-silico* oligomerization prediction, **Figure 3C**) and the thus the ensuing biotinylation. Self-biotinylated PfGet3-BirA*-HA was also detected in western blots. Mass spectrometry results yielded a list of proteins enriched in the BioID fraction of PfGet3-BirA*-HA as compared to the control sample (see **Supplementary Table 4** for the complete list). PfGet3 was one of the top hits with ∼10-fold excess and 24 different peptides, which may have been sourced either from self-biotinylated PfGet3-BirA*-HA or *trans*-biotinylated native PfGet3 (**Figure 4E**). These also included two biotinylated peptides GTLNVL**K**NFTNNEMEFDSLYEK and **K**EMFDNILPELLHSFPGIDEALCFAELMQSIK (**K** indicates the site of biotinylation), thus confirming successful recruitment of the biotinylizer.

**Figure 4.**
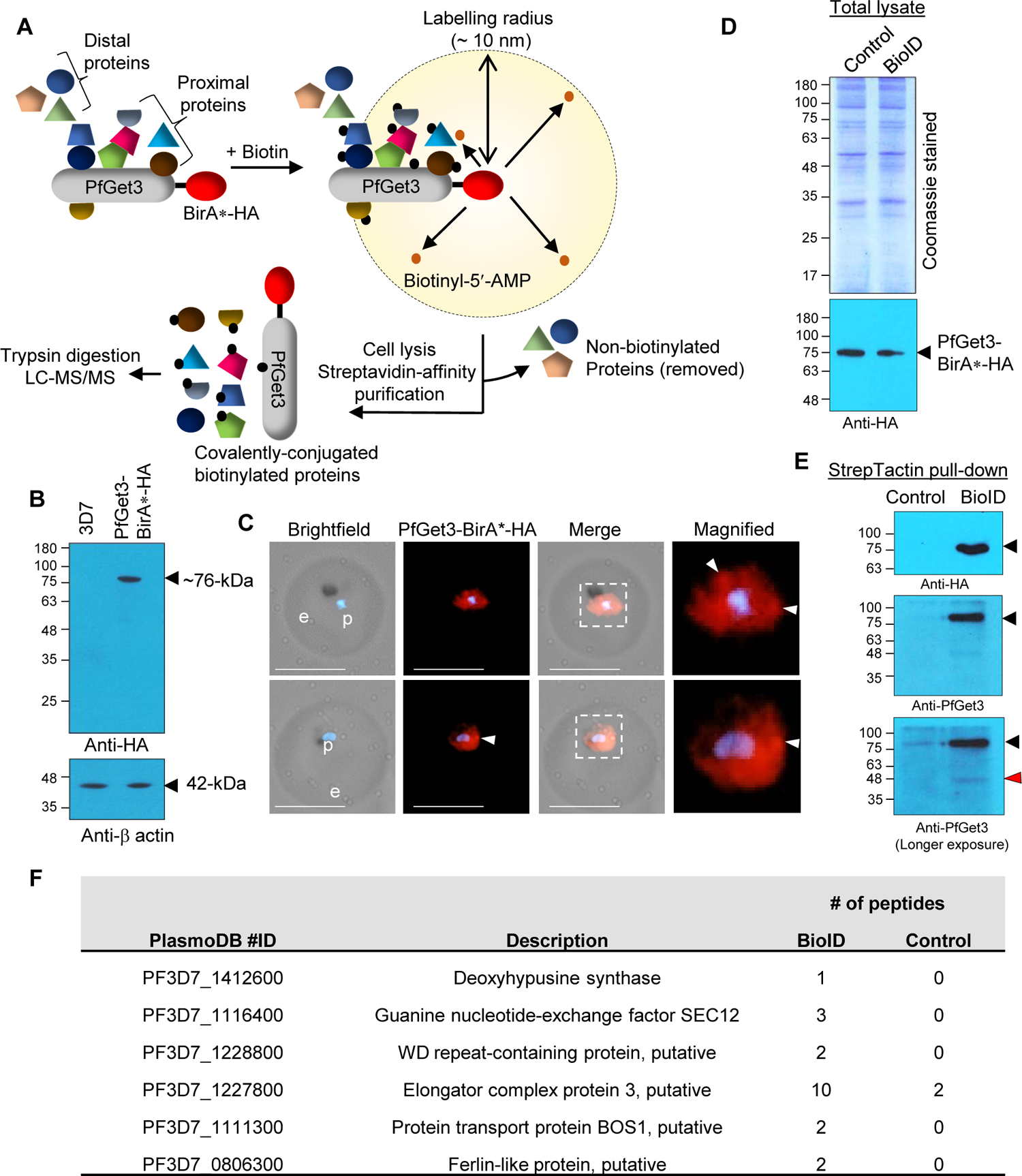
Proximity biotinylation using PfGet3-BirA*-HA and association with TA proteins. **A.** Schematic of the proximity biotinylation by the bacterial biotin ligase BirA^*^ (red) fused to the C-terminus of PfGet3 (with a HA tag downstream). BirA^*^ converts exogenously added free biotin to highly reactive biotinyl-5-AMP (small orange dots) which reacts with the primary amines of proximal proteins (within 10 nm radius, interacting directly or indirectly) and cause their biotinylation (indicated by small solid black dots). Distal proteins beyond the 10 nm radius are not biotinylated irrespective of their interactions with PfGet3-BirA*-HA. Following biotinylation, the biotinylated proteins are affinity purified from cellular lysates using streptavidin beads and identified by LC-MS/MS. Adapted from [157]. **B.** Western blot showing the expression of ∼76-kDa PfGet3-BirA*-HA (arrowhead, top) only in transgenic parasites (right lane) but not in the non-transfected control (3D7, left lane). Antibodies to β-actin at ∼42-kDa (bottom) are used as loading control. **C.** IFA images showing the distribution of PfGet3-BirA*-HA (red) in transgenic parasites. Corresponding brightfield and fluorescent images are as indicated. Parasite nucleus (blue) is stained with Hoechst 33342. Scale bar = 5 μm. **D.** Coomassie stained SDS-PAGE (top) and western blot (bottom) of the total lysates prepared from transgenic parasites expressing PfGet3-BirA*-HA grown in the absence (control fraction) or presence (BioID fraction) of D-biotin. The Coomassie-stained gel indicates equal loading and the western blot using antibodies to the HA tag depicts equal intensity of PfGet3-BirA*-HA expression under both conditions. **E.** Western blots showing the presence of PfGet3-BirA*-HA protein (∼76-kDa; black arrowheads) only in the StrepTactin affinity-purified BioID fraction and not in the control fraction, using antibodies to the HA-tag (top) or PfGet3 (middle and bottom). The blot at the bottom represents longer exposure of the blot in the middle to highlight the detection of native PfGet3 (red arrowhead), possibly due to homomer formation with PfGet3-BirA*-HA. In all SDS-PAGE and western blots, molecular weight standards (in kDa) are as indicated. **F.** Tabular representation of the 6 predicted TA proteins enriched in the BioID sample as compared to the control fraction. The corresponding PlasmoDB ID and descriptions are mentioned along with the relative number of peptides detected by LC-MS/MS (see *Supplementary Table 4* for the complete list).

Among the predicted TA proteins in *P. falciparum*, peptides corresponding to six proteins were quantitatively enriched in the BioID fraction (**Figure 4F**). These include peptides from PF3D7_0806300 (Ferlin-like protein, putative), PF3D7_1111300 (Protein transport protein BOS1, putative), PF3D7_1116400 (Guanine nucleotide-exchange factor SEC12), PF3D7_1227800 (Elongator complex protein 3, putative), PF3D7_1228800 (WD repeat-containing protein, putative) and PF3D7_1412600 (Deoxyhypusine synthase), which accounted for approximately 43% and 10% of the 14 ER-predicted and the total 63 predicted TA proteome, respectively in *P. falciparum*. We also observed that the sum of peptides derived from each TA protein was limiting and we were unable to find biotinylation in any of the peptides corresponding to these proteins. This could either be due to their low abundance or transient interaction with PfGet3-BirA*-HA that limits the scope of biotinylation at all the possible sites within these TA proteins. Nonetheless, we strongly believe that these TA proteins were biotinylated since they were affinity-purified with the StrepTactin-resin and enriched in the BioID fraction as compared to the control. Among the enriched TA proteins, PF3D7_1111300 and PF3D7_1116400 have been classified as ER-localized TA family in this study and corresponding homologs already validated in other systems [117, 118]. Although the remaining hits; PF3D7_0806300, PF3D7_1227800, PF3D7_1228800 and PF3D7_1412600 belong to TA proteins localized to other destinations (see **Supplementary Table 3**), it is possible that their targeting in *P. falciparum* intersects with the GET pathway involving PfGet3. Interestingly, our BioID fraction was devoid of any mitochondrial TA proteins, an expected result since mitochondrial TA targeting essentially recruits GET/TRC-independent mechanisms [12, 33, 119, 120]. Similarly, even within the subset of ER-resident TA proteins, other trafficking mechanisms are known to exist that preferentially utilize the SRP or the Hsc70-Hsp40 systems instead of the GET/TRC route [30, 31, 121, 122], thus explaining the low numbers of TA proteins detected in our BioID fraction. Recently, a study also revealed the inability of TRC40 (the mammalian homolog of PfGet3) to effectively engage TA proteins with moderately hydrophobic TMD. These proteins are instead shielded off from the hydrophilic cytosol by calmodulin (and not by Sgt2/A) and inserted at the ER via the conserved ER membrane protein complex (EMC) rather than the GET pathway [35]. Within the ER-specific TAs grouped in this study, we found at least six proteins (PF3D7_0323100, PF3D7_0618700, PF3D7_0627300, PF3D7_0821800, PF3D7_0826600 and PF3D7_1032400) exhibit comparatively lower GRAVY scores (<2.0) with respect to the others (with GRAVY scores >2.0), which might have contributed towards the absence of their biotinylation and ‘capturing’ in our PfGet3 BioID fraction. Taking this into account, our BioID fraction essentially detected 2/8 (or 25%) of the TA proteins that were most plausible to recruit the GET pathway for ER trafficking. However, the precise GRAVY score that decides preference for either GET (in)dependent pathway is currently unknown in *P. falciparum* and further experiments are warranted to ascertain their detailed characterization. Nonetheless, our results validate the existence of TA proteins in *P. falciparum* and their association with PfGet3 via a functional GET pathway.

### 2.6 PfGet3 also interacts with a putative homolog of Get4 in *P. falciparum*

Our BioID fraction also enriched 8 peptides from PF3D7_1438600 while only one was detected in the control (**Supplementary Table 4**). Interestingly, PF3D7_1438600 was also the only significant hit in BLAST searches with the amino acid sequences of the well-characterized Yor164c (*S. cerevisiae* homolog of Get4) and the *H. sapiens* Golgi to ER traffic protein 4 homolog/TRC35 (HsGet4) against *P. falciparum* 3D7 proteome, with significant e-values of 2e-4 and 9e-11, respectively. PF3D7_1438600 encodes for a 277 amino acid cytosolic protein, without any ER-type secretory signal sequence or TMD and has been annotated as a conserved protein with a domain of unknown function (DUF410) in PlasmoDB. PF3D7_1438600 shares considerable similarity with Yor164c (17.6% identity and 36.3% similarity), HsGet4/TRC35 (15.6% identity and 37.8% similarity) as well as other Get4 homologs (**Supplementary Figure 8**), including the residues involved in the interactions with Get3 (**Supplementary Figure 9**; [123, 124]). Homologs of Get4 form a hetero-tetrameric complex with Get5 and mediates efficient delivery of TA substrates from Sgt2 to Get3 (schematic in **Figure 5A**; adapted from [49]). Evolutionary analyses of the orthologs of Get4 by Maximum-likelihood method [125] revealed PF3D7_1438600 in a clade with other *Plasmodium* parasite species that infect humans (**Figure 5B**). Surprisingly, the Get4 ortholog of the rodent malaria parasite *P. yoelii* (PY00666) also grouped closer to the human malaria parasites (*P. falciparum* and *P. knowlesi*) than other rodent malaria species (*P. berghei* and *P. chabaudi*). We thus annotated PF3D7_1438600 as PfGet4 hereafter in this study. Prior structural studies have shown that yeast Get4 is an alpha-helical repeat protein with the N-terminus (residues 1-148) involved in Get3 binding and its C-terminus interacting with Get5 [123, 124]. Since no structural information about PfGet4 is available, we submitted PfGet4 amino acid sequence to the Phyre2 server (www.sbg.bio.ic.ac.uk/phyre2) for *in silico* secondary structure prediction. Phyre2 output revealed the presence of 72% α-helices and 14% disordered regions (**Supplementary Figure 10**; [103]). PfGet4 structure showed 100% confidence with PDB structures 6AU8, 3LPZ and 2WPV, which are crystal structures of the human TRC35 in complex with Bag6-NLS, C. *thermophilum* Get4 and S. *cerevisiae* Get4-Get5 complex, respectively (**Supplementary Figures 11** and **12**). Essentially, PfGet4 formed 14 right-handed helices arranged pairwise with an α -solenoid similar to the TRC35 and ScGet4 (**Figure 5C**; [123, 124]). In addition, PfGet4 successfully rescued the CuSO_4_-sensitive growth phenotype in *Δget4* strain to a similar level as the wild-type cells as compared to *Δget4* strain transfected with vector only (**Figure 5D**). Custom anti-peptide antibodies against PfGet4 were generated (described in the *materials and methods*) and detected a parasite protein at ∼33-kDa (predicted molecular weight 33,150 Da) in infected erythrocytes but not in uninfected RBC (**Figure 5E**). IFA revealed localization of PfGet4 as diffused fluorescence indicative of the parasite cytosolic staining. Colocalization studies in transgenic parasites expressing PfGet3-BirA*-HA using mouse-anti-HA antibodies in conjunction with anti-PfGet4 (both our anti-PfGet3 and anti-PfGet4 antibodies were generated in rabbits, so we were unable to use them in parallel) correlated well with both PfGet3 and PfGet4 showing characteristic cytosolic staining profiles (**Figure 5F**). To assess the interaction between PfGet4 and PfGet3 *in vitro*, we expressed and purified recombinant MBP-PfGet4 in *E. coli* (see *materials and methods*). Purified recombinant MBP-PfGet4 or recombinant MBP (as a control) were incubated with recombinant PfGet3-6×his and affinity-purified using Ni-NTA beads to assess co-association. Western blots using anti-MBP antibodies confirmed the selective binding of MBP-PfGet4 (but not MBP) to PfGet3 (**Figure 5G**). Based on these observations and our earlier BioID results, we reasonably concluded that the *P. falciparum* genome encodes for a functional PfGet4 exhibiting features similar to other Get4 homologs, and PfGet4 directly interacts with PfGet3, as expected for the canonical GET pathway.

**Figure 5.**
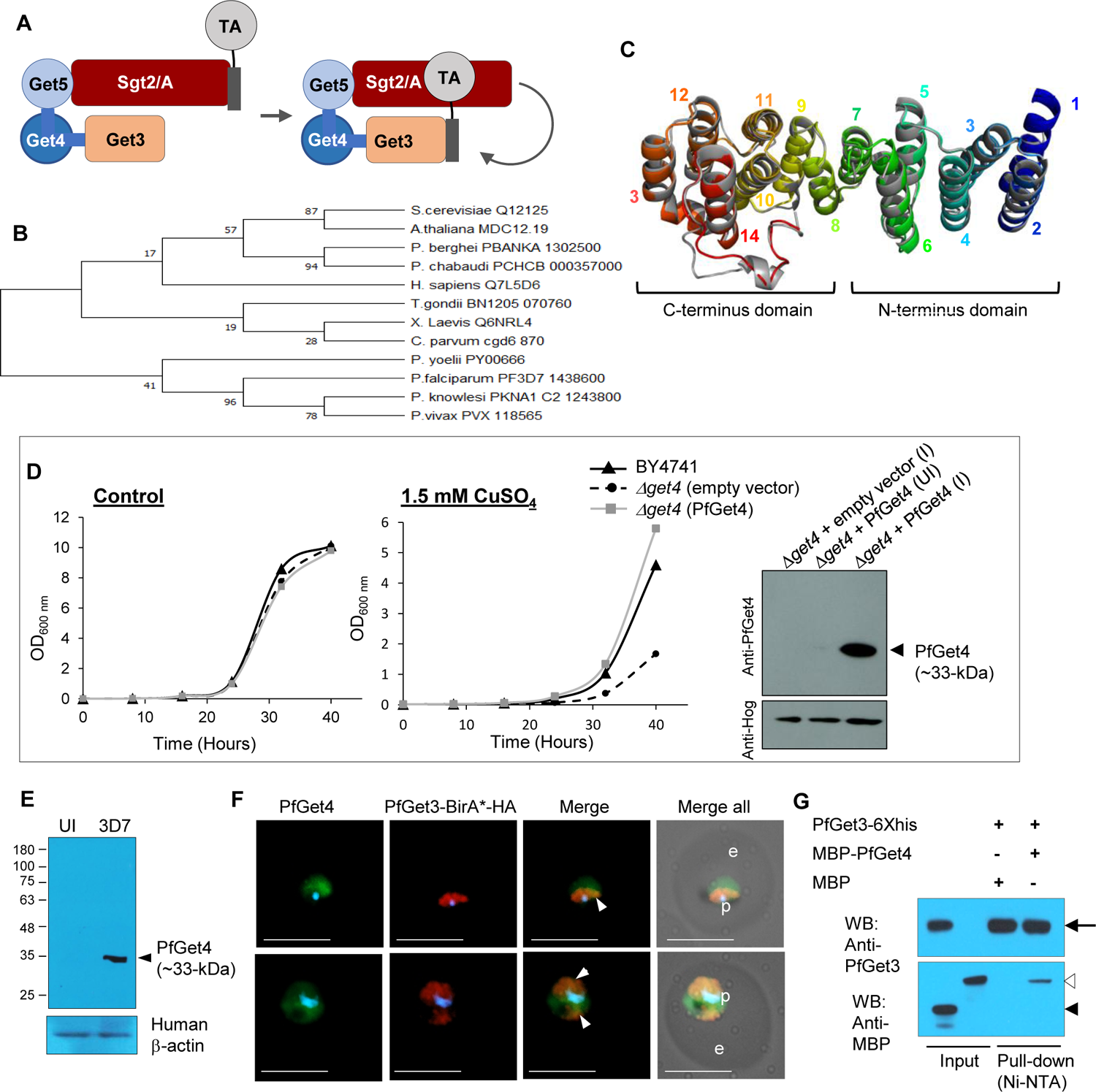
Identification of PfGet4 and its association with PfGet3. **A.** Schematic showing the direct interactions between Get3 and Get4. Get5 and Sgt2/A, however, indirectly interact with Get3 via the Get4. The Sgt2/A-associated TA protein is subsequently transferred to Get3. **B.** Phylogenetic reconstruction of Get4 proteins in the plasmodial lineage with homologs from yeast (*S. cerevisiae*), related apicomplexans (*T. gondii* and *C. parvum*), *X. laevis*, plant (*A. thaliana*) and human (*H. sapiens)*. The evolutionary history was inferred by using the Maximum Likelihood method [125] and branch annotation was done using bootstrapping. The percentage of replicate trees in which the associated taxa clustered together in the bootstrap test are shown next to the branches [148] and evolutionary analyses were conducted in MEGA X [149]. **C.** Phyre2 (www.sbg.bio.ic.ac.uk/phyre2) predicted 3D structure of PfGet4 (rainbow colored) aligned with the crystal structure *H. sapiens* TRC35 (PDB ID 6AU8A/grey) [123]. The α-helices are numbered and the N- and C-terminal domains are as indicated. **D.** Growth curve of *S. cerevisiae* BY4741 wild-type (black lines) or *Δget4* cells transformed with either empty vector (dotted lines) or with sequences encoding for PfGet4 (grey lines) grown at 30°C in the absence (left) or the presence of 1.5 mM CuSO_4_ (right). While no growth differences are apparent in the absence of CuSO_4_, the salt-sensitive growth defect exhibited by *Δget4* cells transformed with the empty vector (dotted lines) is rescued by the expression of PfGet4. Western blot at the extreme right confirm the expression of PfGet4 (arrowhead) in transformed *Δget4* cells only under induced condition (I) as compared to uninduced (UI) control, using custom generated anti-PfGet4 antibodies (top). Anti-Hog antibodies detect *S. cerevisiae* Hog as loading control (bottom). **E.** Western blot showing the expression of PfGet4 (arrowhead, top) at ∼33-kDa in infected erythrocytes (3D7, right lane) but not in uninfected (UI, left lane) red blood cells. Antibodies to the human β-actin were used as loading control to detect β-actin at 42-kDa (bottom). Molecular weight standards (in kDa) are as indicated. **F.** IFA images showing the areas of colocalization between PfGet4 (green) and PfGet3-BirA*-HA (red) indicated by arrowheads in the merged view of fluorescence. Corresponding brightfield, single or merged fluorescence images with nuclear staining (blue) are shown. Scale bar = 5 μm; e, erythrocyte; p, parasite. **G.** Western blots showing the association of purified 6×his tagged recombinant PfGet3 (∼43-kDa; black arrow) with MBP-PfGet4 (empty arrowhead) but not with MBP (empty position indicated by solid arrowhead) as revealed by pull-down assay using Ni-NTA beads followed by western blot using antibodies to MBP (bottom) or PfGet3 (top). Corresponding input and pull-down lanes are as indicated. Molecular weight standards (in kDa) are as shown for all SDS-PAGE and western blot images.

### 2.7 Other homologs of *P. falciparum* GET machinery are also enriched in the BioID fraction

Get3 is a central component of the GET pathway, orchestrating connections between the TA-recognition complex (as it captures the TA substrates from the emerging ribosomal tunnel) and the downstream Get1/2 receptors at the ER membrane that functions to insert the TA proteins. Thus, the transgenic PfGet3-BirA*-HA parasites also offered possibilities to biotinylate both upstream and downstream constituents of the GET pathway. We therefore mined the BioID fraction further to identify the other GET machinery homologs in *P. falciparum*. The TRC comprises of the yeast cochaperone Sgt2 (or SGTA; the equivalent mammalian homolog) and scaffolding proteins Get4 (or TRC35, the mammalian homolog) and Get5 (or mammalian Bag6 complex) [12]. The binding of the TA proteins to Sgt2/A is considered as the first committed step for the GET/TRC trafficking pathway (**Figure 5A**; [49, 93, 126]). However, the role of cytosolic chaperones like the yeast cytosolic Hsp70, Ssa1 and Hsp40 in harnessing the energy of ATP hydrolysis to unfold aggregated TA substrates and preserve their conformational quality prior to Sgt2/A loading has also been reported [48]. The central region of Sgt2/A contains the conserved TPR domains that associates with additional heat-shock proteins like Hsp70, Hsp90 and Hsp104 [49, 127, 128]. Heat-shock cognate 70 (Hsc70) has been found to be associated with *in vitro* translated TA proteins from mammalian lysates and mutations in residues of the TPR domain which prevent Hsp70 binding impair the loading of TA substrates onto Sgt2/A [31, 48]. We observed a selective enrichment of heat-shock proteins like PfHsp90 (PF3D7_0708400), multiple PfHsp70s (PF3D7_0818900, PF3D7_0917900/BiP, PF3D7_1134000), PfHsp110 (PF3D7_0708800), PfHsp60 (PF3D7_1015600), PfHsp101 (PF3D7_1116800) and small heat shock protein (PF3D7_1304500) in our BioID fraction (**Supplementary Table 4**). Neither PfGet3 nor PfGet3-BirA*-HA display any obvious chaperone-binding motif, so we speculated that the proximal presence of the putative plasmodial ortholog of Sgt2/A (PfSgt2/A) with Hsp-recruitment potential to PfGet3-BirA^*-^HA ensured their biotinylation and enrichment in the BioID fraction. To identify the putative PfSgt2/A, we queried *P. falciparum* 3D7 proteome with amino acid sequences of ScSgt2 or HsSGTA. Results retrieved multiple hits with significant e-values, such as PF3D7_1434300 (Hsp70/Hsp90 organizing protein/PfHOP), PF3D7_1241900 (Tetratricopeptide repeat protein), PF3D7_0213500 (Tetratricopeptide-repeat protein), PF3D7_1355500 (Serine threonine protein phosphatase 5), PF3D7_0707700 (E3 ubiquitin ligase) and PF3D7_1334200 (Chaperone binding protein), respectively. Unfortunately, our LC-MS/MS analyses did not yield any peptide corresponding to PfHOP, however, we detected of 12, 4, 3 and 2 peptides corresponding to PF3D7_0707700, PF3D7_1334200, PF3D7_1355500 and PF3D7_1241900, respectively in the BioID fraction and none were detected in the control (**Supplementary Table 4**). Interestingly, all these proteins contain TPR domains and could likely recruit the Hs(c/p) cascade to function as a putative PfSgt2/A. Indeed, a subclass of TPR-domain containing proteins such as Protein phosphatase 5, HOP and Sti1 in yeast act as cochaperones in regulating the nucleotide hydrolysis cycles and connecting protein folding pathways with the alternate route to ubiquitination and degradation [129]. However, the involvement of PfHOP also cannot be ruled out solely on the basis of the absence of any detectable peptide in the ‘non-saturable’ LC-MS/MS and further experiments are required to identify and validate a cognate PfSgt2/A in *P. falciparum*.

The mammalian Bag6 complex also functions as an upstream loading factor, delivering TA substrates to the TRC40 (the mammalian ortholog of Get3) [28, 130]. Comprising of Bag6 (BCL-2-associated athanogene 6), TRC35 (HsGet4) and Ubl4A (Ubiquitin-like protein 4A), the Bag6 complex associates with the ribosomes and awaits the emergence of TMDs from the ribosomal tunnel [131]. The translating TA proteins interacts with HsSGTA via Bag6, thus triaging HsSGTA function to ER-associated degradation (ERAD) and proteasomal activity [83, 132–134]. We thus envisioned that the mammalian Bag6 complex in *P. falciparum* could possibly be biotinylated during their close proximity to the putative PfSgt2/A as it ‘transfers’ the TA substrates to PfGet3-BirA*-HA and accordingly get affinity-purified. PF3D7_0922100 was the top hit for *H. sapiens* Bag6 BLAST search against *P. falciparum* in PlasmoDB with an e-value of 2e-08. However, other hits reported from the BLAST query, like PF3D7_1113400, PF3D7_1365900, PF3D7_1313000 and PF3D7_1211800 also exhibited significant e-values of 1e-05, 3e-06, 2e-05 and 3e-04, respectively. Our BioID fraction detected 14 peptides corresponding to PF3D7_0922100, while none was detected in the control fraction (**Supplementary Table 4**). Among others, peptides corresponding to PF3D7_1365900 were detected both in the BioID and control fraction. However, the BioID fraction yielded 5.03-fold enrichment over the control fraction thereby maintaining PF3D7_1365900 as a valid Bag6 candidate. Surprisingly, BLAST query with the amino acid sequence of human Ubl4A also yielded plasmodial proteins that overlapped with the Bag6 hits, notably PF3D7_1211800, PF3D7_1365900, PF3D7_0922100, PF3D7_1113400 and PF3D7_1313000 with e-values of 8e-12, 3e-12, 3e-08, 1e-07 and 2e-08, respectively. PF3D7_1211800 is 381 amino acids in length and designated as Polyubiquitin (PfpUB); PF3D7_1365900 is a small protein of 128 amino acids in length and annotated as ubiquitin-60S ribosomal protein L40; PF3D7_0922100 is 1542 amino acids in length and annotated as a putative ubiquitin-like protein; PF3D7_1113400 is 388 amino acids in length and annotated as putative ubiquitin domain-containing protein DSK2; and PF3D7_1313000 is 76 amino acids in length and annotated as a putative Ubiquitin-like protein Nedd8 homolog. Similarly, BLAST search using the amino acid sequences of *S. cerevisiae* Get5 (ScGet5) identified PF3D7_1313000 in *P. falciparum* with e-value of 6e-06. No peptides corresponding to PF3D7_1313000 was detected in the BioID fraction. We reasoned that the overlapping hits between the BLAST output of Bag6, Ubl4A and ScGet5 were due to the shared ubiquitin-like domain feature and thus, we were unable to precisely identify or eliminate any candidate as a homolog of Bag6, Ubl4A or Get5 in *P. falciparum.* It is also possible that multiple proteins play redundant roles as Get5/Bag6/Ubl4A in *P. falciparum* and further experiments are necessary to identify and characterize the plasmodial homolog.

The terminal step in TA translocation involves interaction of the TA-loaded Get3 with Get1/2 (or the mammalian WRB/CAML) receptors at the ER membrane to achieve membrane insertion, a process coupled with the ATP hydrolysis [135–139]; reviewed in [91, 42, 140, 141]). During this process, the positively charged residues at N-terminus of the ER-membrane anchored Get2/CAML are proposed to function as an antenna, firstly seeking to capture the TA complex and subsequently promoting its disassembly for insertion into the ER membranes, facilitated by the insertase activity of Get1/WRB [136, 142]. The C-terminus of CAML, on the other hand, has been implicated to play an important role as a membrane anchor as well as contribute towards the stability of Get1/WRB receptor [143, 144]. We were unable to identify any prospective plasmodial homolog Get2/CAML in our BioID fraction based on the amino acid sequence similarity/identity. However, cross-kingdom members of Get2/CAML have recently been shown to display extreme variations in sequences; rather the members exhibited conservation in their apparent structural properties, namely: a positively charged N-terminus stretch (with at least four arginine or lysine residues in a stretch), the putative numbers of TMDs (three) and the topology of the protein (N-terminus_Cytosol_-TMDs-C-terminus_Luminal_) [145]. We therefore mined the *P. falciparum* 3D7 proteome in PlasmoDB to shortlist proteins fulfilling both the criteria, *i.e.,* (i) the presence of the RR8422 (Arg-Arg-Polar-Basic-Aliphatic-Aliphatic) motif (as a representation of the conserved RRRK motif commonly seen in CAMLs; [146]) within the first 50 amino acids from the N-terminus, and (ii) the presence of at least 3 TMDs (**Figure 6A**). Our results retrieved PFD_0215600 as the only hit. Interestingly, two peptides corresponding to PF3D7_0215600 were detected in the BioID fraction while none in the control (iBAQ fold enrichment >1000; see **Supplementary Table 4**) thereby indicating it as a potential homolog of Get2/CAML. In the PlasmoDB, PF3D7_0215600 is a 403 amino acid protein, without the ER-type SS and annotated as a conserved plasmodial protein of unknown function. It has three transmembrane regions spanning between 247-267, 299-322 and 348-368 amino acids and shared only 27% similarity and 14.4% identity with the *H. sapiens* CAML and 22.9% similarity and 12.4% identity with ScGet2 (**Figures 6B-C** and **Supplementary Figure 13**). Since the transmembrane topology prediction for PfGet2 by MEMSAT 3 [147] revealed that residues 1-246 are oriented towards the cytosol, we expressed and purified the recombinant cytosolic domain of PF3D7_0215600 (PfGet2^CD^; PF3D7_0215600 is designated henceforth as PfGet2) as protein fused to the MBP at the N-terminus (see *materials and methods*). Accordingly, purified PfGet2^CD^ or recombinant MBP (as a control) was incubated with purified PfGet3-6×his and followed by enrichment with Ni-NTA beads and western blotting using anti-MBP or anti-PfGet3 antibodies (**Figure 6B**). Results revealed co-purification of MBP-PfGet2^CD^ (but not MBP) with recombinant PfGet3-6×his bound to Ni-NTA. Thus, on the basis of our bioinformatics selection parameters, BioID data and *in vitro* interactions results, we concluded that the *P. falciparum* genome encodes for a functional PfGet2 which exhibits properties similar to other Get2 homologs.

**Figure 6.**
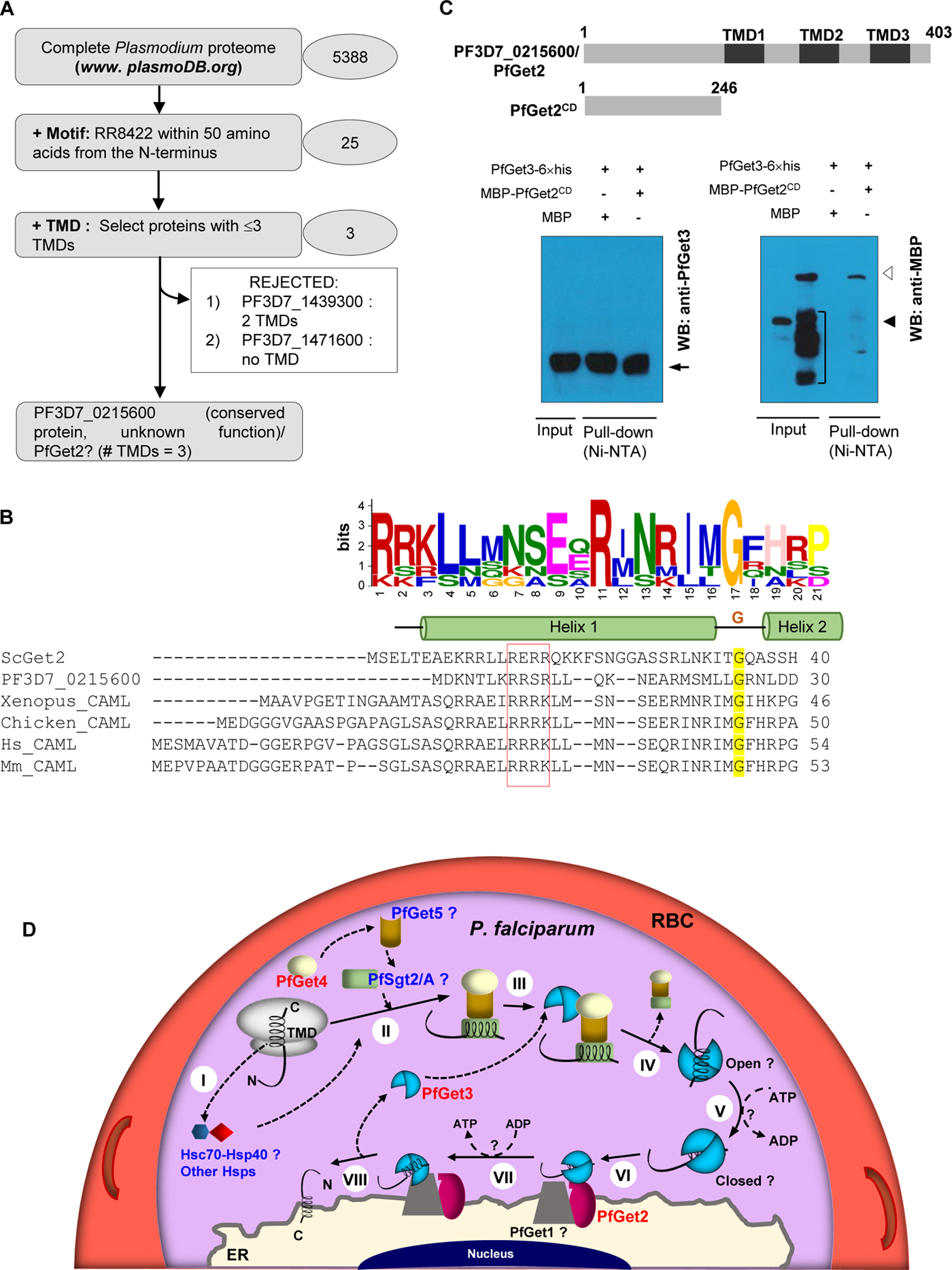
Identification of PfGet2, *in vitro* association with PfGet3 and schematic of the GET pathway in *P. falciparum* infected erythrocytes. **A.** Flow diagram showing the successive selection and elimination criteria in the bioinformatic prediction of PfGet2 in *P. falciparum* 3D7. **B.** Sequence alignments between the amino acids at the N-terminus of PF3D7_0215600/PfGet2, *S. cerevisiae* Get2 (ScGet2) and homologs of CAML by Clustal Omega (https://www.ebi.ac.uk/Tools/msa/clustalo/) and the representative logo generated by MEME suite (https://meme-suite.org/meme/tools/meme). The conservation of RERR motif in ScGet2 (implicated in the interactions with the negatively charged residues of ScGet3; [136]) with PfGet3 and corresponding residues in CAML are boxed in red. The two helices connected by a flexible glycine linker (yellow highlighted) is also shown in the above schematic. **C.** Schematic of PfGet2 and PfGet2^CD^ (top). Bottom: western blots showing the association of purified 6×his tagged recombinant PfGet3 (∼43-kDa; black arrow) with MBP-PfGet2^CD^ (empty arrowhead) but not with MBP (empty position indicated by solid black arrowhead) as revealed by pull-down assay using Ni-NTA beads followed by western blots using antibodies to MBP (right) or PfGet3 (left). Corresponding input and pull-down lanes are as indicated. Non-specific degradation products of MBP-PfGet2CD are demarcated by black square bracket (right). **D.** Cartoon representation summarizing the predicted role(s) of GET homologs in the TA-protein trafficking within *P. falciparum* parasites. The ribosome emerging TMD of the translating TA protein either intermediately enlists the Heat shock (cognate) protein repertoire (Hs(c/p); step I), or the PfSgt2/A (in a complex with PfGet4 and PfGet5; step II) directly to sequester the hydrophobic TMD from the hydrophilic cytosol. PfGet3 then interacts with PfGet4 (step III) to enable the transfer of TA protein, binding of the TMD with PfGet3 and the release of PfSgt2/A-PfGet4-PfGet5 complex (step IV). PfGet3 may then shift from an open to a tighter closed conformation, dependent on ATP binding (step V). Subsequently, the TA-bound PfGet3 approaches the ER-membrane receptors, where it interacts first with PfGet2 (step VI), followed by an ATP-hydrolysis event (to revert to the loosely TA-bound open conformation) and concomitant binding to PfGet1 (step VII). Finally (step VIII), PfGet1 functions as an insertase to mediate the release of the TA protein from (now loosely bound) PfGet3 and insertion into the ER membrane with the N-terminus facing the parasite cytosol. The homologs of the GET pathway in *P. falciparum* validated or shortlisted in this study are represented in red or blue text, respectively. The identity of the homolog of Get1/WRB in *P. falciparum* (PfGet1) remains unknown. ER, endoplasmic reticulum; RBC, red blood cell.

### 2.9 Concluding remarks

The outcome from this study revealed two novel aspects of protein trafficking in the human malaria parasite *P. falciparum*, namely the existence of TA proteins in the proteome, and the active role of GET machinery for the protein translocation onto destined organelles. We initiated this study with the bioinformatics-based identification of a *P. falciparum* homolog of Get3 (PfGet3) because Get3 is reported to operate at the crossroads of the GET trafficking pathway across both the prokaryotic and eukaryotic systems. Evaluations of the structural and functional features of PfGet3 *in silico* revealed conservations in all *Plasmodium* species as well as across related apicomplexan parasites, including *Toxoplasma*. Our cellular assays further showed dual localization of PfGet3 in *P. falciparum*; the parasite cytosol and at the perinuclear ER regions, reminiscent of the distribution profile expected for a functional Get3 homolog as it receives the TA substrates in communication with the soluble TRC in the cytosol, and the ER-proximal Get3 where it associates with membrane receptors to facilitate TA insertions. The yeast complementation assay further validated the functional equivalence of PfGet3 with the yeast counterpart.

A recent study identified 59 putative TA proteins encoded by the *Toxoplasma* genome [27]. Similarly, our bioinformatics predictions revealed a total of 63 TA proteins in the *P. falciparum*. We further validated these predictions in cellular assays using the BioID approach with PfGet3-mediated proximal labelling and showed that a subset of the TA repertoire is indeed associated with PfGet3, thus directly implicating the role of PfGet3 in their translocation. In addition, our in-depth mining and analyses of mass spectrometric data from the PfGet3-BioID pool either precisely identified or shortlisted a few other homologs of the GET machinery in *P. falciparum*. These included PfGet4 and PfGet2, which were also conclusively validated by *in vitro* binding assays using either the full-length recombinant protein (for PfGet4) or the predicted cytosolic domain (for PfGet2^CD^). We were, however, only able to shortlist (and not conclusively identify) the candidate plasmodial homologs of some of the other GET components due to the omnipresence of the following conserved domains/sequence motifs across multiple hits in the BioID fraction: (i) tetratricopeptide repeat region (TPR) in Sgt2/A and, (ii) ubiquitin-like domain in Get5/UBL4A/Bag6. We were also unable to identify Get1/WRB in *P. falciparum*, possibly due to the extreme sequence divergence from the canonical Get1/WRB. Identification and validation of PfGet1 along with homologs of the plasmodial Get5/UBL4A/Bag6 forms the next challenge in our understanding of GET pathway in this parasite. Until then, we offer our results to the community to investigate further alongside us.

## 3 MATERIAL AND METHODS

### Evolutionary analyses of PfGet3 and PfGet4 by Maximum Likelihood method

The evolutionary history of PfGet3 (PF3D7_0415000) and PfGet4 (PF3D7_1438600) were inferred by using the Maximum Likelihood method and Whelan and Goldman plus frequency model [125]. The bootstrap consensus tree inferred from 1000 replicates is taken to represent the evolutionary history of the taxa analyzed [148]. Branches corresponding to partitions reproduced in less than 50% bootstrap replicates are collapsed. The percentage of replicate trees in which the associated taxa clustered together in the bootstrap test (1000 replicates) have been shown next to the branches. Initial tree(s) for the heuristic search were obtained automatically by applying Neighbor-Join and BioNJ algorithms to a matrix of pairwise distances estimated using the JTT model, and then selecting the topology with superior log likelihood value. A discrete Gamma distribution was used to model evolutionary rate differences among sites [5 categories (+*G*, parameter = 4.1388 for PfGet3, and 66.6765 for PfGet4)]. The rate variation model allowed for some sites to be evolutionarily invariable ([+*I*], 0.46% sites) for PfGet4. The analysis involved 13 and 12 amino acid sequences for homologs of PfGet3 and PfGet4, respectively. There was a total of 409 (for PfGet3) or 327 (for PfGet4) positions in the final datasets. Evolutionary analyses were conducted in MEGA X [149].

### Plasmids and constructs

All plasmid constructs used for this study were generated using molecular biology grade reagents and verified by Sanger sequencing. For all the PCRs involving the amplification of plasmodial homologs of the GET pathway and sub-cloning into different plasmids, *P. falciparum* 3D7 genomic DNA was used as a template. Briefly, pET42b (PfGet3-6×his) was generated by PCR amplification of *pfget3* using the forward PfGet3-NdeIF (5’-AGATCTCATATGAGTGAGGATGAATCGAATTCCGTTTCTTGTTCATTAAGC-3’) and reverse PfGet3-XhoIR (5’-AGATCTCTCGAGCAGATTATCTTTATATATAGGTATATCTTTCGATTGTAAGAGCATCTCCG-3’) primers, digestion of the PCR product with NcoI-XhoI and cloning at corresponding sites within similarly digested pET42b. The yeast complementation plasmid pYES260 was purchased from Euroscarf (http://www.euroscarf.de/) and contained *ura3* selectable marker [150]. For generating the plasmid pYES260 (PfGet3), the *pfget3* was PCR amplified using PfGet3-NcoIF forward (5’-AGATCTCCATGGATAGTGAGGATGAATCGAATTCCG-3’) and PfGet3-XhoI reverse (5’-AGATCTGTCGACCTCGAGTTACAGATTATCTTTATATATAGGTATATCTTTCGATTG-3’) primers. The PCR product was digested with NcoI-XhoI and cloned at corresponding sites in digested pYES260. Similar strategy was also used to generate pYES260 (PfGet4/PF3D7_1438600) using PfGet4-HindIIF (5’-CCCATAAAGCTTATGTATCCATATGATGTTCCAGATTATGCTGCAGCTGCTAAAAAGTTCAAATTT AGTAAAGAAAAGCTAGCC-3’) and PfGet4-XhoISalIR (5’-AGATCTGTCGACCTCGAGTTATGCAAATATGTTTTGGAACATACTGAACAAGTTATG-3’) primers. Cloning within these sites also ensured incorporation of sequences encoding for the HA-tag at the N-terminus of the recombinant PfGet4 in the *S. cerevisiae* transfectants. The *P. falciparum* expression plasmid construct pA150 (PfGet3-GFP) was generated by PCR amplifying *pfget3* using PfGet3-AvrIIF (5’-GTACCGCCTAGGATGAGTGAGGATGAATCGAATTC-3’) and PfGet3-AAA-BglIIR (5’-TCCTTTAGATCTAGCTGCTGCCAGATTATCTTTATATATAGGTATATCTTTCGATTG-3’), restriction digestion and cloning into similarly digested pA150. For the BioID experiments, the plasmid pcDNA3.1 MCS-BirA(R118G)-HA was purchased from Addgene and used to construct pA150 (PfGet3-BirA*-HA) through a series of subcloning steps. Briefly, *pfget3* was PCR amplified using PfGet3-AvrIIF and PfGet3-AAA-BglIIR primers, first cloned at NheI-BamHI sites of pcDNA3.1 MCS-BirA(R118G)-HA. Subsequently, this plasmid used as template to PCR amplify out the entire *pfget3-birA (R118G)-ha* fragment using PfGet3-AvrIIF and HA-XhoIR (5’-TCTAGACTCGAGCTATGCGTAATCCGGTACATCGTAAG-3’) primers, digested with AvrII-XhoI and subcloned into similarly digested pA150.

The constructs for the expression of N-terminus MBP tagged recombinant PfGet4 or PfGet2^CD^, *i.e.*, pMALc2X (PfGet4) or pMALc2X (PfGet2^CD^) were generated by PCR amplification of either full-length *pfget4* or nucleotides encoding for the first 246 amino acids of PfGet2^CD^ using PfGet4-EcoRIF (5’-GTACCGGAATTCATGAAAAAGTTCAAATTTAGTAAAGAAAAGCTAGCC-3’)/PfGet4-XhoISalIR or PfGet2CD-BamHIF (5’-GTACGCGGATCCATGGATAAAAATACATTAAAAAGAA-3’)/PfGet2CD-XhoISalIR (5’-AGACCGGTCGACCTCGAGTTATTCATGTTTCGTAATAATAAATTG-3’) primer pairs and subsequently cloning at corresponding sites in pMALc2X plasmid (New England Biolabs).

### Expression and purification of recombinant PfGet3-6**×**his, MBP, MBP-PfGet4 and MBP-PfGet2^CD^ from *E. coli*

*E. coli* BL21 arabinose-inducible (AI) cells (Invitrogen) were transformed with the plasmids pET42b (PfGet3-6×his), pMALc2X (PfGet4) or pMALc2X (PfGet2^CD^) and transformants were selected on 100 µg/ml ampicillin-containing LB plates. Primary cultures of transformed *E. coli* cells were grown overnight at 37°C under shaking conditions and then used to seed secondary cultures. Cultures were induced with 1 mM isopropyl β-D-1-thiogalctopyranoside (IPTG) and 0.2% L-Arabinose at OD_600_ ≍ 0.6 for 16 h at 25°C. For the purification of PfGet3-6×his, induced cultures were harvested at 6,000 rpm for 20 mins and resuspended in lysis buffer (50 mM Tris. HCl pH 8.0, 500 mM NaCl, 10 mM imidazole,1 mM MgCl_2_, 5mM β-mercaptoethanol, 5% glycerol) containing 100 mM Phenylmethylsulphonylfluoride (PMSF) and EDTA-free Protease inhibitor cocktail tablets (Roche). Resuspended cells were first treated with 1 mg/ml Lysozyme to digest the cell wall and subsequently frozen at −80°C overnight. The thawed cell suspension was sonicated at 30% amplitude with repetitive pulses of 30 s and off for 30 s till clarity. The sonicated cells were centrifuged at 14,000 rpm for 1 h at 4°C to remove the cell debris and the supernatant was incubated to Ni^2+^-NTA resin (Thermo Fisher) pre equilibrated with lysis buffer. After binding 4°C for 4 h under rotating conditions, the supernatant-resin suspension was packed onto a column and washed with at least 20 column volumes of wash buffer (lysis buffer containing 40 mM imidazole). PfGet3-6×his was eluted by elution buffer (lysis buffer containing 300 mM imidazole) and the eluted fractions were analyzed by running SDS-PAGE. Fractions containing PfGet3-6×his were pooled, dialyzed against 10 mM Tris. HCl pH 8.0, 150 mM NaCl, 1 mM MgCl_2_, 5 mM β-mercaptoethanol and 5% glycerol with minimum of three changes in buffer and concentrated using membrane concentrators (10-kDa MWCO; Millipore). The purified PfGet3-6×his was subsequently used for the custom generation of anti-PfGet3 antibodies or for *in vitro* binding experiments.

For the purification of recombinant MBP, MBP-PfGet4 and MBP-PfGet2^CD^, the respective secondary cultures (OD_600_ ≍ 0.6) of transformed AI cells were induced with 0.3 mM IPTG at 25°C for 16 h, harvested at 6,000 rpm for 20 min and resuspended in column buffer (20 mM Tris. HCl, pH 7.4, 200 mM NaCl, 1 mM EDTA and 10 mM β-mercaptoethanol). Soluble supernatant fractions were generated following similar protocol as described above. MBP, MBP-PfGet4 and MBP-PfGet2^CD^ were affinity purified using amylose-resin (Thermo Fisher) and eluted with elution buffer (50 mM Tris. HCl pH 8.0, 200 mM NaCl, 1 mM EDTA, 5 mM β-mercaptoethanol and 10 mM maltose) as according to manufacturer’s instructions. The corresponding proteins were dialysed against 10 mM Tris. HCl pH 8.0, 150 mM NaCl, 1 mM MgCl_2_, 5 mM β-mercaptoethanol and concentrated. The purity of each protein was assessed by SDS-PAGE and subsequently used for *in vitro* binding studies.

### Generation of custom antibodies against PfGet3 and PfGet4

Antibodies were generated against PfGet3 and PfGet4. Anti-PfGet3 antibodies were custom-generated in rabbits by Ms. Biobharati LifeScience Inc. using the purified PfGet3-6×his as immunogen. Following booster, the anti-PfGet3 antibodies were affinity purified from harvested sera using column bound recombinant PfGet3-6×his. For anti-PfGet4 antibodies, the KLH-conjugated peptide CPPNSTEEKYKFMN, corresponding to the amino acids 97-110 in PfGet4 (PF3D7_1438600), was used as an immunogen in New Zealand rabbits and custom-generated by Genscript Inc. Antibodies were also affinity purified against the same peptide.

### Transformation of *S. cerevisiae* BY4741 wild type, *Δget3* or *Δget4* knockout strains with *pfget3* or *pfget4*

S. cerevisiae parental BY4741 (S288C isogenic yeast strain: MATa; his3D1; leu2D0; met15D0; ura3D0), Δget3 (BY4741; MATa; ura3Δ0; leu2Δ0; his3Δ1; met15Δ0; YDL100c::kanMX4) or Δget4 (BY4741; MATa; ura3Δ0; leu2Δ0; his3Δ1; met15Δ0; YOR164c::kanMX4) strains were purchased from Euroscarf (http://www.euroscarf.de/). The constructs pYES260 (PfGet3), pYES260 (PfGet4) or empty plasmid pYES260 were transformed in either Δget3 or Δget4 strains, respectively. All the yeast transformations were performed using previously published protocol [151] and selected in media lacking uracil. Following transformation, the cells were grown overnight in Yeast Nitrogen Base (YNB) Sc-Ura media at 30°C under shaking conditions.

To verify the expression of recombinant PfGet3 or PfGet4 in transformed *S. cerevisiae*, cells were inoculated in YNB Sc-Ura media and grown. Cell density was subsequently adjusted to OD_600 nm_ ≍ 0.01 and induced with 2.5% galactose and 1% raffinose for 8 h following which cells were harvested at 6,000 rpm for 5 min at 25°C and washed with autoclaved milliQ. Cells were then resuspended in lysis buffer (50 mM Tris.HCl, pH 7.5, 1% sodium deoxycholate; 1% Triton X-100; 0.1% SDS; 50 mM sodium fluoride; 0.1 mM sodium vanadate; 5 mM sodium pyrophosphate) containing 0.05% PMSF and EDTA-free Complete protease inhibitor cocktail (Roche). Equal volume of acid washed 0.45 mm glass beads (Sigma) were added to the cells and homogenized using BeadBeater (BioSpec) at 3,000 rpm for 30 s followed by 30 s on ice for a total of 8 cycles. Cell suspensions were decanted away from the glass beads and centrifuged twice at 10,000 rpm at 4°C to remove unlysed cells and cellular debris. Supernatants were quantified and analyzed by western blotting using anti-PfGet3 or anti-PfGet4 antibodies.

### Quantitative growth assay of parental BY4741 and *Δget3* or *Δget4* transformants

The *S. cerevisiae* parental BY4741 strain or the *Δget3* or *Δget4* knockouts transformed with either the empty pYES260 plasmids or expressing PfGet3 or PfGet4 were grown overnight in YNB Sc-Ura at 30°C under shaking conditions at 200 rpm. After growth, the respective cell density was adjusted to OD_600 nm_ ≍ 0.01, cells transferred to 50 ml falcon tube containing fresh 15 ml YNB SC-Ura and induced with 2.5% galactose and 1% raffinose in the presence or absence of 1.5 mM CuSO_4_. Cells were grown for 32 h at 30°C under shaking conditions during which samples were harvested at every 8 h time interval to measure OD_600 nm_ and the growth curve was plotted accordingly.

### Culturing of *P. falciparum* parasites, generation of transgenic strains and microscopy

*P. falciparum* 3D7 parasites were grown in fresh O+ human RBCs (kindly provided by the Rotary Blood Bank) in RPMI 1640 media supplemented with 0.5% Albumax II, 0.2 mM hypoxanthine, 11 mM glucose, 5% NaHCO_3_ and 50 μg/mL gentamycin in a humidified CO_2_ incubator as described previously [65]. Cultures were monitored daily by Giemsa staining of methanol-fixed smears and fed as necessary. The parasitemia was generally maintained below 12% for healthy culture growth. Transgenic parasites were generated by electroporation of plasmid DNA according to previously published protocols [152]. For live cell imaging of transgenic parasites expressing PfGet3-GFP as well as for immunofluorescence microscopy, the parasite nucleus was stained with Hoechst 33342, processed as described previously [153] and imaged with a 100× NA objective on a Zeiss ApoTome 2 (Axiovert 40 CFL) under fluorescence and brightfield optics.

### Indirect Immunofluorescence assay (IFA) of *P. falciparum* parasites

Indirect immunofluorescence assays (IFA) were performed with either non-transfected *P. falciparum* 3D7 or transgenic parasites following protocol as described previously [154]. Briefly, fixed smears of parasites were fixed with 4% paraformaldehyde/0.0075% glutaraldehyde and permeabilized with 0.1% Triton X-100. The free aldehyde group was neutralized with 50 mM NH_4_Cl and slides blocked with 2% BSA. The respective primary antibodies were diluted in blocking buffer and incubated at RT for 2 h. The following antibody concentrations were used: anti-PfGet3, 10 µg/ml; anti-Exp2 ([152]); 20g/µg/ml; anti-BiP ([152]); 10 µg/ml and anti-plasmepsin V (MRA-815A, MR4), 1:200 dilution. The corresponding FITC- or TRITC-labelled secondary IgG antibodies (ICN Biochemicals) were used at 1:200 dilution. The parasite nuclei were stained with 5 μ ml Hoechst 33342 (Molecular Probes) and slides were mounted with DABCO. The slides were viewed as described previously.

### SDS-PAGE and western blotting

The preparation of lysates from transformed bacterial and yeast cultures are described previously. For the preparation *P. falciparum* parasites, cultures are 10-12% parasitemia were permeabilized with PBS (137 mM NaCl, 2.7 mM KCl,10 mM Na_2_HPO_4_, 2 mM KH_2_PO_4_, pH 7.4) containing 0.05% saponin to release the hemoglobin content of the host red blood cell. The parasite fraction was repeatedly washed (at 3,200 rpm for 10 min) with ice cold PBS till no hemoglobin was detected in the supernatant fraction. The parasite pellet was subsequently solubilised in Laemmli’s sample buffer [155] containing 1 mM dithiothreitol (DTT), denatured for 10 min at 95°C and resolved by 12% SDS-PAGE. Resolved proteins were then transferred onto a nitrocellulose membrane and blocked with 5% fat-free skim milk for 2 h. Membranes were incubated with either anti-PfGet3, anti-Hog (kindly provided by Prof. A. Mondal), anti-GFP (Invitrogen) or anti-β actin antibodies for 3 h on a platform shaker and followed by respective HRP-conjugated secondary antibodies. The blots were finally developed by chemiluminescence detection (Amersham).

### Proximity labelling (BioID) experiments

Proximity biotinylation experiment was performed with the transgenic *P. falciparum* 3D7 parasites expressing PfGet3-BirA*-HA following previously published protocol [156]. Briefly, 140 ml of asynchronous cultures were split equally into two 175 cm^2^ flasks at 2% parasitemia and 5% hematocrit. D-Biotin was added at a final concentration of 50 µg/ml to one of the flasks (designated as the BioID sample) and not to the other (representing the control). The parasites were allowed to grow for 48 h at 37°C with replacement of fresh media (± D-Biotin) after 24 h. Cultures were subsequently harvested, washed twice with cold PBS and RBCs were lysed in PBS containing 0.03% saponin for 10 min. The parasite pellet was washed at least five times with 10 volumes of cold PBS to remove all the hemoglobin and stored at −80°C till further use. When planned, the frozen parasites were thawed at 4°C and extracted for 1 h at 4°C with 2 ml lysis buffer (50 mM Tris. HCl pH 7.5, 500 mM NaCl, 1% Triton-X-100) containing 1 mM DTT, 1 mM PMSF and protease inhibitor cocktail (Roche) under rotating conditions. The soluble extract was separated from insoluble cellular debris by centrifugation at 16,000 × g for 30 min at 4°C and immediately diluted two-fold with 50 mM Tris. HCl, pH 7.5. An aliquot each of the BioID and control extracts was stored at −80°C for SDS-PAGE and western blotting. To the remaining extracts, 30 µl of StrepTactin XT was added and allowed to incubate overnight under rotating condition at 4°C. After incubation, the beads were pelleted by centrifugation at 3,000 rpm for 5 min at 4°C and washed sequentially: thrice with lysis buffer, twice with cold double distilled water and twice with 50 mM Tris. HCl, pH 7.5. The beads were finally boiled in SDS-PAGE sample buffer and processed either for western blotting using anti-PfGet3 or anti-HA, or for LC-MS/MS analyses.

### Mass Spectrometry analyses and peptide quantitation

Mass spectrometry of the control and BioID fractions were performed at Wistar Proteomics Facility, Philadelphia, PA, USA. Briefly, the fractions were resolved by SDS-PAGE (only to the extent when the samples entered 0.5 cm within the resolving gel) and stained with colloidal Coomassie. Bands were excised from the gel, de-stained with water, alkylated with iodoacetamide, and digested with trypsin as previously described [65]. The digests were analyzed by LC-MS/MS using reverse phase capillary HPLC with a 75 µm nano-column interfaced on-line with a Thermo Electron LTQ OrbiTrap XL mass spectrometer. The mass spectrometer was operated in data dependent mode with full scans performed in the Orbitrap and parallel MS/MS analysis of the six most intense precursor ions in the linear ion trap. MS data were searched with full tryptic specificity against the UniProt *P. falciparum* 3D7 database and a common contaminant database using MaxQuant 1.6.17.0. The custom database consisted of a combination of the expected sequences with the various mutations of the proteins used in this experiment, the *E. coli* proteome (to provide a reasonable sized database), and expected common contaminants including keratins, trypsin, etc. A reversed sequence database was appended to the front of the forward database and used to estimate peptide false positive rates. Data were searched using partial tryptic specificity, a maximum of three missed cleavages, mass tolerance of 100 ppm, cysteine fixed as the carboxymethyl derivative, and dynamic methionine oxidation and N-terminal acetylation. Resulting data was filtered on 5 ppm and dCn of 0.07. The false positive rate for peptide identifications using these database search and data filtering parameters was set at 1%. Variable modification search included Biotinylation (+226.0776) on Lysines, Glycine-Glycine (+114.0429) on Lysines, *i.e*., tryptic remnant of ubiquitinylation or neddylation, acetylation (+42.01056) on protein N-terminus and oxidation (+15.99491) on Methionines. Fixed modifications searched include carbamidomethylation (+57.02146) on Cysteines. Protein quantification was performed using Razor + unique peptides. Razor peptides were the shared (non-unique) peptides assigned to the protein group with the most other peptides (Occam’s razor principle). MS/MS count referred to how many times peptides belonging to the protein were sequenced and Intensity denoted the sum of the peptide MS peak areas for the protein. The iBAQ (Intensity Based Absolute Quantification) values represented protein intensity divided by the number of theoretical peptides and are roughly considered to be proportional to the molar quantities of the proteins.

### *In vitro* binding studies with recombinant PfGet3-6×his, MBP, MBP-PfGet4 and MBP-PfGet2CD

For *in vitro* binding studies, purified recombinant MBP-PfGet4 or MBP-PfGet2^CD^ (2-5 µg) were incubated with recombinant PfGet3-6×his in 1 mg/mL BSA containing binding buffer (50 mM Tris. HCl, pH 8.0, 200 mM NaCl, 0.1% NP-40, 20 mM Imidazole and 1 mM DTT) for 2 h at 4°C. As a control, PfGet3-6×his was also incubated with purified MBP at similar concentration and duration. Following binding, binding buffer-equilibrated Ni^2+^-NTA beads were added to all protein mixtures and allowed to bind at 4°C for 2 h under shaking conditions. Beads were then washed extensively with binding buffer (without BSA), boiled in the presence of SDS-PAGE sample buffer for 5 min at 95°C and resolved by SDS-PAGE. Nitrocellulose membrane-transferred proteins were incubated with anti-PfGet3 or anti-MBP (NEB), probed with corresponding HRP-conjugated secondary antibodies and developed by chemiluminescence.

## Acknowledgements

The authors would like to thank Dr. Hsin-Yao Tang at the Wistar Proteomics Core Facility, The Wistar Institute, Philadelphia, PA, USA for help in LC-MS/MS and analyses. This work was supported by funding from the Science and Engineering Research Board, Department of Science and Technology, Government of India (DST ECR/2015/000387), Department of Biotechnology Ramalingaswami Re-entry Fellowship (BT/HRD/35/02/ 2006) and University for Potential Excellence–II (Project ID 245) (S.B.). T.K., S.M. and A.R. are recipients of research fellowships from the University Grant Commission, Council of Scientific and Industrial Research and Indian Council of Medical Research, Government of India, respectively.

## Author contributions

T.K., S.M., A.R. and S.B. designed, performed, and interpreted the experimental work; S.B. provided key intellectual insight into the aspects of this study; T.K. and S.B. wrote the manuscript; and all authors commented on the manuscript.

---

**Supplementary Figure 1. Percentage identity and similarity of PfGet3 with other homologs of Get3. A.** Table summarizing the similarity and identity between the amino acid sequences of PfGet3 (PF3D7_0415000) in comparison to other validated or predicted homologs of Get3. **B-O.** Amino acid alignment (https://www.ebi.ac.uk/Tools/psa/emboss_needle/) between PfGet3 and the putative *P. berghei* Get3 homolog PBANKA_0717000 (B), putative *P. vivax* Get3 homolog PVP01_0521500 (C), putative *P. knowlesi* Get3 homolog PKNH_0506600 (D), putative *P. chabaudi* Get3 homolog PCHAS_0726100 (E), putative *P. yoelii* 17X Get3 homolog PY17X_0717200 (F), *S. cerevisiae* Get3 YDL100C (G), *S. pombe* Get3 SPAC1142.06 (H), *H. sapiens* ASNA1 ENSG00000198356 (I), *T. gondii* ASNA1 TGVEG_231190 (J), *L. donovani* ASNA1 LDBPK_110710 (K), *T. cruzi* ASNA1 Tc00.1047053507763.30 (L), *A. thaliana* Get3 At1g01910 (M), bacterial Arsenite transporter ArsA (N) and *C. parvum* (strain Iowa II) Get3 Cgd7_4070 (O).

**Supplementary Figure 2. Sequence alignments between a few representative Get3 homologs. A.** Sequence alignments of the three eukaryotic (PF3D7_0415000/PfGet3, ScGet3, HsGet3) and one prokaryotic (ArsA) Get3 homologs. The four conserved ATPase sequence motifs are boxed in rectangles and the zinc-binding ‘ÇxxC’ motif is boxed in oval. The ∼20 residue Get3/TRC40 insert is underlined in blue, and the conserved hydrophobic residues involved in forming the TMD-binding groove are labeled above with red dots [104]. **B.** Sequences from a few Get3 homologs were aligned using ClustalX [158]. Residue colouring is based on the program output (type of amino acid). The shading of the bars from brown to yellow reflects the conservation number, quality, and consensus amino acids of the ordinates. Occupancy at a particular residue position is indicated by increasing intensity of light to dark grey shading.

**Supplementary Figure 3. Expression of recombinant PfGet3-6**×**his in *E. coli*.** SDS PAGE (left) and western blot (right) showing the expression of recombinant 6×his tagged PfGet3 in *E. coli* cells only under IPTG induced conditions, as compared to the uninduced control. The induced recombinant PfGet3 is indicated as filled arrowhead in the SDS PAGE (left) and by empty arrowhead in the western blot (right) using custom-generated antibodies to PfGet3. Molecular weight standards (in kDa) are as indicated.

**Supplementary Figure 4. Secondary structure prediction for PfGet3. A.** Predicted secondary structure of PfGet3 by the Phyre2 server (www.sbg.bio.ic.ac.uk/phyre2) revealing the presence of 51% α helices, 10% β-strands and 21% disordered regions. Residues are colored according to a simple property-based scheme: A, S, T, G and P; small/polar are in yellow, M, I, L and V; hydrophobic are in green, K, R, E, N, D, H and Q; charged are in red, and W, Y, F, C; aromatic + cysteine are in purple. The secondary structure prediction comprises three states: α-helix, β-strand or coil. Green-sthelices represent α-helices, blue arrows indicate β-strands and faint lines indicate coils. The ‘SS confidence’ line indicates the confidence in the prediction from PSIPRED, with red indicating high confidence and blue showing low confidence. A large amount of blue or green in the confidence line is indicative of few homologous sequences detected and a consequent low probability of modeling success. **B.** Outcome of the TMD prediction for PfGet3 by the TMHMM server (www.cbs.dtu.dk/services/TMHMM-2.0). No transmembrane helix was predicted in PfGet3.

**Supplementary Figure 5. Cumulative hits of structural homologs of PfGet3 from the PDB.** Hits were retrieved by the Phyre2 server (www.sbg.bio.ic.ac.uk/phyre2). The PDB ID and 3D structure of each template is shown along with the percentage confidence and identity in decreasing order.

**Supplementary Figure 6. Ligand binding prediction for PfGet3 and ScGet3.** The nucleotide binding pocket of PfGet3 is shown (right) in comparison to that of ScGeT3 (left) with residues shown as sticks, generated by the GalaxyWEB server ((http://galaxy.seoklab.org) [106]. The lengths of hydrogen bonds (in Å) are as indicated.

**Supplementary Figure 7. Predicted homomers of PfGet3. A.** Predicted dimeric and tetrameric associations of PfGet3 as compared to the crystal structures available in the PDB. The interface area (in Angstrom^2^) between one chain and the other chains is calculated using the Naccess program (http://www.bioinf.manchester.ac.uk/naccess/). Sequence identities between query and template proteins are shown and ranges from 0 (totally different) to 100 (identical). Structural similarity between the query and the template protein is also shown (as measured by TM-align) and ranges from 0 (totally different) to 1 (identical). **B.** PfGet3 dimer illustrated by surface representation in PyMOL with individual chains colored orange and grey, respectively. The ELLE motifs at each dimer interface are shown in red (arrows). **C.** Putative PfGet3 tetramer in PyMOL surface representation with hydrophobic, polar, positive- and negative-charged residues colored green, orange, blue and red, respectively. Magnified image of the hydrophobic pocket (dotted boxed) region is shown at the right (in 50% transparency mode along with the cartoon representation).

**Supplementary Figure 8. Sequence alignment of a few representative homologs of Get4. A.** Table showing the percentage identity and similarity between the various homologs of Get4 as compared to PfGet4. Corresponding alignments, as shown below, were generated using EMBOSS needle (https://www.ebi.ac.uk/Tools/psa/emboss_needle/). **B.** Multiple sequences were also aligned using ClustalX [158]. Residue colouring is based on the program output (type of amino acid). The shading of the bars from brown to yellow reflects the conservation number, quality, and consensus amino acids of the ordinates. Occupancy at a particular residue position is indicated by increasing intensity of light to dark grey shading.

**Supplementary Figure 9. Alignment of PFD_1438600/PfGet4 and *S. cerevisiae* Yor164c/ScGet4.** Clustal alignment showing conservation of residues (red asterisks) between PfGet4 and ScGet4 implicated in interactions with Get3 [123].

**Supplementary Figure 10. Secondary structure prediction for PfGet4.** Results from secondary structure prediction for PfGet4 by the Phyre2 server (www.sbg.bio.ic.ac.uk/phyre2) revealing the presence of 72% α-helices and 14% disordered regions. Residues are colored according to a simple property-based scheme: A, S, T, G and P; small/polar are in yellow, M, I, L and V; hydrophobic are in green, K, R, E, N, D, H and Q; charged are in red, and W, Y, F, C; aromatic + cysteine are in purple. The secondary structure prediction comprises three states: α elix, β-strand or coil. Green helices represent α helices, blue arrows indicate β-strands and faint lines indicate coils. The ‘SS confidence’ line indicates the confidence in the prediction from PSIPRED, with red indicating high confidence and blue showing low confidence. A large amount of blue or green in the confidence line is indicative of few homologous sequences detected and a consequent low probability of modeling success.

**Supplementary Figure 11. Cumulative hits of structural homologs of PfGet4 from the PDB.** Hits were retrieved by the Phyre2 server. The PDB ID and 3D structure of each template is shown along with the percentage confidence and identity in decreasing order.

**Supplementary Figure 12. Surface representation conserved residues in PfGet4 and other Get4 homologs.** The degree of sequence conservation within Get4 proteins is mapped onto the surface of the PfGet4 structure in three different orientations: the convex surface (left panel), the tip (middle) and the concave surface (right). Red, dark orange, light orange, dark yellow and light yellow indicate residues with high or gradually decreasing degree of conservation, respectively. The clefts are indicated by arrows. The figure is adapted from a similar representation in [124].

**Supplementary Figure 13. Alignment between the amino acid sequences of PfGet2 and *H. sapiens* CAML.** Alignments were performed in Clustal Omega server (https://www.ebi.ac.uk/Tools/msa/clustalo/).

**Supplementary Table 1.** List of the total 130 predicted TA proteins in the *P. falciparum* 3D7 proteome.

**Supplementary Table 2. List of the 67 misrepresented proteins in the TA proteome of *P. falciparum* 3D7**. The list includes RIFINs (64), EVP1, REX-2 and MSP5.

**Supplementary Table 3. Final list of the 63 predicted TA proteins in the *P. falciparum* 3D7 proteome.** The PlasmoDB ID, description and other features are shown for each predicted TA protein. Corresponding GRAVY and Adagir scores are also shown and shaded according to the scale provided. Intracellular localization is also predicted for each TA protein based on three different machine learning tools (LOCKTREE 3, BUSCA and DeepLoc 1.0) and reveals no clear consensus for any particular organelle. Thus, the 63 predicted TA proteins were manually grouped (in this study) into three broad categories based on their Gene Ontology (GO) annotations in the Uniprot database (http://uniprot.org): ER (shaded light orange), mitochondria (grey) and other destinations (not shaded). Thus, a total of 14 ER-specific TA proteins, 7 mitochondrial-specific TAs and 42 TAs with diverse cellular destinations were predicted.

**Supplementary Table 4.** Raw data of the LC-MS/MS analyses of the BioID and control fractions.

