## Supplementary Figure 1 for "A conserved Guided Entry of Tail-anchored pathway is involved in the trafficking of tail-anchored membrane proteins in *Plasmodium falciparum*"

**A**

| | ID | Plasmodium falciparum 3D7 ASNA1 Homolog (Pf3D7_0415000) | Identity | Similarity |
|---|---|---|---|---|
| **01** | PBANKA_0717000 | *Plasmodium berghei* ASNA1 Homolog | 81.4% | 90.6% |
| **02** | PVP01_0521500 | *Plasmodium vivax* ASNA1 Homolog | 84.2% | 93.4% |
| **03** | PKNH_0506600 | *Plasmodium knowlesi* ASNA1 homolog | 83.9% | 93.4% |
| **04** | PCHAS_0726100 | *Plasmodium chabaudi* ASNA1 homolog | 79.8% | 90.3% |
| **05** | PY17X_0717200 | *Plasmodium yoelii* ASNA1 homolog | 81.4% | 89.5% |
| **06** | YDL100C | *Saccharomyces cerevisiae* GET3 | 35.9% | 46.9% |
| **07** | SPAC1142.06 | *Schizosaccharomyces pombe* GET3 | 41.8% | 54.4% |
| **08** | ENSG00000198356 | *Homo sapiens* ASNA1 Homolog | 40.8% | 59.7% |
| **09** | TGVEG_231190 | *Toxoplasma gondii* ASNA1 Homolog | 45.5% | 61.1% |
| **10** | LDBPK_110710 | *Leishmania donovani* ASNA1 homolog | 31.0% | 43.2% |
| **11** | Tc00.1047053507763.30 | *Trypanosoma cruzi* ASNA1 homolog | 36% | 49.2% |
| **12** | At1g01910 | *Arabidopsis thaliana* GET3 | 34.1% | 51.0% |
| **13** | ArsA | Bacterial Arsenite transporter ArsA | 16.3% | 37.0% |
| **14** | Cgd7_4070 | *Cryptosporidium parvum* (strain Iowa II) | 47.6% | 64.2% |

**B**

**PF3D7_0415000 vs. PBANKA_0717000**

```
PF3D7_0415000      1 MSE--DESNSVSCSLSLESDGYSDEEYDTNLNKLIENESLNWIFVGGKGG     48
                     ||:  .:.:|:||||||:||...||.|:|||||||||.||||||||||||
PBANKA_071700      1 MSKAGSDVSSISCSLSLDSDSCEDEYYETNLNKLIENTSLNWIFVGGKGG     50

PF3D7_0415000     49 VGKTTTSCSIAVQLSKRRESVLLLSTDPAHNTSDAFNQKFTNQPTLINSF     98
                     |||||||||||:||:|:|||||||||||||||||||||||||:|||||||
PBANKA_071700     51 VGKTTTSCSIAIQLAKKRESVLLLSTDPAHNTSDAFNQKFTNKPTLINSF    100

PF3D7_0415000     99 DNLYCMEIDTNYSENTAFKLNKKEMFDNILPELLHSFPGIDEALCFAELM    148
                     ||||||||||.:||:||||:||.:.|::|:||||.|||||||||||||||
PBANKA_071700    101 DNLYCMEIDTTFSEDTAFKINKSDFFNSIIPELLQSFPGIDEALCFAELM    150

PF3D7_0415000    149 QSIKNMKYSVIVFDTAPTGHTLRLLAFPDLLKKALGYLINIREKLKGTLN    198
                     |||||||||||||||||||||||||||||||||||||||::||||||||:
PBANKA_071700    151 QSIKNMKYSVIVFDTAPTGHTLRLLAFPDLLKKALGYLINLKEKLKGTLS    200

PF3D7_0415000    199 VLKNFTNNEMEFDSLYEKINHLNAMSSSIQANFQNPMKTTFVCVCIPEFL    248
                     :|::.|||||||:.:|:||||||.||.|||.|||||:|||||||||||||
PBANKA_071700    201 MLQSLTNNEMEFEGMYDKINHLNTMSISIQENFQNPLKTTFVCVCIPEFL    250

PF3D7_0415000    249 SVYETERLIQELTKKNISCYNIVVNQVVFPLDSPNVNLENCKNLLSQIKN    298
                     |||||||||||||||||||||||||||||||.||:||:|.|:.||.|||:
PBANKA_071700    251 SVYETERLIQELTKKNISCYNIVVNQVVFPLTSPDVNIEKCEKLLKQIKD    300

PF3D7_0415000    299 EQIQSYFNDLISKTEELEDVYISRRKLQSKYLTQIKNLYSNDFHIVCMPQ    348
                     ..||:.||.||.|.:||||||||||||||||||||||||.|.||||||||
PBANKA_071700    301 TNIQNSFNSLILKAKELEDVYISRRKLQSKYLTQIKNLYGNYFHIVCMPQ    350

PF3D7_0415000    349 LKNEIRGLNNISSFSEMLLQSKDIPIYKDNL    379
                     ||.|||||:.||:||||||||||||||..
PBANKA_071700    351 LKTEIRGLDKISNFSEMLLQSKDIPIYST--    379
```

#### Supplementary Figure 1 (contd.)

**C**

##### PF3D7_0415000 vs. PVP01_0521500

```
PF3D7_0415000      1 MSEDESNSVSCSLSLESDGYSDEEYDTNLNKLIENESLNWIFVGGKGGVG     50
                     ||  :::|:||||:||||.|.:|||||||:||:||::|||||||||||||
PVP01_0521500      1 MS--DADSLSCSLTLESDEYDEEEYDTNLSKLLENKTLNWIFVGGKGGVG     48

PF3D7_0415000     51 KTTTSCSIAVQLSKRRESVLLLSTDPAHNTSDAFNQKFTNQPTLINSFDN    100
                     ||||||||||||:|||||||||||||||||||||||||||||||||||||
PVP01_0521500     49 KTTTSCSIAVQLAKRRESVLLLSTDPAHNTSDAFNQKFTNQPTLINSFDN     98

PF3D7_0415000    101 LYCMEIDTNYSENTAFKLNKKEMFDNILPELLHSFPGIDEALCFAELMQS    150
                     ||||||||.|||||||||||.|.||||:||||.|||||||||||||||||
PVP01_0521500     99 LYCMEIDTTYSENTAFKLNKTEFFDNIIPELLQSFPGIDEALCFAELMQS    148

PF3D7_0415000    151 IKNMKYSVIVFDTAPTGHTLRLLAFPDLLKKALGYLINIREKLKGTLNVL    200
                     ||||||||||||||||||||||||||:|||||||||||:|||||||||:|
PVP01_0521500    149 IKNMKYSVIVFDTAPTGHTLRLLAFPELLKKALGYLINLREKLKGTLNML    198

PF3D7_0415000    201 KNFTNNEMEFDSLYEKINHLNAMSSSIQANFQNPMKTTFVCVCIPEFLSV    250
                     |:|||||||.:.:|||||||||||.|||:|||||:|||||||||||||||
PVP01_0521500    199 KSFTNNEMELEGIYEKINHLNAMSISIQSNFQNPLKTTFVCVCIPEFLSV    248

PF3D7_0415000    251 YETERLIQELTKKNISCYNIVVNQVVFPLDSPNVNLENCKNLLSQIKNEQ    300
                     |||||||||||||||||||||||||||||||..|::.:|:.||.|||::|
PVP01_0521500    249 YETERLIQELTKKNISCYNIVVNQVVFPLDSMTVDVAHCEGLLKQIKDKQ    298

PF3D7_0415000    301 IQSYFNDLISKTEELEDVYISRRKLQSKYLTQIKNLYSNDFHIVCMPQLK    350
                     :|..|:.|:.||:||||||||||||||||||||||||.||||||||||||
PVP01_0521500    299 VQESFSSLVQKTKELEDVYISRRKLQSKYLTQIKNLYGNDFHIVCMPQLK    348

PF3D7_0415000    351 NEIRGLNNISSFSEMLLQSKDIPIYKDNL    379
                     :|||||.|||:||||||:||:||||:
PVP01_0521500    349 SEIRGLQNISNFSEMLLESKEIPIYR---    374
```

**D**

##### PF3D7_0415000 vs. PKNH_0506600

```
PF3D7_0415000      1 MSEDESNSVSCSLSLESDGYSDEEYDTNLNKLIENESLNWIFVGGKGGVG     50
                     ||  :::|:||||:||||.|.:|||||||:||:||::|||||||||||||
PKNH_0506600       1 MS--DADSLSCSLTLESDEYDEEEYDTNLSKLLENKTLNWIFVGGKGGVG     48

PF3D7_0415000     51 KTTTSCSIAVQLSKRRESVLLLSTDPAHNTSDAFNQKFTNQPTLINSFDN    100
                     ||||||||||||:|||||||||||||||||||||||||||||||||||||
PKNH_0506600      49 KTTTSCSIAVQLAKRRESVLLLSTDPAHNTSDAFNQKFTNQPTLINSFDN     98

PF3D7_0415000    101 LYCMEIDTNYSENTAFKLNKKEMFDNILPELLHSFPGIDEALCFAELMQS    150
                     ||||||||.|||||||||||.|.||:|:||||.|||||||||||||||||
PKNH_0506600      99 LYCMEIDTTYSENTAFKLNKTEFFDSIIPELLQSFPGIDEALCFAELMQS    148

PF3D7_0415000    151 IKNMKYSVIVFDTAPTGHTLRLLAFPDLLKKALGYLINIREKLKGTLNVL    200
                     ||||||||||||||||||||||||||:||||||||||::|||||||||:|
PKNH_0506600     149 IKNMKYSVIVFDTAPTGHTLRLLAFPELLKKALGYLISLREKLKGTLNML    198

PF3D7_0415000    201 KNFTNNEMEFDSLYEKINHLNAMSSSIQANFQNPMKTTFVCVCIPEFLSV    250
                     |:|||||:|.:.:|||||||||||.|||:|||||:|||||||||||||||
PKNH_0506600     199 KSFTNNEVELEGIYEKINHLNAMSISIQSNFQNPLKTTFVCVCIPEFLSV    248

PF3D7_0415000    251 YETERLIQELTKKNISCYNIVVNQVVFPLDSPNVNLENCKNLLSQIKNEQ    300
                     ||||||||||||||||||||||||||||||.|.||:.:|:.||.|||:::
PKNH_0506600     249 YETERLIQELTKKNISCYNIVVNQVVFPLDCPTVNVSHCEGLLKQIKDKK    298

PF3D7_0415000    301 IQSYFNDLISKTEELEDVYISRRKLQSKYLTQIKNLYSNDFHIVCMPQLK    350
                     ||..|:.|:.||:||||||||||||||||||||||||.||||||||||||
PKNH_0506600     299 IQESFSSLVQKTKELEDVYISRRKLQSKYLTQIKNLYGNDFHIVCMPQLK    348

PF3D7_0415000    351 NEIRGLNNISSFSEMLLQSKDIPIYKDNL    379
                     :|||||.|||:||||||:|||||||:...
PKNH_0506600     349 SEIRGLENISNFSEMLLESKDIPIYRSEG    377
```

#### Supplementary Figure 1 (contd.)

**E**

**PF3D7_0415000 vs. PCHAS_0726100**

```
PF3D7_0415000      1 MSE--DESNSVSCSLSLESDGYSDEEYDTNLNKLIENESLNWIFVGGKGG     48
                     ||:  .:::|:||||||:||...||.|:||||||||.||||||||||||
PCHAS_0726100      1 MSKAGSDASSISCSLSLDSDSCDDEFYETNLNKLIENTSLNWIFVGGKGG     50

PF3D7_0415000     49 VGKTTTSCSIAVQLSKRRESVLLLSTDPAHNTSDAFNQKFTNQPTLINSF     98
                     |||||||||||:||:|:|||||||||||||||||||||||||:|||||||
PCHAS_0726100     51 VGKTTTSCSIAIQLAKKRESVLLLSTDPAHNTSDAFNQKFTNKPTLINSF    100

PF3D7_0415000     99 DNLYCMEIDTNYSENTAFKLNKKEMFDNILPELLHSFPGIDEALCFAELM    148
                     ||||||||||.:||:||||:||.:..::|:||||.|||||||||||||||
PCHAS_0726100    101 DNLYCMEIDTTFSEDTAFKINKSDFLNSIIPELLQSFPGIDEALCFAELM    150

PF3D7_0415000    149 QSIKNMKYSVIVFDTAPTGHTLRLLAFPDLLKKALGYLINIREKLKGTLN    198
                     |||:||||||||||||||||||||||||||||||||||||::||||||||
PCHAS_0726100    151 QSIRNMKYSVIVFDTAPTGHTLRLLAFPDLLKKALGYLINLKEKLKGTLN    200

PF3D7_0415000    199 VLKNFTNNEMEFDSLYEKINHLNAMSSSIQANFQNPMKTTFVCVCIPEFL    248
                     :|::.|:|||||:.:|:||||||.||.|||.|||||:|||||||||||||
PCHAS_0726100    201 MLQSLTSNEMEFEGMYDKINHLNTMSISIQENFQNPLKTTFVCVCIPEFL    250

PF3D7_0415000    249 SVYETERLIQELTKKNISCYNIVVNQVVFPLDSPNVNLENCKNLLSQIKN    298
                     |||||||||||||||||||||||||||||||.|.:.|:|:|:.||.|||:
PCHAS_0726100    251 SVYETERLIQELTKKNISCYNIVVNQVVFPLTSQDANIESCEGLLKQIKD    300

PF3D7_0415000    299 EQIQSYFNDLISKTEELEDVYISRRKLQSKYLTQIKNLYSNDFHIVCMPQ    348
                     ..|:..|:.||.|.:||||||||||||||||||||||||.|.||||||||
PCHAS_0726100    301 TNIKDSFSSLILKAKELEDVYISRRKLQSKYLTQIKNLYGNYFHIVCMPQ    350

PF3D7_0415000    349 LKNEIRGLNNISSFSEMLLQSKDIPIYKDNL    379
                     ||:|||||:.|:|||||||||||||||...
PCHAS_0726100    351 LKSEIRGLDKIASFSEMLLQSKDIPIYSPQ-    380
```

**F**

**PF3D7_0415000 vs. PY17X_0717200**

```
PF3D7_0415000      1 MSEDES--NSVSCSLSLESDGYSDEEYDTNLNKLIENESLNWIFVGGKGG     48
                     |||..|  :|:||||||:||...||.|:||||||||.||||||||||||
PY17X_0717200      1 MSEGGSDVSSLSCSLSLDSDSCEDEFYETNLNKLIENTSLNWIFVGGKGG     50

PF3D7_0415000     49 VGKTTTSCSIAVQLSKRRESVLLLSTDPAHNTSDAFNQKFTNQPTLINSF     98
                     |||||||||||:||:|:|||||||||||||||||||||||||:|||||||
PY17X_0717200     51 VGKTTTSCSIAIQLAKKRESVLLLSTDPAHNTSDAFNQKFTNKPTLINSF    100

PF3D7_0415000     99 DNLYCMEIDTNYSENTAFKLNKKEMFDNILPELLHSFPGIDEALCFAELM    148
                     ||||||||||.:||:||||:|:....::|:||||.|||||||||||||||
PY17X_0717200    101 DNLYCMEIDTTFSEDTAFKINQSNFLNSIIPELLQSFPGIDEALCFAELM    150

PF3D7_0415000    149 QSIKNMKYSVIVFDTAPTGHTLRLLAFPDLLKKALGYLINIREKLKGTLN    198
                     ||||||||||||||||||||||||||||||||||||||||::||||||||
PY17X_0717200    151 QSIKNMKYSVIVFDTAPTGHTLRLLAFPDLLKKALGYLINLKEKLKGTLN    200

PF3D7_0415000    199 VLKNFTNNEMEFDSLYEKINHLNAMSSSIQANFQNPMKTTFVCVCIPEFL    248
                     :|::.|||||||:.:|:||||||.||.|||.|||||:|||||||||||||
PY17X_0717200    201 MLQSLTNNEMEFEGMYDKINHLNTMSISIQENFQNPLKTTFVCVCIPEFL    250

PF3D7_0415000    249 SVYETERLIQELTKKNISCYNIVVNQVVFPLDSPNVNLENCKNLLSQIKN    298
                     |||||||||||||||||||||||||||||||..|:.|:|.|:|||.|||:
PY17X_0717200    251 SVYETERLIQELTKKNISCYNIVVNQVVFPLICPDANIEKCENLLKQIKD    300

PF3D7_0415000    299 EQIQSYFNDLISKTEELEDVYISRRKLQSKYLTQIKNLYSNDFHIVCMPQ    348
                     ..||..||.||.|.:||||||||||||||||||||||||.|.||||||||
PY17X_0717200    301 TNIQDSFNTLILKAKELEDVYISRRKLQSKYLTQIKNLYGNYFHIVCMPQ    350

PF3D7_0415000    349 LKNEIRGLNNISSFSEMLLQSKDIPIYKDNL    379
                     ||.|||||:.||:|||||||||||||||...
PY17X_0717200    351 LKTEIRGLDKISNFSEMLLQSKDIPIYSPQ-    380
```

**Supplementary Figure 1 (contd.)**

**G**

**PF3D7_0415000 vs. YDL100C**

```
PF3D7_0415000      1 MSEDESNSVSCSLSLESDGYSDEEYDTNLNKLIENESLNWIFVGGKGGVG     50
                                         .|...:.||:.||.:.:..|||||||||||
YDL100C            1 --------------------MDLTVEPNLHSLITSTTHKWIFVGGKGGVG     30

PF3D7_0415000     51 KTTTSCSIAVQ--LSKRRESVLLLSTDPAHNTSDAFNQKFTNQPTLINSF     98
                     |||:|||||:|  ||:..:..||:|||||||.||||.:||......:...
YDL100C           31 KTTSSCSIAIQMALSQPNKQFLLISTDPAHNLSDAFGEKFGKDARKVTGM     80

PF3D7_0415000     99 DNLYCMEIDTNYSENTAFKLNKKEMFD-----------------------    125
                     :||.|||||.:.:        .|:|.|
YDL100C           81 NNLSCMEIDPSAA--------LKDMNDMAVSRANNNGSDGQGDDLGSLLQ    122

PF3D7_0415000    126 -NILPELLHSFPGIDEALCFAELMQSIKNMK------YSVIVFDTAPTGH    168
                      ..|.:|..|.|||||||.|.|:|:.||..:      :..::||||||||
YDL100C          123 GGALADLTGSIPGIDEALSFMEVMKHIKRQEQGEGETFDTVIFDTAPTGH    172

PF3D7_0415000    169 TLRLLAFPDLLKKALGYLINIREKLKGTLNVLKNFTNNEMEFDSLYEKIN    218
                     |||.|..|:.|.|.|.....|..||...||......|.::.     .|:|
YDL100C          173 TLRFLQLPNTLSKLLEKFGEITNKLGPMLNSFMGAGNVDIS-----GKLN    217

PF3D7_0415000    219 HLNAMSSSIQANFQNPMKTTFVCVCIPEFLSVYETERLIQELTKKNISCY    268
                     .|.|...:|:..|.:|..||||||||.||||:||||||||||...::...
YDL100C          218 ELKANVETIRQQFTDPDLTTFVCVCISEFLSLYETERLIQELISYDMDVN    267

PF3D7_0415000    269 NIVVNQVVFPLDSPNVNLENCKNLLSQIKNEQIQSYFNDLISKTEELEDV    318
                     :|:|||::|..:....|.:.|:
YDL100C          268 SIIVNQLLFAENDQEHNCKRCQ----------------------------    289

PF3D7_0415000    319 YISRRKLQSKYLTQIKNLYSNDFHIVCMPQLKNEIRGLNNISSFSEML--    366
                       :|.|:|.|||.||..|| .|||:|.||....|||||||::.||:.|
YDL100C          290 --ARWKMQKKYLDQIDELY-EDFHVVKMPLCAGEIRGLNNLTKFSQFLNK    336

PF3D7_0415000    367 -------------LQSKDIPIYKDNL    379
                                  |:.|:
YDL100C          337 EYNPITDGKVIYELEDKE--------    354
```

**H**

**PF3D7_0415000 vs. SPAC1142.06**

```
PF3D7_0415000        1 MSEDESNSVSCSLSLESDGYSDEEYDTNLNKLIENESLNWIFVGGKGGVG     50
                                           .|.:.....|..|:|..||.|||||||||||
SPAC1142.06          1 -------------------MSFDPLPGTLENLLEQTSLKWIFVGGKGGVG     31

PF3D7_0415000       51 KTTTSCSIAVQLSKRRESVLLLSTDPAHNTSDAFNQKFTNQPTLINSFDN    100
                       |||||||:|:|:||.|.||||:|||||||.||||..||......:..|||
SPAC1142.06         32 KTTTSCSLAIQMSKVRSSVLLISTDPAHNLSDAFGTKFGKDARKVPGFDN     81

PF3D7_0415000      101 LYCMEIDTNYS--ENT--AFKLNKKEMFDNILPELLHSFPGIDEALCFAE    146
                       |..||||.|.|  |.|  |.:.|.......::.:|..:.|||||||.|||
SPAC1142.06         82 LSAMEIDPNLSIQEMTEQADQQNPNNPLSGMMQDLAFTIPGIDEALAFAE    131

PF3D7_0415000      147 LMQSIKNMKYSVIVFDTAPTGHTLRLLAFPDLLKKALGYLINIREKLKGT    196
                       :::.||:|::..::|||||||||||.|.||.:|:||||.|..:.:.....
SPAC1142.06        132 ILKQIKSMEFDCVIFDTAPTGHTLRFLNFPTVLEKALGKLGGLSSRFGPM    181

PF3D7_0415000      197 LNVLKNFTNNEMEFDSLYEKINHLNAMSSSIQANFQNPMKTTFVCVCIPE    246
                       :|.:.:..........|:.|:..:.|..|.:...|:||..||||||||.|
SPAC1142.06        182 INQMGSIMGVNANEQDLFGKMESMRANISEVNKQFKNPDLTTFVCVCISE    231

PF3D7_0415000      247 FLSVYETERLIQELTKKNISCYNIVVNQVVFPLDSPNVNLENCKNLLSQI    296
                       |||:|||||:|||||...|..:||||||::  || ||.....|
SPAC1142.06        232 FLSLYETERMIQELTSYEIDTHNIVVNQLL--LD-PNTTCPQC-------    271

PF3D7_0415000      297 KNEQIQSYFNDLISKTEELEDVYISRRKLQSKYLTQIKNLYSNDFHIVCM    346
                                               ::|||:|.|||.||:.|| .|||:|.:
SPAC1142.06        272 -----------------------MARRKMQQKYLAQIEELY-EDFHVVKV    297

PF3D7_0415000      347 PQLKNEIRGLNNISSFSEMLLQ-------SKDIPIYKDNL    379
                       ||:..|:||..:.||||||::        .|:
SPAC1142.06        298 PQVPAEVRGTEALKSFSEMLVKPYVYPTSGKE--------    329
```

#### Supplementary Figure 1 (contd.)

**I**

**PF3D7_0415000 vs. ENSG00000198356**

```
PF3D7_0415000      1 MSEDESNSVSCSLS---LESDGYSD----EEYDTNLNKLIENESLNWIFV     43
                             ::..::   :|::.:.|    |..:..|:.:||..||.||||
ENSG000001983      1 --------MAAGVAGWGVEAEEFEDAPDVEPLEPTLSNIIEQRSLKWIFV     42

PF3D7_0415000     44 GGKGGVGKTTTSCSIAVQLSKRRESVLLLSTDPAHNTSDAFNQKFTNQPT     93
                     ||||||||||.|||:||||||.|||||::|||||||.||||:|||:..||
ENSG000001983     43 GGKGGVGKTTCSCSLAVQLSKGRESVLIISTDPAHNISDAFDQKFSKVPT     92

PF3D7_0415000     94 LINSFDNLYCMEIDTN----------YSENTAFKLNKKEMFDNILPELLH    133
                     .:..:|||:.||||.:          :.|:....:.||     ::.|.:.
ENSG000001983     93 KVKGYDNLFAMEIDPSLGVAELPDEFFEEDNMLSMGKK-----MMQEAMS    137

PF3D7_0415000    134 SFPGIDEALCFAELMQSIKNMKYSVIVFDTAPTGHTLRLLAFPDLLKKAL    183
                     :|||||||:.:||:|:.:|.|.:||:|||||||||||||||.||.::::.|
ENSG000001983    138 AFPGIDEAMSYAEVMRLVKGMNFSVVVFDTAPTGHTLRLLNFPTIVERGL    187

PF3D7_0415000    184 GYLINIREKLKGTLNVLKNFTN-NEMEFDSLYEKINHLNAMSSSIQANFQ    232
                     |.|:.|:.::...::.:.|... .:|..|.|..|:.....:..|:...|:
ENSG000001983    188 GRLMQIKNQISPFISQMCNMLGLGDMNADQLASKLEETLPVIRSVSEQFK    237

PF3D7_0415000    233 NPMKTTFVCVCIPEFLSVYETERLIQELTKKNISCYNIVVNQVVFPLDSP    282
                     :|.:|||:||||.||||:||||||||||.|..|..:||:|||:|||  .|
ENSG000001983    238 DPEQTTFICVCIAEFLSLYETERLIQELAKCKIDTHNIIVNQLVFP--DP    285

PF3D7_0415000    283 NVNLENCKNLLSQIKNEQIQSYFNDLISKTEELEDVYISRRKLQSKYLTQ    332
                     ....:.|:                              :|.|:|:|||.|
ENSG000001983    286 EKPCKMCE------------------------------ARHKIQAKYLDQ    305

PF3D7_0415000    333 IKNLYSNDFHIVCMPQLKNEIRGLNNISSFSEMLLQSKDIPIYKDNL    379
                     :::|| .|||||.:|.|.:|:||.:.:::||.:||:....|..:
ENSG000001983    306 MEDLY-EDFHIVKLPLLPHEVRGADKVNTFSALLLEPYKPPSAQ---    348
```

**J**

**PF3D7_0415000 vs. TGVEG_231190**

```
PF3D7_0415000      1 MSEDESNSVSCSLSLESDGYSDEEYDTNLNKLIENESLNWIFVGGKGGVG     50
                                       ..|.|.:.:|.:|.|..||.||||||||||
TGVEG_231190       1 ------------------MEDLELEGSLKELFETPSLRWIFVGGKGGVG     31

PF3D7_0415000     51 KTTTSCSIAVQLSKRRESVLLLSTDPAHNTSDAFNQKFTNQPTLINSFDN    100
                     ||||||::|.||:|.|||||::|||||||.||||.|||:|.|||:|.|||
TGVEG_231190      32 KTTTSCAVAAQLAKTRESVLIISTDPAHNISDAFTQKFSNTPTLVNGFDN     81

PF3D7_0415000    101 LYCMEIDTNYSENTAFKLNKKEMFD-------NILPELLHSFPGIDEALC    143
                     ||.||||:.|.|...||::.....:       ::|||:|.:.|||||||.
TGVEG_231190      82 LYAMEIDSRYQETFDFKMSNLPSAEAASFSLTSLLPEMLQAVPGIDEALS    131

PF3D7_0415000    144 FAELMQSIKNMKYSVIVFDTAPTGHTLRLLAFPDLLKKALGYLINIREKL    193
                     ||||||::::|||||||||||||||||||||||||||::.|..|...::|:
TGVEG_231190     132 FAELMQNVQSMKYSVIVFDTAPTGHTLRLLAFPDLLERGLKKLSTFKDKI    181

PF3D7_0415000    194 KGTLNVLKNFTNNEMEFDSLYEKINHLNAMSSSIQANFQNPMKTTFVCVC    243
                     :..|.:|...:..::::......||.:|.|:::|::..||:|..|||||||
TGVEG_231190     182 QSALQMLNAVSGQQIQEQDFAAKIENLKAVTTSVREAFQDPAHTTFVCVC    231

PF3D7_0415000    244 IPEFLSVYETERLIQELTKKNISCYNIVVNQVVFPLDSPNVNLENCK---    290
                     |||||||||||||:|||.|:.|.|.|||||||:||:.  .|..|.|:
TGVEG_231190     232 IPEFLSVYETERLVQELAKQKIDCSNIVVNQVLFPVG--GVQDEGCRPPA    279

PF3D7_0415000    291 NLL-------------------------SQIKNEQIQSYFNDLISKTEE    314
                     :||                          :|.:|.:::.....:..:...
TGVEG_231190     280 SLLASADAETPAPLEELLAPPAARGEKETAQEENARLRQLIRRMQIRLLA    329

PF3D7_0415000    315 LEDVYISRRKLQSKYLTQIKNLYSNDFHIVCMPQLKNEIRGLNNISSFSE    364
                     ||..|.|||.:||:||.||::|||.|||:|.:||...|:||:..:..|.:
TGVEG_231190     330 LEKSYHSRRAMQSRYLQQIQDLYSFDFHVVPIPQQPEEVRGIERLLRFGD    379

PF3D7_0415000    365 MLLQSKDIPIYKDNL-------    379
                     :|...:.:||    |
TGVEG_231190     380 LLSSCRPLPI----LPPAPSSP    397
```

**Supplementary Figure 1 (contd.)**

**K**

**PF3D7_0415000 vs. LDBPK_110710**

```
PF3D7_0415000      1 MSEDESNSVSCSLSLESDGYSDEEYDTNLNKLIENESLNWIFVGGKGGVG     50
                                             .|..|.:|: :.:|.|||||||||||
LDBPK_110710       1 ------------------------MDPTLTELL-HANLEWIFVGGKGGVG     25

PF3D7_0415000     51 KTTTSCSIAVQLS------------KRRESVLLLSTDPAHNTSDAFNQKF     88
                     ||||||::|...:            .|...|||:|||||||.||||||:|
LDBPK_110710      26 KTTTSCALATLFATTPISDAASPGGTRPRRVLLISTDPAHNLSDAFNQRF     75

PF3D7_0415000     89 TNQPTLINSF-DNLYCMEID-TNYS-------------ENTAFKLNKKEM    123
                     ...||.:... ::|..||:| .|::             :.:|..|.::..
LDBPK_110710      76 GPHPTPVKGLEESLAAMEVDPKNFTHGALMSSLTGTKRDGSASSLPEEAA    125

PF3D7_0415000    124 FD------------NILPELLHSFPGIDEALCFAELMQSIKNMKYSVIVF    161
                     .|            .:|.|...:.|||||...|||::..::.:.|.:::|
LDBPK_110710     126 ADAAQHTTTFARIGAVLKEAARTMPGIDEISVFAEILHYVRTLSYDLLIF    175

PF3D7_0415000    162 DTAPTGHTLRLLAFPDLLKKALGYLINIREKLKGTLNVLKNFTNNEM---    208
                     |||||||||||||.|..|......|::: |.|...:....:...:.:
LDBPK_110710     176 DTAPTGHTLRLLALPQTLNSTFDKLMSL-EGLAPMIEAASHLIGSNLGDL    224

PF3D7_0415000    209 -----------------------------------------EFDSLYEKI    217
                                                              ..|.:....
LDBPK_110710     225 GGACGDTAGSCEQATAAPSPSSAAPGAGSAAAASPQSRWCITADEVRSTA    274

PF3D7_0415000    218 NHLNAMSSSIQANFQNPMKTTFVCVCIPEFLSVYETERLIQELTKKNISC    267
                     .|.......:||.|.:|.:|:||||||.|||||||||||:|||.|.||.|
LDBPK_110710     275 LHWRQTMEEVQARFNDPNRTSFVCVCIAEFLSVYETERLVQELMKYNIGC    324

PF3D7_0415000    268 YNIVVNQVVF-PLDSPNVNLENCKNLLSQIKNEQIQSYFNDLISKTEELE    316
                     .:|||||:|. |...|...: |.
LDBPK_110710     325 DSIVVNQLVLKPSSEPPCRM--CS-------------------------    346

PF3D7_0415000    317 DVYISRRKLQSKYLTQIKNLYSNDFHIVCMPQLKNEIRGLNNISSFSEML    366
                         :|:|:|:|||.||..|| .|||:|.||.|.:|:||:..:..|:..|
LDBPK_110710     347 ----ARQKIQAKYLEQIDLLY-EDFHVVKMPLLSDEVRGVPALKKFARFL    391

PF3D7_0415000    367 LQ--SKDIPIYKDNL---    379
                     .:  |.:...|.|..
LDBPK_110710     392 QEPYSPETHGYIDVQEPC   409
```

**L**

**PF3D7_0415000 vs. Tc00.1047053507763.30**

```
PF3D7_0415000      1 MSEDESNSVSCSLSLESDGYSDEEYDTNLNKLIENESLNWIFVGGKGGVG     50
                                 :|||          ..|..|:.:: |.|||||||||||
Tc00.10470535      1 ------------MSLE----------PTLRDLLHSK-LQWIFVGGKGGVG     27

PF3D7_0415000     51 KTTTSCSI-----------AVQLSKRRESVLLLSTDPAHNTSDAFNQKFT     89
                     ||||||::           ||..:.|...|||:|||||||.||||:|||.
Tc00.10470535     28 KTTTSCALATLFASTPVHDAVTNTTRPRRVLLISTDPAHNLSDAFSQKFG     77

PF3D7_0415000     90 NQPTLINSF-DNLYCMEID-TNYSE-----------NTAFKLNKKEMF--    124
                     ..|..:|.. :.|:.||:| |.::.           :.|...:....|
Tc00.10470535     78 KTPVPVNGMEETLFAMEVDPTTFTHGGFGAMLGFPGHIATDADAPSPFAA    127

PF3D7_0415000    125 -DNILPELLHSFPGIDEALCFAELMQSIKNMKYSVIVFDTAPTGHTLRLL    173
                      .|||.|...:.|||||...|||:::.::.:.|.|::|||||||||||||
Tc00.10470535    128 LGNILKEAAGTLPGIDELSVFAEILRGVQQLSYDVVIFDTAPTGHTLRLL    177

PF3D7_0415000    174 AFPDLLKKALGYLINIR--EKLKGTLNVLKNFTNNEMEFDSLYEKINHLN    221
                     |.|..|...:..|:::.  ..|....:.:.:.|.|..:..||........
Tc00.10470535    178 ALPHTLNSTMEKLLSVEGLNTLIQAASAVLSSTTNLGDMSSLMPAFKQWR    227

PF3D7_0415000    222 AMSSSIQANFQNPMKTTFVCVCIPEFLSVYETERLIQELTKKNISCYNIV    271
                     .....:|..|.:..||.|:||||||||||||||||:|||.|.:|||.:||
Tc00.10470535    228 ENVQEVQRQFTDAEKTAFICVCIPEFLSVYETERLVQELMKYDISCDSIV    277

PF3D7_0415000    272 VNQVVF-PLDSPNVNLENCKNLLSQIKNEQIQSYFNDLISKTEELEDVYI    320
                     |||:|. |...|:..:.|
Tc00.10470535    278 VNQLVLKPSSEPDCRMCN--------------------------------    295

PF3D7_0415000    321 SRRKLQSKYLTQIKNLYSNDFHIVCMPQLKNEIRGLNNISSFSEMLLQSK    370
                     :|:|:|||||.||.:|| .|||:|.||.|.:|:||:..:..|::.||:
Tc00.10470535    296 ARQKIQSKYLAQIDSLY-EDFHVVKMPLLSDEVRGVPALQRFAQFLLE--    342

PF3D7_0415000    371 DIPIYKDNL----------    379
                       |...|..
Tc00.10470535    343 --PYDADRHGYIDVCGAAS    359
```

#### Supplementary Figure 1 (contd.)

**M**

**PF3D7_0415000 vs. At1g01910**

```
PF3D7_0415000      1 MSEDESNSVSCSLSLESDGYSDEEYDTNLNKLIENESLNWIFVGGKGGVG     50
                                        .:.:..:..:..:::.|||.|:|||||||||
At1g01910          1 -------------------MAADLPEATVQNILDQESLKWVFVGGKGGVG     31

PF3D7_0415000     51 KTTTSCSIAVQLSKRRESVLLLSTDPAHNTSDAFNQKFTNQPTLINSFDN    100
                     |||.|..:|:.|:..|.|||::|||||||.||||.|:||..|||:..|.|
At1g01910         32 KTTCSSILAICLASVRSSVLIISTDPAHNLSDAFQQRFTKSPTLVQGFSN     81

PF3D7_0415000    101 LYCMEIDTNYSENTAFKLNKKEMFDNILPELLHSFPGIDEALCFAELMQS    150
                     |:.||:|.....:   .:...:..|.:..:|.::.||||||:.|||:::.
At1g01910         82 LFAMEVDPTVETD---DMAGTDGMDGLFSDLANAIPGIDEAMSFAEMLKL    128

PF3D7_0415000    151 IKNMKYSVIVFDTAPTGHTLRLLAFPDLLKKALGYLINIREKLKGTLNVL    200
                     ::.|.|:.|||||||||||||||.||..|:|.|..|::::.:..|.:..:
At1g01910        129 VQTMDYATIVFDTAPTGHTLRLLQFPATLEKGLSKLMSLKSRFGGLMTQM    178

PF3D7_0415000    201 KNFTNNEMEF--DSLYEKINHLNAMSSSIQANFQNPMKTTFVCVCIPEFL    248
                     ......|.||  |:|..::..|..:...:...|::|..||||||||||||
At1g01910        179 SRMFGMEDEFGEDALLGRLEGLKDVIEQVNRQFKDPDMTTFVCVCIPEFL    228

PF3D7_0415000    249 SVYETERLIQELTKKNISCYNIVVNQVVFPLDSPNVNLENCKNLLSQIKN    298
                     |:||||||:|||.|..|..:||::|||::                   .:
At1g01910        229 SLYETERLVQELAKFEIDTHNIIINQVLY-------------------DD    259

PF3D7_0415000    299 EQIQSYFNDLISKTEELEDVYISRRKLQSKYLTQIKNLYSNDFHIVCMPQ    348
                     |.::|             .:..:|.::|.|||.|...|| :||:|..:|.
At1g01910        260 EDVES-------------KLLRARMRMQQKYLDQFYMLY-DDFNITKLPL    295

PF3D7_0415000    349 LKNEIRGLNNISSFSEMLLQSKDIPIYKDNL-------------------    379
                     |..|:.|:..:.:||...|........:.|:
At1g01910        296 LPEEVTGVEALKAFSHKFLTPYHPTTSRSNVEELERKVHTLRLQLKTAEE    345

PF3D7_0415000    380 --------    379

At1g01910        346 ELERVKSG    353
```

**N**

**PF3D7_0415000 vs. ArsA**

```
PF3D7_0415000      1 MSEDESNSVSCSLSLESDGYSDEEYDTNLNKLIENESLNWIFVGGKGGVG     50
                                        .|.:....:::.::.:.:...:...|.||||
ArsA               1 -------------------MSAKPPALDMHAILSDTANRVVVCCGAGGVG     31

PF3D7_0415000     51 KTTTSCSIAVQLSKRRESVLLLSTDPAHNTSDAFNQK-FTNQPTLI----     95
                     ||||:.::|::.::...:|::|:.|||...:.|...| ..|.|..:
ArsA              32 KTTTAAAMALRAAEYGRTVVVLTIDPAKRLAQALGIKDLGNTPQRVPLAP     81

PF3D7_0415000     96 NSFDNLYCMEID---------TNYSE-NTAFKLNKKEMFDNILPELLHSF    135
                     .....|:.|.:|         ..||: ..|..:.:.:.:..:..    |.
ArsA              82 EVTGELHAMMLDMRRTFDEMVVQYSDPGRADAILENQFYQTVAT----SL    127

PF3D7_0415000    136 PGIDEALCFAELMQSIKNMKYSVIVFDTAPTGHTLRLLAFPDLLKKALGY    185
                     .|..|.:...:|.|.:...|:.::|.||.|:.:.|..|..|    |.||.
ArsA             128 AGTQEYMAMEKLGQLLAEDKWDLVVVDTPPSRNALDFLDAP----KRLGS    173

PF3D7_0415000    186 LINIR-------------EKLKGTLNVLKNFTNNEMEFDSLYEKINHLNA    222
                     .::.|             :.:.|.:.:.....:.:....|.:.....:.|
ArsA             174 FMDSRLWRLLLAPGRGIGKLVTGAVGLAMKALSTVLGSQMLSDAAGFVQA    223

PF3D7_0415000    223 MSSSIQANFQNPMK---------TTFVCVCIPEFLSVYETERLIQELTKK    263
                     :.::.....|...|         |.||.|...|..::.|....:..|:.:
ArsA             224 LDATFDGFRQKADKTYELLKRRGTQFVVVSAAEPDALREASFFVDRLSNE    273

PF3D7_0415000    264 NISCYNIVVNQVVFPLDSPNVNLENCKNLLSQIKNEQIQSYFNDLISKTE    313
                     ::....:::|:.     .|.         ||.:..|:.:...::|.::..
ArsA             274 HMPLAGLILNRT-----HPT---------LSDLHAEKAEEAADELAAEDP    309

PF3D7_0415000    314 E-----LEDVYISRRKLQSKYLTQIKNLYSNDFH--IVCMPQLKNEIRGL    356
                     :     :..::..|.....:.:..:......:.|  ||.:|.|..::..|
ArsA             310 DSLAAAVLRIHADRAHTAKREVRLLSRFTGANPHVAIVGVPSLPFDVSDL    359

PF3D7_0415000    357 NNISSFSEMLL-QSKDIPIYKDNL    379
                     :.:.:.::.:. ::.|..
ArsA             360 DALRAIADQITGEAADAA------    377
```

### Supplementary Figure 1 (contd.)

**O**

**PF3D7_0415000 vs. cgd7_4070**

```
PF3D7_0415000      1 MSEDESNSVSCSLSLESDGYSDEEYD--TNLNKLIENESLNWIFVGGKGG     48
                                    .|..|.|.:.|  .:|..|...::|.|||||||||
cgd7_4070          1 ---------------MSTAYFDADCDLEPSLKSLFSLKTLKWIFVGGKGG     35

PF3D7_0415000     49 VGKTTTSCSIAVQLSKRRESVLLLSTDPAHNTSDAFNQKFTNQPTLINSF     98
                     |||||||||||.:|::.|||||:||||||||.||||.|||:|.|||:|.:
cgd7_4070         36 VGKTTTSCSIASRLAEERESVLILSTDPAHNLSDAFVQKFSNAPTLVNGY     85

PF3D7_0415000     99 DNLYCMEIDTNYSENTAFKLNKK-EMFDNILPELLHSFPGIDEALCFAEL    147
                     .|||.||:|.:|.:...|||.:: .:|...||:|:.:.|||||||.||.|
cgd7_4070         86 KNLYAMELDASYQQAVEFKLKEENSLFSKFLPDLISALPGIDEALGFATL    135

PF3D7_0415000    148 MQSIKNMKYSVIVFDTAPTGHTLRLLAFPDLLKKALGYLINIREKLKGTL    197
                     |||:|:|.||||||||||||||||||:||.||:|.|..|.:|::.:.|.|
cgd7_4070        136 MQSVKSMSYSVIVFDTAPTGHTLRLLSFPSLLEKGLSKLFSIKQNMSGAL    185

PF3D7_0415000    198 NVLKNFTNNEMEFDSLYEKINHLNAMSSSIQANFQNPMKTTFVCVCIPEF    247
                     .::.:.:.|.:|.::|..|:..|.|:::|::..||:|.||||||||||||
cgd7_4070        186 QLINSVSGNAIEEETLNSKLEDLKAITTSVKETFQDPSKTTFVCVCIPEF    235

PF3D7_0415000    248 LSVYETERLIQELTKKNISCYNIVVNQVVFPLDSPNVN------LENCKN    291
                     |||||||||||||.|::|||.:||||||:||:|.|:.|      |::...
cgd7_4070        236 LSVYETERLIQELAKQSISCSHIVVNQVMFPIDLPSGNDQGESVLKDSSE    285

PF3D7_0415000    292 LL---------SQIK--NEQIQSYFNDLISKTEELEDVYISRRKLQSKYL    330
                     ||         |::.  .|:|...:|.|:|.::.|...|.|:|.:|.|||
cgd7_4070        286 LLKLEDIPSDHSKLVEFTEKIVCSYNKLLSYSKLLYSKYYSKRNMQMKYL    335

PF3D7_0415000    331 TQIKNLYSNDFHIVCMPQLKNEIRGLNNISSFSEMLLQSKDIPIYKDNL    379
                     .||::|||.|||:..:|.|.||:..:..:.|
cgd7_4070        336 EQIRDLYSYDFHVAYIPTLNNEVSKIRVLIS------------------    366
```
