## Supplementary Figure 2 for "A conserved Guided Entry of Tail-anchored pathway is involved in the trafficking of tail-anchored membrane proteins in *Plasmodium falciparum*"

**A**

```
                                                                                P-loop
ArsA                ----------------------------MTVGAPPRGYTQRVRVILFTGKGGVGKTSVAA     32
PF3D7_0415000       MSEDESNSVSCSLSLES----DGYSDEEYDTNLNKLIENESLNWIFVGGKGGVGKTTTSC     56
ScGet3              ------------------------MDLTVEPNLHSLITSTTHKWIFVGGKGGVGKTTSSC     36
HsGet3              MA-----AGVAGWGVEAEEFEDAPDVEPLEPTLSNIIEQRSLKWIFVGGKGGVGKTTCSC     55
                                                              . *:. ********: :.

                                         Switch I motif
ArsA                ATAVRC--ARAGYRTVIMSTDPAHSLGDSFDVELGSDLTPIS--DNLWAHEVSSLHEMQR     88
PF3D7_0415000       SIAVQL--SKRRESVLLLSTDPAHNTSDAFNQKFTNQPTLINSFDNLYCMEIDTNYSENT     114
ScGet3              SIAIQMALSQPNKQFLLISTDPAHNLSDAFGEKFGKDARKVTGMNNLSCMEIDPSAALKD     96
HsGet3              SLAVQL--SKGRESVLIISTDPAHNISDAFDQKFSKVPTKVKGYDNLFAMEIDPSLGVAE     113
                    : *::   ::     :::******. .*:*. :: .    :.  :** . *:.

                                                •   •   •• •
ArsA                ---HWVKLHEY-----AVEVFATQGLDEVVADEVANPPGMDEIASLMWIKHYAQ------     134
PF3D7_0415000       AFKL--N--------------KKEMFDNILPELLHSFPGIDEALCFAELMQSIK------     152
ScGet3              MNDMAVSRANNNGSDGQGDDLGSLLQGGALADLTGSIPGIDEALSFMEVMKHIKRQEQGE     156
HsGet3              LPDEFFEEDNML-----------SMGKKMMQEAMSAFPGIDEAMSYAEVMRLVK------     156
                                                 : :     **:**  .   : :  :

                       B-motif  Switch II motif
                                             •   ••  •  •   •  •••   ••       •
ArsA                RAEHDVLIVDCAPTGETLQLLTFPDAAKWWLDKIYPWERRAMKVARPVLQPMMG-IPLPS     193
PF3D7_0415000       NMKYSVIVFDTAPTGHTLRLLAFPDLLKKALGYLINIREKLKGTL-NVLKNF-TNNEMEF     210
ScGet3              GETFDTVIFDTAPTGHTLRFLQLPNTLSKLLEKFGEITNKLGPML-NSF---MGAGNVD-     211
HsGet3              GMNFSVVVFDTAPTGHTLRLLNFPTIVERGLGRLMQIKNQISPFI-SQMCNMLGLGDMNA     215
                       ...::.* ****.**::* :*   .  *  :    .:        :        :

ArsA                DEVYASLKDLLLDLGGMRKVLTDPATTTVRIVLNLEKMVVKEAKRAYTYLSLFGYLTDAV     253
PF3D7_0415000       DSLYEKINHLNAMSSSIQANFQNPMKTTFVCVCIPEFLSVYETERLIQELTKKNISCYNI     270
ScGet3              --ISGKLNELKANVETIRQQFTDPDLTTFVCVCISEFLSLYETERLIQELISYDMDVNSI     269
HsGet3              DQLASKLEETLPVIRSVSEQFKDPEQTTFICVCIAEFLSLYETERLIQELAKCKIDTHNI     275
                      :  .::.       :   : :*  **.  *   * : : *::*    *         :

                                CxxC motif (eukaryotes)
ArsA                VVNRLLPSE-----------------------------------LHDELFQRWQRIHKRY     278
PF3D7_0415000       VVNQVVFPLDSPNVNLENCKNLLSQIKNEQIQSYFNDLISKTEELEDVYISRR-KLQSKY     329
ScGet3              IVNQLLFAENDQEHNCKRCQ------------------------------ARW-KMQKKY     298
HsGet3              IVNQLVFPD--PEKPCKMCE------------------------------ARH-KIQAKY     302
                    :**:::                                             *  ::: :*

                                          A-loop
ArsA                QVEVEQSFA-GIPIFNVPLFDREVVGESMLSRMAEETYGDRDPAQHFATASPQRIDKEGA     337
PF3D7_0415000       LTQIKNLYSNDFHIVCMPQLKNEIRGLNNISSFSEMLLQSKDIPIYKDNL----------     379
ScGet3              LDQIDELYE-DFHVVKMPLCAGEIRGLNNLTKFSQFLNKEYNPITDGKVIY-ELEDKE--     354
HsGet3              LDQMEDLYE-DFHIVKLPLLPHEVRGADKVNTFSALLLEPYKPPSAQ-------------     348
                      ::.: :  .: :. :*    *: * . :.  ::       .

ArsA                DYVLALKVPFADRSSVDLSRHNGELFVTVGNYRREIALPRVLAQRDTSGATIHDGELRVR     397
PF3D7_0415000       ------------------------------------------------------------     379
ScGet3              ------------------------------------------------------------     354
HsGet3              ------------------------------------------------------------     348

ArsA                FTRKEARAGAGGSAPSPDLAPRGKR          422
PF3D7_0415000       -------------------------          379
ScGet3              -------------------------          354
HsGet3              -------------------------          348
```

B

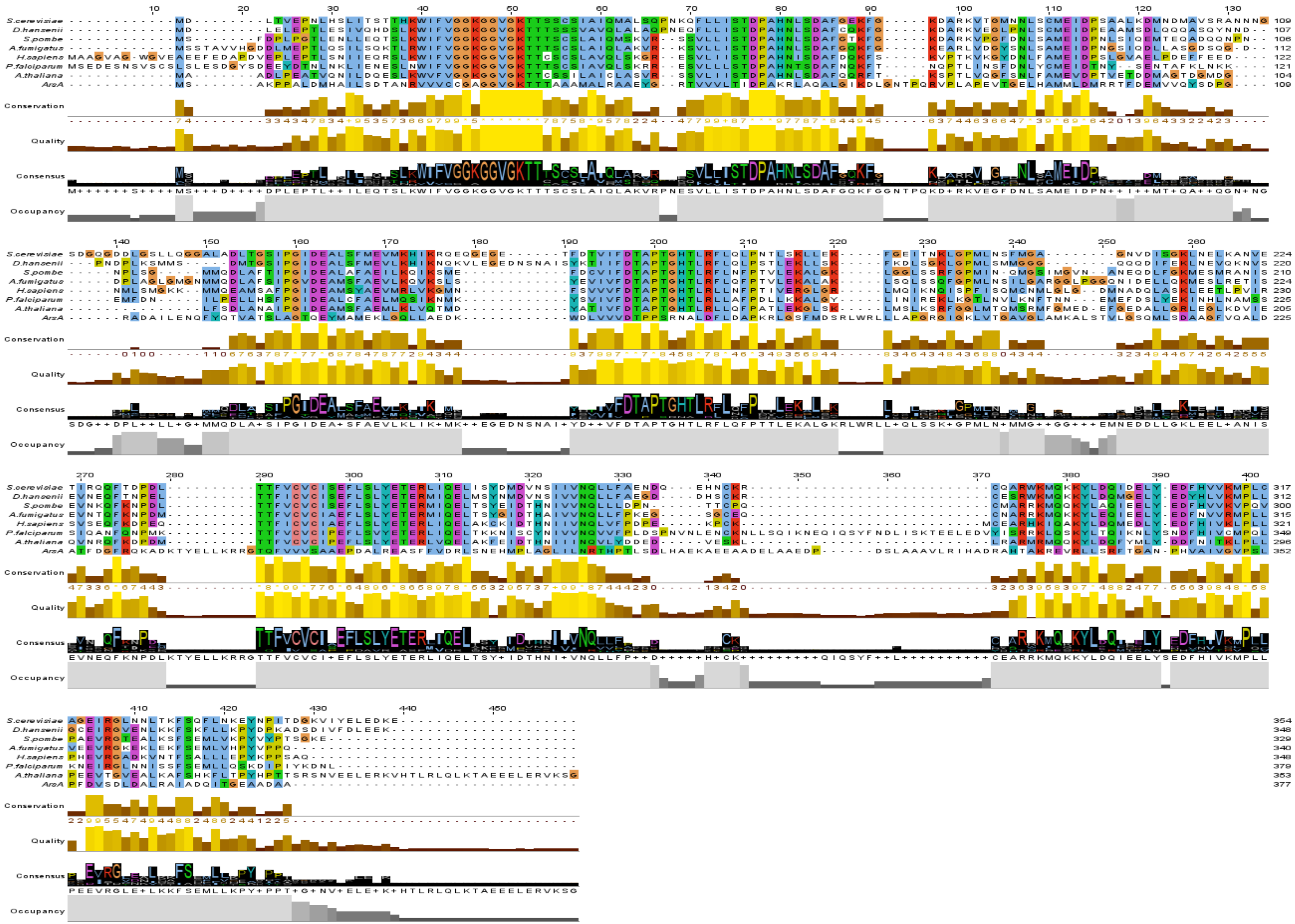
S.cerevisiae
D.hansenii
S.pombe
A.fumigatus
H.sapiens
P.falciparum
A.thaliana
ArsA
Conservation
Quality
Consensus
Occupancy
