## Supplementary figures and images for "A conserved Guided Entry of Tail-anchored pathway is involved in the trafficking of tail-anchored membrane proteins in *Plasmodium falciparum*"

### Supplementary Figure 3

**Supplementary Figure 3**

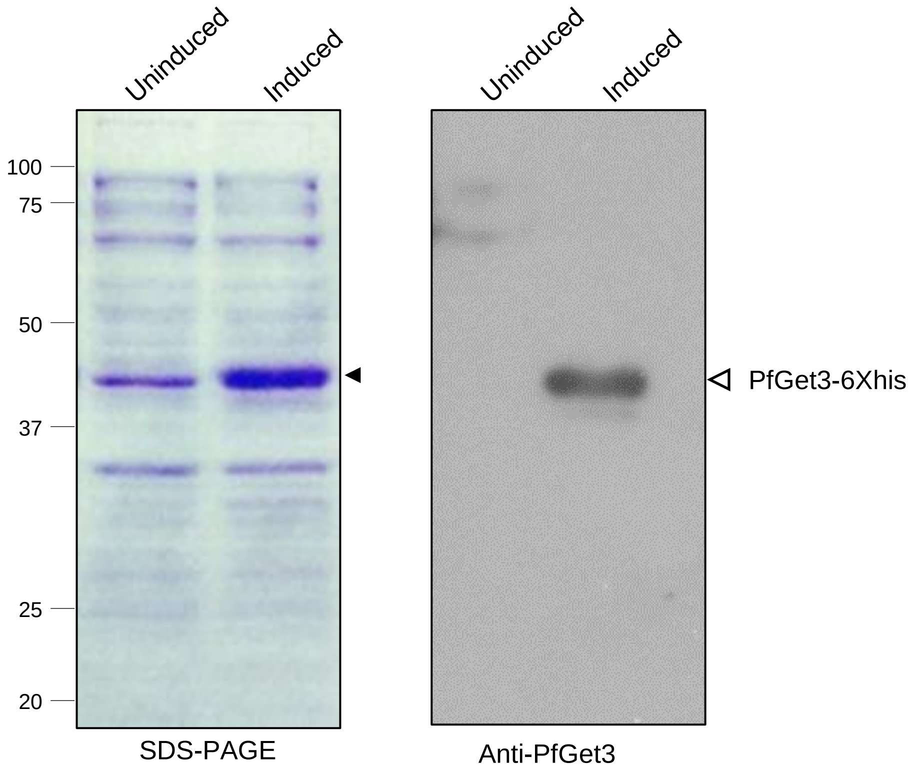

### Supplementary Figure 6

**Supplementary Figure 6**

ScGet3 PfGet3

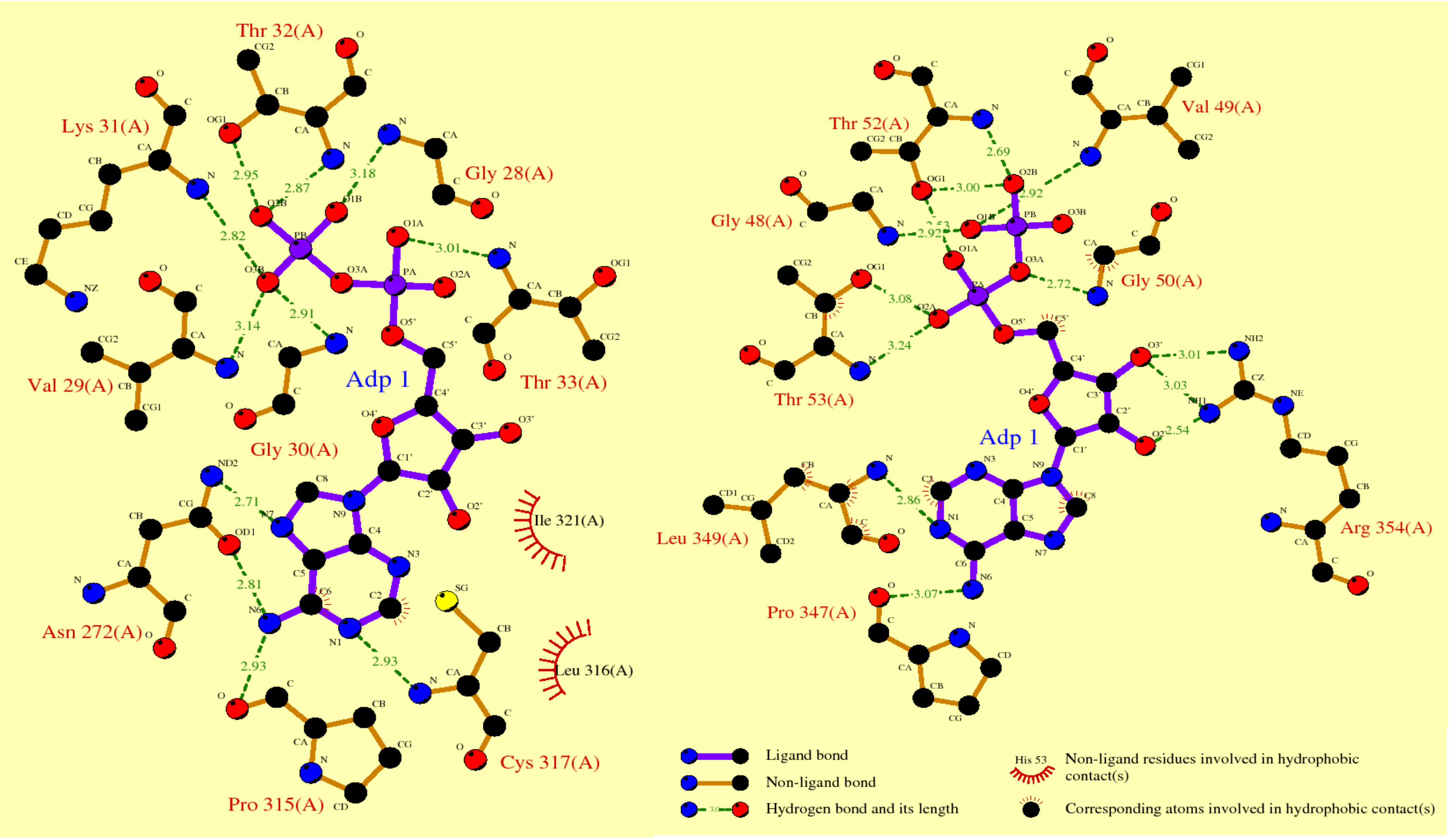

### Supplementary Figure 10

#### Supplementary Figure 10

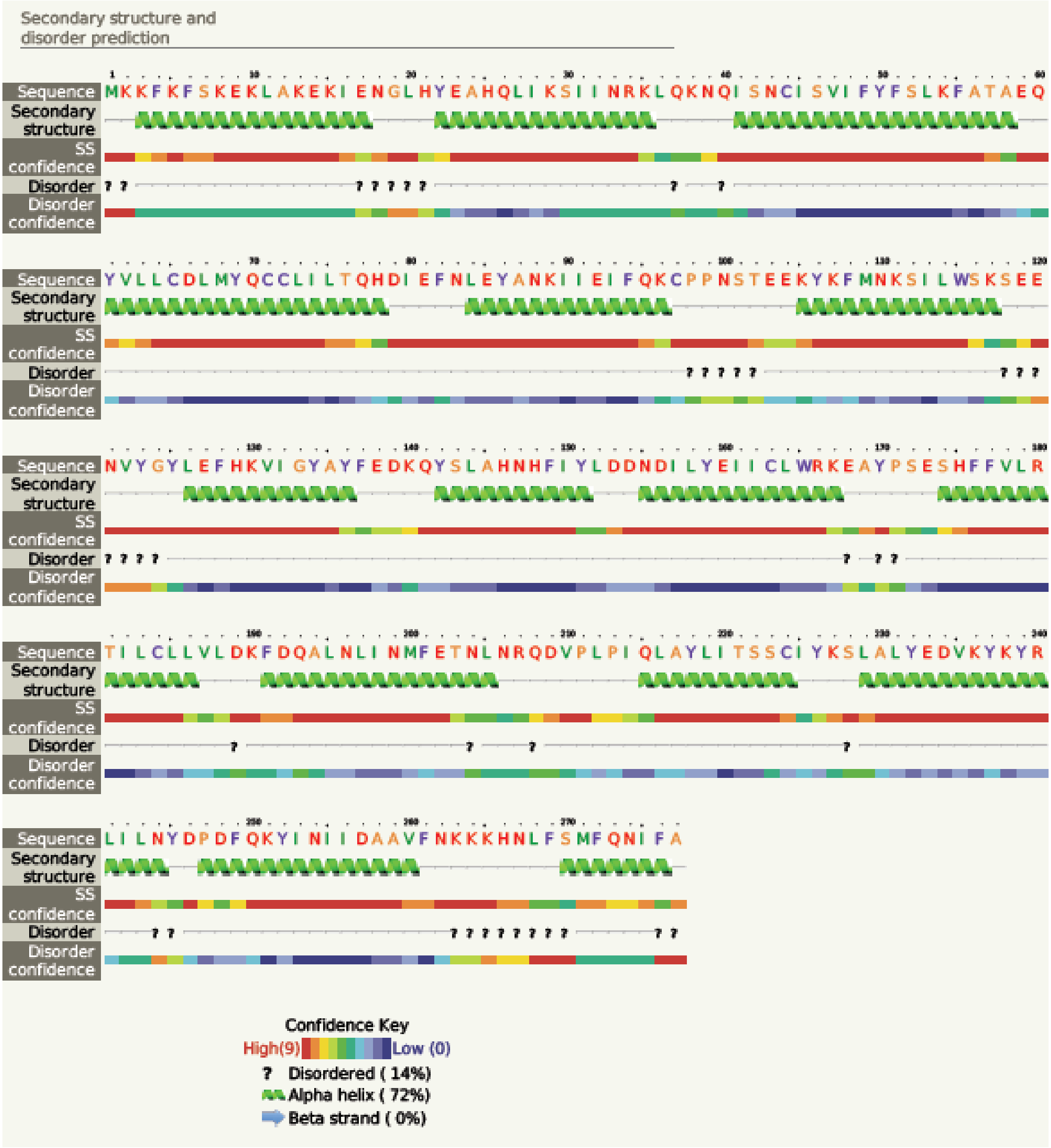

### Supplementary Figure 12

**Supplementary Figure 12**

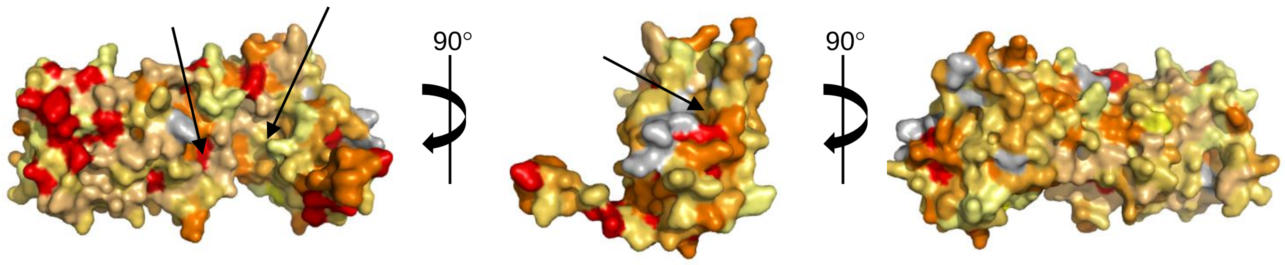
