## Supplementary Figure 4 for "A conserved Guided Entry of Tail-anchored pathway is involved in the trafficking of tail-anchored membrane proteins in *Plasmodium falciparum*"

**A**

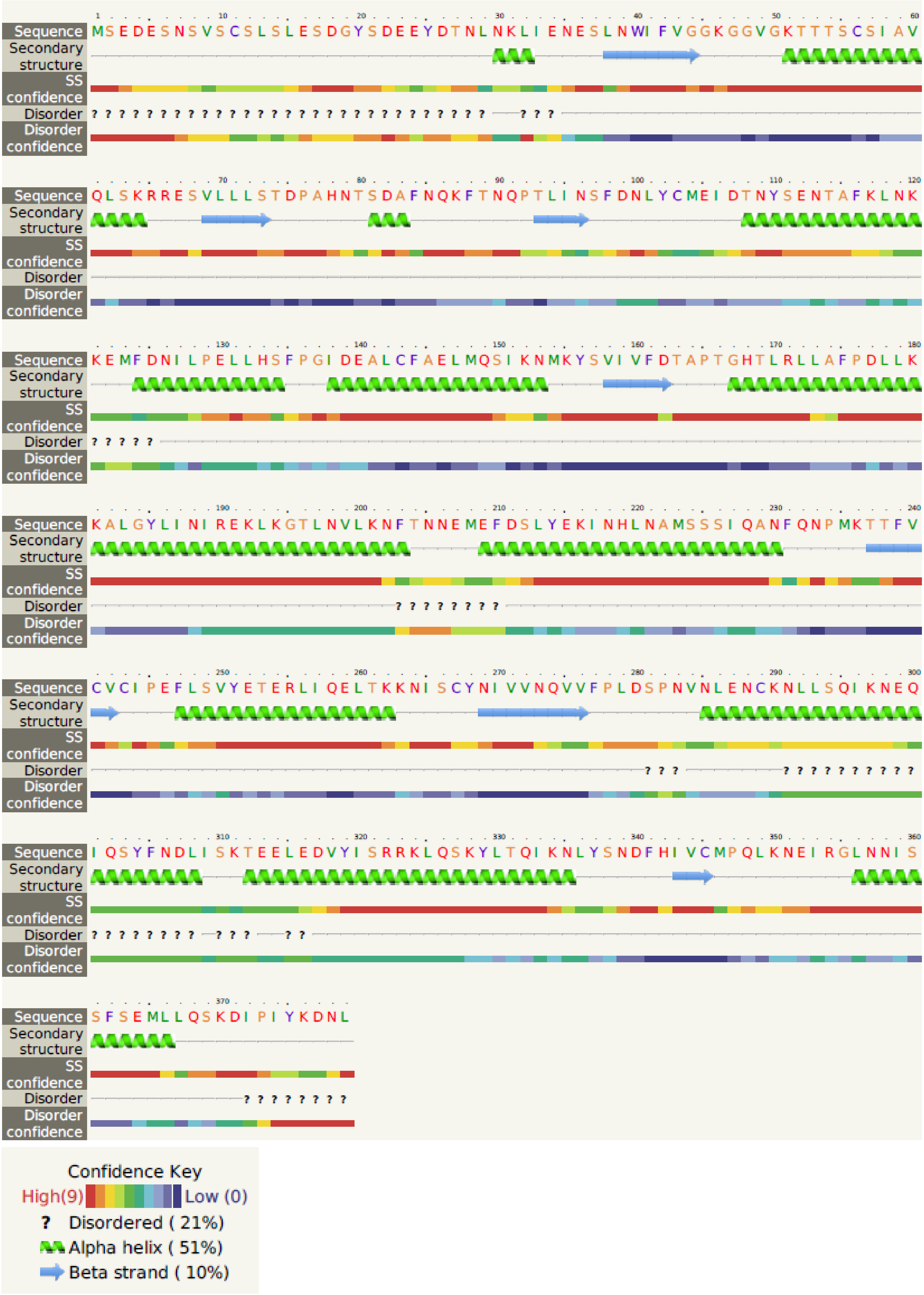

**B**

### PfGet3 Length: 379
### PfGet3 Number of predicted TMHs: 0
### PfGet3 Exp number of AAs in TMHs: 0.04026
### PfGet3 Exp number, first 60 AAs: 0.00986
### PfGet3 Total prob of N-in: 0.01042

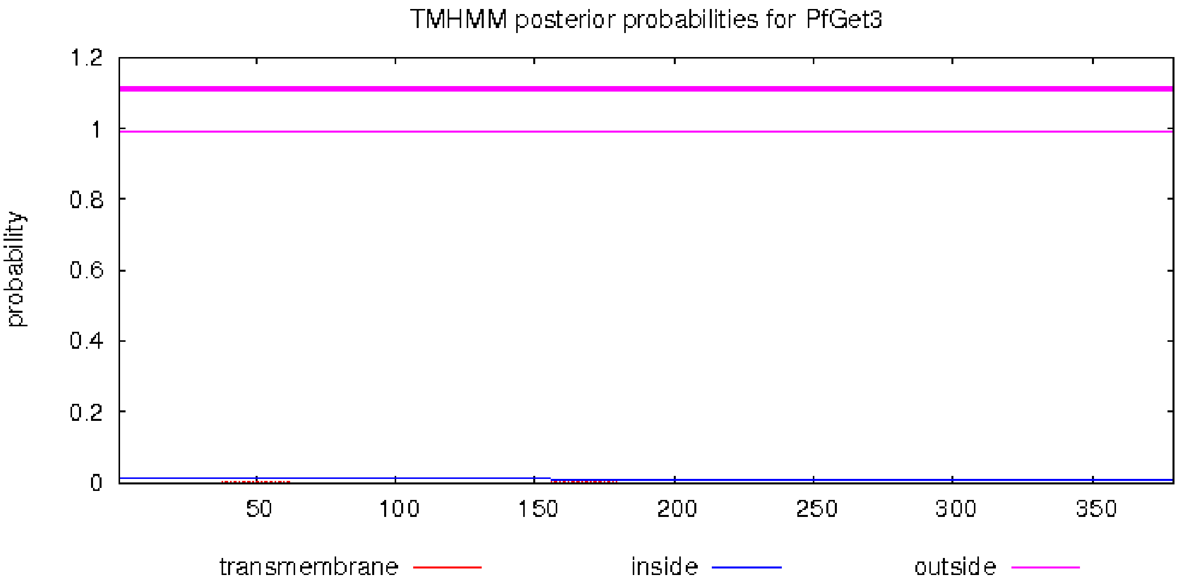
