## Supplementary Figure 5 for "A conserved Guided Entry of Tail-anchored pathway is involved in the trafficking of tail-anchored membrane proteins in *Plasmodium falciparum*"

Supplementary Figure 5. Hit report of PfGet3 predicted by Phyre2 server (www.sbg.bio.ic.ac.uk/phyre2). Schematic tabular representation showing the % confidence and identity of PfGet3 with the retrieved hits from Phyre2 server.

| # | Template | Alignment Coverage | 3D Model | Confidence | % i.d. | Template Information |
|---|---|---|---|---|---|---|
| 1 | c5zmfA_ | Alignment | 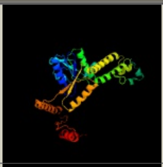 | 100.0 | 29 | **PDB header:**hydrolase/transport protein **Chain:** A: **PDB Molecule:**atpase arsa1; **PDBTitle:** amppnp complex of c. reinhardtii arsa1 |
| 2 | c6bs5B_ | Alignment | 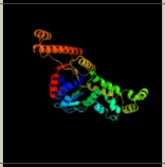 | 100.0 | 17 | **PDB header:**unknown function **Chain:** B: **PDB Molecule:**anion transporter; **PDBTitle:** crystal structure of amp-pnp-bound bacterial get3-like a and b in2 mycobacterium tuberculosis |
| 3 | c1ii0A_ | Alignment | 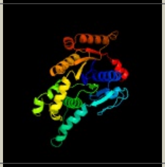 | 100.0 | 24 | **PDB header:**hydrolase **Chain:** A: **PDB Molecule:**arsenical pump-driving atpase; **PDBTitle:** crystal structure of the escherichia coli arsenite-translocating2 atpase |
| 4 | c6so5A_ | Alignment | 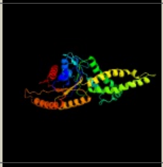 | 100.0 | 50 | **PDB header:**membrane protein **Chain:** A: **PDB Molecule:**atpase asna1; **PDBTitle:** homo sapiens wrb/caml heterotetramer in complex with a trc40 dimer |
| 5 | c3ug7D_ | Alignment | 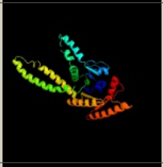 | 100.0 | 33 | **PDB header:**hydrolase **Chain:** D: **PDB Molecule:**arsenical pump-driving atpase; **PDBTitle:** crystal structure of get3 from methanocaldococcus jannaschii |
| 6 | c3zq6D_ | Alignment | 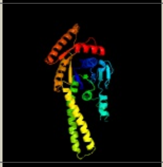 | 100.0 | 34 | **PDB header:**hydrolase **Chain:** D: **PDB Molecule:**putative arsenical pump-driving atpase; **PDBTitle:** adp-alf4 complex of m. therm. trc40 |
| 7 | c2wooC_ | Alignment | 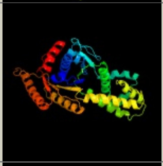 | 100.0 | 51 | **PDB header:**hydrolase **Chain:** C: **PDB Molecule:**atpase get3; **PDBTitle:** nucleotide-free form of s. pombe get3 |
| 8 | c3igfB_ | Alignment | 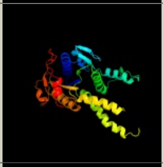 | 100.0 | 20 | **PDB header:**atp binding protein **Chain:** B: **PDB Molecule:**all4481 protein; **PDBTitle:** crystal structure of the all4481 protein from nostoc sp. pcc 7120,2 northeast structural genomics consortium target nsr300 |
| 9 | c3ibgF_ | Alignment | 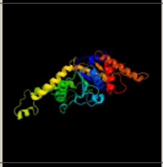 | 100.0 | 50 | **PDB header:**hydrolase **Chain:** F: **PDB Molecule:**atpase, subunit of the get complex; **PDBTitle:** crystal structure of aspergillus fumigatus get3 with bound2 adp |
| 10 | c2wojD_ | Alignment | 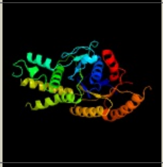 | 100.0 | 51 | **PDB header:**hydrolase **Chain:** D: **PDB Molecule:**atpase get3; **PDBTitle:** adp-alf4 complex of s. cerevisiae get3 |
| 11 | c5bwkA_ | Alignment | 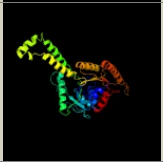 | 100.0 | 46 | **PDB header:**hydrolase/transport **Chain:** A: **PDB Molecule:**atpase get3; **PDBTitle:** 6.0 a crystal structure of a get3-get4-get5 intermediate complex from2 s.cerevisiae |

| | | | | | | |
|---|---|---|---|---|---|---|
| 12 | c6bs3A_ | Alignment | 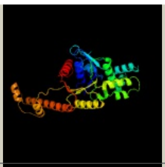 | 100.0 | 19 | **PDB header:**unknown function<br>**Chain:** A: **PDB Molecule:**putative atpase rv3679;<br>**PDBTitle:** crystal structure of adp-bound bacterial get3-like a and b in2 mycobacterium tuberculosis |
| 13 | d1ihua1 | Alignment | 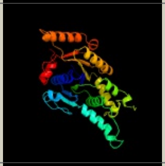 | 100.0 | 24 | **Fold:**P-loop containing nucleoside triphosphate hydrolases<br>**Superfamily:**P-loop containing nucleoside triphosphate hydrolases<br>**Family:**Nitrogenase iron protein-like |
| 14 | c3io3A_ | Alignment | 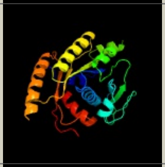 | 100.0 | 59 | **PDB header:**chaperone<br>**Chain:** A: **PDB Molecule:**deha2d07832p;<br>**PDBTitle:** get3 with adp from d. hansenii in closed form |
| 15 | d1ihua2 | Alignment | 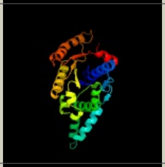 | 100.0 | 26 | **Fold:**P-loop containing nucleoside triphosphate hydrolases<br>**Superfamily:**P-loop containing nucleoside triphosphate hydrolases<br>**Family:**Nitrogenase iron protein-like |
| 16 | c3kjgB_ | Alignment | 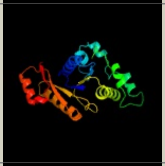 | 99.9 | 17 | **PDB header:**hydrolase, metal binding protein<br>**Chain:** B: **PDB Molecule:**co dehydrogenase/acetyl-coa synthase complex, accessory<br>**PDBTitle:** adp-bound state of cooc1 |
| 17 | d2afhe1 | Alignment | 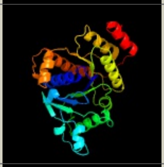 | 99.9 | 14 | **Fold:**P-loop containing nucleoside triphosphate hydrolases<br>**Superfamily:**P-loop containing nucleoside triphosphate hydrolases<br>**Family:**Nitrogenase iron protein-like |
| 18 | c6g2gA_ | Alignment | 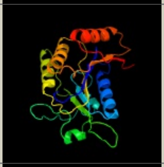 | 99.9 | 16 | **PDB header:**cytosolic protein<br>**Chain:** A: **PDB Molecule:**cytosolic fe-s cluster assembly factor cfd1;<br>**PDBTitle:** fe-s assembly cfd1 |
| 19 | c2ozeA_ | Alignment | 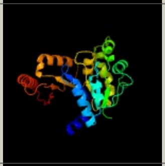 | 99.9 | 22 | **PDB header:**dna binding protein<br>**Chain:** A: **PDB Molecule:**orf delta';<br>**PDBTitle:** the crystal structure of delta protein of psm19035 from2 streptoccocus pyogenes |
| 20 | c2ph1A_ | Alignment | 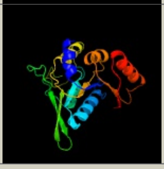 | 99.9 | 20 | **PDB header:**ligand binding protein<br>**Chain:** A: **PDB Molecule:**nucleotide-binding protein;<br>**PDBTitle:** crystal structure of nucleotide-binding protein af2382 from2 archaeoglobus fulgidus, northeast structural genomics target gr165 |
| 21 | c6iucC_ | Alignment | not modelled | 99.9 | 20 | **PDB header:**dna binding protein/dna<br>**Chain:** C: **PDB Molecule:**spooj regulator (soj);<br>**PDBTitle:** structure of helicobacter pylori soj-atp complex bound to dna |
| 22 | c4ru8C_ | Alignment | not modelled | 99.9 | 20 | **PDB header:**hydrolase<br>**Chain:** C: **PDB Molecule:**uncharacterized protein;<br>**PDBTitle:** structure of pnob8 para with amppnp |
| 23 | c3endA_ | Alignment | not modelled | 99.9 | 17 | **PDB header:**oxidoreductase<br>**Chain:** A: **PDB Molecule:**light-independent protochlorophyllide reductase<br>**PDBTitle:** crystal structure of the l protein of rhodobacter2 sphaeroides light-independent protochlorophyllide3 reductase (bchl) with mgadp bound: a homologue of the4 nitrogenase fe protein |
| 24 | c3ez6B_ | Alignment | not modelled | 99.9 | 17 | **PDB header:**dna binding protein<br>**Chain:** B: **PDB Molecule:**plasmid partition protein a;<br>**PDBTitle:** structure of para-adp complex:tetragonal form |
| 25 | c3cioA_ | Alignment | not modelled | 99.9 | 14 | **PDB header:**signaling protein, transferase<br>**Chain:** A: **PDB Molecule:**tyrosine-protein kinase etk;<br>**PDBTitle:** the kinase domain of escherichia coli tyrosine kinase etk |
| 26 | c3k9gA_ | Alignment | not modelled | 99.9 | 18 | **PDB header:**biosynthetic protein<br>**Chain:** A: **PDB Molecule:**pf-32 protein;<br>**PDBTitle:** crystal structure of a plasmid partition protein from borrelia2 burgdorferi at 2.25a resolution, iodide soak |
| 27 | c3fkqA_ | Alignment | not modelled | 99.9 | 20 | **PDB header:**structural genomics, unknown function<br>**Chain:** A: **PDB Molecule:**ntrc-like two-domain protein;<br>**PDBTitle:** crystal structure of ntrc-like two-domain protein (rer070207001320)2 from eubacterium rectale at 2.10 a resolution |
| 28 | c4rz3B_ | Alignment | not modelled | 99.9 | 23 | **PDB header:**structural protein<br>**Chain:** B: **PDB Molecule:**site-determining protein;<br>**PDBTitle:** crystal structure of the mind-like atpase flhg |

| | | | | | | |
|---|---|---|---|---|---|---|
| 29 | c5j1jA_ | Alignment | not modelled | 99.9 | 21 | **PDB header:**transcription<br>**Chain:** A: **PDB Molecule:**site-determining protein;<br>**PDBTitle:** structure of flen-amppnp complex |
| 30 | d1cp2a_ | Alignment | not modelled | 99.9 | 17 | **Fold:**P-loop containing nucleoside triphosphate hydrolases<br>**Superfamily:**P-loop containing nucleoside triphosphate hydrolases<br>**Family:**Nitrogenase iron protein-like |
| 31 | c3la6P_ | Alignment | not modelled | 99.9 | 18 | **PDB header:**transferase<br>**Chain:** P: **PDB Molecule:**tyrosine-protein kinase wzc;<br>**PDBTitle:** octameric kinase domain of the e. coli tyrosine kinase wzc with bound2 adp |
| 32 | c3pg5A_ | Alignment | not modelled | 99.9 | 24 | **PDB header:**structural genomics, unknown function<br>**Chain:** A: **PDB Molecule:**uncharacterized protein;<br>**PDBTitle:** crystal structure of protein dip2308 from corynebacterium diphtheriae,2 northeast structural genomics consortium target cdr78 |
| 33 | d1iona_ | Alignment | not modelled | 99.9 | 20 | **Fold:**P-loop containing nucleoside triphosphate hydrolases<br>**Superfamily:**P-loop containing nucleoside triphosphate hydrolases<br>**Family:**Nitrogenase iron protein-like |
| 34 | c6nonB_ | Alignment | not modelled | 99.9 | 16 | **PDB header:**dna binding protein<br>**Chain:** B: **PDB Molecule:**cobyrinic acid ac-diamide synthase;<br>**PDBTitle:** structure of cyanthece apo mcda |
| 35 | d1byia_ | Alignment | not modelled | 99.8 | 15 | **Fold:**P-loop containing nucleoside triphosphate hydrolases<br>**Superfamily:**P-loop containing nucleoside triphosphate hydrolases<br>**Family:**Nitrogenase iron protein-like |
| 36 | c3vx3A_ | Alignment | not modelled | 99.8 | 26 | **PDB header:**adp binding protein<br>**Chain:** A: **PDB Molecule:**atpase involved in chromosome partitioning, para/mind<br>**PDBTitle:** crystal structure of [nife] hydrogenase maturation protein hypb from2 thermococcus kodakarensis kod1 |
| 37 | c4v02B_ | Alignment | not modelled | 99.8 | 19 | **PDB header:**cell cycle<br>**Chain:** B: **PDB Molecule:**site-determining protein;<br>**PDBTitle:** minc:mind cell division protein complex, aquifex aeolicus |
| 38 | c2vedA_ | Alignment | not modelled | 99.8 | 17 | **PDB header:**transferase<br>**Chain:** A: **PDB Molecule:**membrane protein capa1, protein tyrosine kinase;<br>**PDBTitle:** crystal structure of the chimerical mutant capabk55m2 protein |
| 39 | c4pfsA_ | Alignment | not modelled | 99.8 | 17 | **PDB header:**ligase<br>**Chain:** A: **PDB Molecule:**cobyrinic acid a,c-diamide synthase;<br>**PDBTitle:** crystal structure of cobyrinic acid a,c-diamide synthase from2 mycobacterium smegmatis |
| 40 | d1g3qa_ | Alignment | not modelled | 99.8 | 21 | **Fold:**P-loop containing nucleoside triphosphate hydrolases<br>**Superfamily:**P-loop containing nucleoside triphosphate hydrolases<br>**Family:**Nitrogenase iron protein-like |
| 41 | c3q9lB_ | Alignment | not modelled | 99.8 | 24 | **PDB header:**cell cycle, hydrolase<br>**Chain:** B: **PDB Molecule:**septum site-determining protein mind;<br>**PDBTitle:** the structure of the dimeric e.coli mind-atp complex |
| 42 | c3of5A_ | Alignment | not modelled | 99.8 | 15 | **PDB header:**ligase<br>**Chain:** A: **PDB Molecule:**dethiobiotin synthetase;<br>**PDBTitle:** crystal structure of a dethiobiotin synthetase from francisella2 tularensis subsp. tularensis schu s4 |
| 43 | c2bekB_ | Alignment | not modelled | 99.8 | 21 | **PDB header:**chromosome segregation<br>**Chain:** B: **PDB Molecule:**segregation protein;<br>**PDBTitle:** structure of the bacterial chromosome segregation protein soj |
| 44 | c3ea0B_ | Alignment | not modelled | 99.8 | 16 | **PDB header:**hydrolase<br>**Chain:** B: **PDB Molecule:**atpase, para family;<br>**PDBTitle:** crystal structure of para family atpase from chlorobium tepidum tls |
| 45 | c3ezfA_ | Alignment | not modelled | 99.8 | 16 | **PDB header:**biosynthetic protein<br>**Chain:** A: **PDB Molecule:**para;<br>**PDBTitle:** partition protein |
| 46 | c6u1qA_ | Alignment | not modelled | 99.8 | 18 | **PDB header:**biosynthetic protein<br>**Chain:** A: **PDB Molecule:**vpso;<br>**PDBTitle:** crystal structure of vpso (vc0937) kinase domain |
| 47 | c2qmoA_ | Alignment | not modelled | 99.8 | 13 | **PDB header:**ligase<br>**Chain:** A: **PDB Molecule:**dethiobiotin synthetase;<br>**PDBTitle:** crystal structure of dethiobiotin synthetase (biod) from helicobacter2 pylori |
| 48 | d1hyqa_ | Alignment | not modelled | 99.8 | 20 | **Fold:**P-loop containing nucleoside triphosphate hydrolases<br>**Superfamily:**P-loop containing nucleoside triphosphate hydrolases<br>**Family:**Nitrogenase iron protein-like |
| 49 | c1hyqA_ | Alignment | not modelled | 99.8 | 20 | **PDB header:**cell cycle<br>**Chain:** A: **PDB Molecule:**cell division inhibitor (mind-1);<br>**PDBTitle:** mind bacterial cell division regulator from a. fulgidus |
| 50 | c3cwqB_ | Alignment | not modelled | 99.8 | 27 | **PDB header:**structural genomics, unknown function<br>**Chain:** B: **PDB Molecule:**para family chromosome partitioning protein;<br>**PDBTitle:** crystal structure of chromosome partitioning protein (para) in complex2 with adp from synechocystis sp. northeast structural genomics3 consortium target sgr89 |
| 51 | c4dzzB_ | Alignment | not modelled | 99.8 | 23 | **PDB header:**unknown function<br>**Chain:** B: **PDB Molecule:**plasmid partitioning protein parf;<br>**PDBTitle:** structure of parf-adp, crystal form 1 |
| 52 | c2xj9B_ | Alignment | not modelled | 99.8 | 15 | **PDB header:**replication<br>**Chain:** B: **PDB Molecule:**mipz;<br>**PDBTitle:** dimer structure of the bacterial cell division regulator mipz |
| 53 | c3fmfA_ | Alignment | not modelled | 99.7 | 13 | **PDB header:**ligase<br>**Chain:** A: **PDB Molecule:**dethiobiotin synthetase;<br>**PDBTitle:** crystal structure of mycobacterium tuberculosis dethiobiotin2 synthetase complexed with 7,8 diaminopelargonic acid carbamate |
| | | | | | | **PDB header:**protein transport |

| | | | | | | |
|---|---|---|---|---|---|---|
| 54 | c5l3qB_ | Alignment | not modelled | 99.6 | 19 | **Chain:** B: **PDB Molecule:**signal recognition particle receptor subunit alpha;<br>**PDBTitle:** structure of the gtpase heterodimer of human srp54 and sralpha |
| 55 | c2qy9A_ | Alignment | not modelled | 99.4 | 19 | **PDB header:**protein transport<br>**Chain:** A: **PDB Molecule:**cell division protein ftsy;<br>**PDBTitle:** structure of the ng+1 construct of the e. coli srp receptor2 ftsy |
| 56 | c1zu4A_ | Alignment | not modelled | 99.4 | 18 | **PDB header:**protein transport<br>**Chain:** A: **PDB Molecule:**ftsy;<br>**PDBTitle:** crystal structure of ftsy from mycoplasma mycoides-space2 group p21212 |
| 57 | c2og2A_ | Alignment | not modelled | 99.4 | 15 | **PDB header:**protein transport<br>**Chain:** A: **PDB Molecule:**putative signal recognition particle receptor;<br>**PDBTitle:** crystal structure of chloroplast ftsy from arabidopsis2 thaliana |
| 58 | c2cnwF_ | Alignment | not modelled | 99.4 | 17 | **PDB header:**signal recognition<br>**Chain:** F: **PDB Molecule:**cell division protein ftsy;<br>**PDBTitle:** gdpalf4 complex of the srp gtpases ffh and ftsy |
| 59 | c2j7pA_ | Alignment | not modelled | 99.4 | 17 | **PDB header:**signal recognition<br>**Chain:** A: **PDB Molecule:**signal recognition particle protein;<br>**PDBTitle:** gmppnp-stabilized ng domain complex of the srp gtpases ffh2 and ftsy |
| 60 | c5l3rC_ | Alignment | not modelled | 99.3 | 19 | **PDB header:**protein transport<br>**Chain:** C: **PDB Molecule:**signal recognition particle 54 kda protein, chloroplastic;<br>**PDBTitle:** structure of the gtpase heterodimer of chloroplast srp54 and ftsy from2 arabidopsis thaliana |
| 61 | c2q9cA_ | Alignment | not modelled | 99.3 | 16 | **PDB header:**signaling protein<br>**Chain:** A: **PDB Molecule:**cell division protein ftsy;<br>**PDBTitle:** structure of ftsy:gmppnp with mgcl complex |
| 62 | c2yhsA_ | Alignment | not modelled | 99.3 | 19 | **PDB header:**cell cycle<br>**Chain:** A: **PDB Molecule:**cell division protein ftsy;<br>**PDBTitle:** structure of the e. coli srp receptor ftsy |
| 63 | c1vmaA_ | Alignment | not modelled | 99.3 | 17 | **PDB header:**protein transport<br>**Chain:** A: **PDB Molecule:**cell division protein ftsy;<br>**PDBTitle:** crystal structure of cell division protein ftsy (tm0570) from2 thermotoga maritima at 1.60 a resolution |
| 64 | c3b9qA_ | Alignment | not modelled | 99.3 | 16 | **PDB header:**protein transport<br>**Chain:** A: **PDB Molecule:**chloroplast srp receptor homolog, alpha subunit<br>**PDBTitle:** the crystal structure of cpftsy from arabidopsis thaliana |
| 65 | c2iy3A_ | Alignment | not modelled | 99.3 | 21 | **PDB header:**rna-binding<br>**Chain:** A: **PDB Molecule:**signal recognition particle protein,signal recognition<br>**PDBTitle:** structure of the e. coli signal regognition particle |
| 66 | c3dm5A_ | Alignment | not modelled | 99.3 | 20 | **PDB header:**rna binding protein, transport protein<br>**Chain:** A: **PDB Molecule:**signal recognition 54 kda protein;<br>**PDBTitle:** structures of srp54 and srp19, the two proteins assembling the2 ribonucleic core of the signal recognition particle from the archaeon3 pyrococcus furiosus. |
| 67 | c5l3sF_ | Alignment | not modelled | 99.3 | 17 | **PDB header:**protein transport<br>**Chain:** F: **PDB Molecule:**signal recognition particle receptor ftsy;<br>**PDBTitle:** structure of the gtpase heterodimer of crenarchaeal srp54 and ftsy |
| 68 | c2j37W_ | Alignment | not modelled | 99.3 | 19 | **PDB header:**ribosome<br>**Chain:** W: **PDB Molecule:**signal recognition particle 54 kda protein (srp54);<br>**PDBTitle:** model of mammalian srp bound to 80s rncs |
| 69 | c6cy1B_ | Alignment | not modelled | 99.3 | 17 | **PDB header:**signaling protein<br>**Chain:** B: **PDB Molecule:**signal recognition particle receptor ftsy;<br>**PDBTitle:** crystal structure of signal recognition particle receptor ftsy from2 elizabethkingia anophelis |
| 70 | c5gafi_ | Alignment | not modelled | 99.2 | 21 | **PDB header:**ribosome<br>**Chain:** I: **PDB Molecule:**50s ribosomal protein l10;<br>**PDBTitle:** rnc in complex with srp |
| 71 | c4ak9A_ | Alignment | not modelled | 99.2 | 17 | **PDB header:**protein transport<br>**Chain:** A: **PDB Molecule:**cpftsy;<br>**PDBTitle:** structure of chloroplast ftsy from physcomitrella patens |
| 72 | c3dmdA_ | Alignment | not modelled | 99.2 | 19 | **PDB header:**transport protein<br>**Chain:** A: **PDB Molecule:**signal recognition particle receptor;<br>**PDBTitle:** structures and conformations in solution of the signal recognition2 particle receptor from the archaeon pyrococcus furiosus |
| 73 | c1qzwC_ | Alignment | not modelled | 99.2 | 21 | **PDB header:**signaling protein/rna<br>**Chain:** C: **PDB Molecule:**signal recognition 54 kda protein;<br>**PDBTitle:** crystal structure of the complete core of archaeal srp and2 implications for inter-domain communication |
| 74 | c2j289_ | Alignment | not modelled | 99.0 | 21 | **PDB header:**ribosome<br>**Chain:** 9: **PDB Molecule:**signal recognition particle 54;<br>**PDBTitle:** model of e. coli srp bound to 70s rncs |
| 75 | c2v3cC_ | Alignment | not modelled | 98.9 | 24 | **PDB header:**signaling protein<br>**Chain:** C: **PDB Molecule:**signal recognition 54 kda protein;<br>**PDBTitle:** crystal structure of the srp54-srp19-7s.s srp rna complex2 of m. jannaschii |
| 76 | d1vmaa2 | Alignment | not modelled | 98.5 | 18 | **Fold:**P-loop containing nucleoside triphosphate hydrolases<br>**Superfamily:**P-loop containing nucleoside triphosphate hydrolases<br>**Family:**Nitrogenase iron protein-like |
| 77 | d1qzxa3 | Alignment | not modelled | 98.5 | 15 | **Fold:**P-loop containing nucleoside triphosphate hydrolases<br>**Superfamily:**P-loop containing nucleoside triphosphate hydrolases<br>**Family:**Nitrogenase iron protein-like |
| 78 | c2px0D_ | Alignment | not modelled | 98.4 | 23 | **PDB header:**biosynthetic protein<br>**Chain:** D: **PDB Molecule:**flagellar biosynthesis protein flhf;<br>**PDBTitle:** crystal structure of flhf complexed with gmppnp/mg(2+) |
| | | | | | | **Fold:**P-loop containing nucleoside triphosphate hydrolases |

| | | | | | | |
|---|---|---|---|---|---|---|
| 79 | d1j8yf2 | Alignment | not modelled | 98.4 | 17 | **Superfamily:**P-loop containing nucleoside triphosphate hydrolases<br>**Family:**Nitrogenase iron protein-like |
| 80 | c3nvaB_ | Alignment | not modelled | 98.1 | 22 | **PDB header:**ligase<br>**Chain:** B: **PDB Molecule:**ctp synthase;<br>**PDBTitle:** dimeric form of ctp synthase from sulfolobus solfataricus |
| 81 | c4a0gC_ | Alignment | not modelled | 98.1 | 16 | **PDB header:**transferase<br>**Chain:** C: **PDB Molecule:**adenosylmethionine-8-amino-7-oxononanoate<br>**PDBTitle:** structure of bifunctional dapa aminotransferase-dtb synthetase from2 arabidopsis thaliana in its apo form. |
| 82 | c1j8yF_ | Alignment | not modelled | 98.1 | 17 | **PDB header:**signaling protein<br>**Chain:** F: **PDB Molecule:**signal recognition 54 kda protein;<br>**PDBTitle:** signal recognition particle conserved gtpase domain from a.2 ambivalens t112a mutant |
| 83 | c4a0rB_ | Alignment | not modelled | 98.0 | 16 | **PDB header:**transferase<br>**Chain:** B: **PDB Molecule:**adenosylmethionine-8-amino-7-oxononanoate<br>**PDBTitle:** structure of bifunctional dapa aminotransferase-dtb synthetase from2 arabidopsis thaliana bound to dethiobiotin (dtb). |
| 84 | d1vcoa2 | Alignment | not modelled | 98.0 | 19 | **Fold:**P-loop containing nucleoside triphosphate hydrolases<br>**Superfamily:**P-loop containing nucleoside triphosphate hydrolases<br>**Family:**Nitrogenase iron protein-like |
| 85 | d1mo6a1 | Alignment | not modelled | 97.9 | 13 | **Fold:**P-loop containing nucleoside triphosphate hydrolases<br>**Superfamily:**P-loop containing nucleoside triphosphate hydrolases<br>**Family:**RecA protein-like (ATPase-domain) |
| 86 | c4ydsA_ | Alignment | not modelled | 97.9 | 12 | **PDB header:**hydrolase<br>**Chain:** A: **PDB Molecule:**flagella-related protein h;<br>**PDBTitle:** flah from sulfolobus acidocaldarius with atp and mg-ion |
| 87 | c2recB_ | Alignment | not modelled | 97.9 | 13 | **PDB header:**helicase<br>**PDB COMPND:** |
| 88 | c1vcnA_ | Alignment | not modelled | 97.9 | 20 | **PDB header:**ligase<br>**Chain:** A: **PDB Molecule:**ctp synthetase;<br>**PDBTitle:** crystal structure of t.th. hb8 ctp synthetase complex with sulfate2 anion |
| 89 | d1xp8a1 | Alignment | not modelled | 97.9 | 12 | **Fold:**P-loop containing nucleoside triphosphate hydrolases<br>**Superfamily:**P-loop containing nucleoside triphosphate hydrolases<br>**Family:**RecA protein-like (ATPase-domain) |
| 90 | d2qy9a2 | Alignment | not modelled | 97.9 | 35 | **Fold:**P-loop containing nucleoside triphosphate hydrolases<br>**Superfamily:**P-loop containing nucleoside triphosphate hydrolases<br>**Family:**Nitrogenase iron protein-like |
| 91 | c5u03C_ | Alignment | not modelled | 97.9 | 21 | **PDB header:**ligase, protein fibril<br>**Chain:** C: **PDB Molecule:**ctp synthase 1;<br>**PDBTitle:** cryo-em structure of the human ctp synthase filament |
| 92 | d1okkd2 | Alignment | not modelled | 97.8 | 25 | **Fold:**P-loop containing nucleoside triphosphate hydrolases<br>**Superfamily:**P-loop containing nucleoside triphosphate hydrolases<br>**Family:**Nitrogenase iron protein-like |
| 93 | c6l6zH_ | Alignment | not modelled | 97.8 | 22 | **PDB header:**ligase<br>**Chain:** H: **PDB Molecule:**ctp synthase;<br>**PDBTitle:** cryo-em structure of the drosophila ctp synthase substrate-bound2 filament |
| 94 | d1s1ma2 | Alignment | not modelled | 97.8 | 22 | **Fold:**P-loop containing nucleoside triphosphate hydrolases<br>**Superfamily:**P-loop containing nucleoside triphosphate hydrolases<br>**Family:**Nitrogenase iron protein-like |
| 95 | c1xp8A_ | Alignment | not modelled | 97.8 | 19 | **PDB header:**dna binding protein<br>**Chain:** A: **PDB Molecule:**reca protein;<br>**PDBTitle:** deinococcus radiodurans reca in complex with atp-gamma-s |
| 96 | c3hr8A_ | Alignment | not modelled | 97.8 | 14 | **PDB header:**recombination<br>**Chain:** A: **PDB Molecule:**protein reca;<br>**PDBTitle:** crystal structure of thermotoga maritima reca |
| 97 | c4zdiE_ | Alignment | not modelled | 97.8 | 15 | **PDB header:**ligase<br>**Chain:** E: **PDB Molecule:**ctp synthase;<br>**PDBTitle:** crystal structure of the m. tuberculosis ctp synthase pyrg (apo form) |
| 98 | c2ad5B_ | Alignment | not modelled | 97.7 | 18 | **PDB header:**ligase<br>**Chain:** B: **PDB Molecule:**ctp synthase;<br>**PDBTitle:** mechanisms of feedback regulation and drug resistance of ctp2 synthetases: structure of the e. coli ctps/ctp complex at 2.8-3 angstrom resolution. |
| 99 | c4ohvA_ | Alignment | not modelled | 97.7 | 21 | **PDB header:**rna binding protein<br>**Chain:** A: **PDB Molecule:**protein clpf-1;<br>**PDBTitle:** c. elegans clp1 bound to amp-pnp, and mg2+ |
| 100 | d1eg7a_ | Alignment | not modelled | 97.7 | 23 | **Fold:**P-loop containing nucleoside triphosphate hydrolases<br>**Superfamily:**P-loop containing nucleoside triphosphate hydrolases<br>**Family:**Nitrogenase iron protein-like |
| 101 | d1tf7a2 | Alignment | not modelled | 97.7 | 17 | **Fold:**P-loop containing nucleoside triphosphate hydrolases<br>**Superfamily:**P-loop containing nucleoside triphosphate hydrolases<br>**Family:**RecA protein-like (ATPase-domain) |
| 102 | c4nmnA_ | Alignment | not modelled | 97.7 | 12 | **PDB header:**replication<br>**Chain:** A: **PDB Molecule:**replicative dna helicase;<br>**PDBTitle:** aquifex aeolicus replicative helicase (dnab) complexed with adp, at2 3.3 resolution |
| 103 | c2zroA_ | Alignment | not modelled | 97.7 | 23 | **PDB header:**hydrolase<br>**Chain:** A: **PDB Molecule:**protein reca;<br>**PDBTitle:** msreca adp form iv |
| 104 | d1u94a1 | Alignment | not modelled | 97.7 | 14 | **Fold:**P-loop containing nucleoside triphosphate hydrolases<br>**Superfamily:**P-loop containing nucleoside triphosphate hydrolases<br>**Family:**RecA protein-like (ATPase-domain) |
| 105 | d1ls1a2 | Alignment | not modelled | 97.7 | 29 | **Fold:**P-loop containing nucleoside triphosphate hydrolases<br>**Superfamily:**P-loop containing nucleoside triphosphate hydrolases |

| | | | | | | |
|---|---|---|---|---|---|---|
| | | Alignment | | | | **Family:**Nitrogenase iron protein-like |
| 106 | c4wiaA_ | Alignment | not modelled | 97.7 | 19 | **PDB header:**atp-binding protein<br>**Chain:** A: **PDB Molecule:**putative flagella-related protein h;<br>**PDBTitle:** crystal structure of flagellar accessory protein flah from2 methanocaldococcus jannaschii |
| 107 | c3cmvG_ | Alignment | not modelled | 97.7 | 15 | **PDB header:**recombination<br>**Chain:** G: **PDB Molecule:**protein reca;<br>**PDBTitle:** mechanism of homologous recombination from the reca-ssdna/dsdna2 structures |
| 108 | d1ubea1 | Alignment | not modelled | 97.7 | 16 | **Fold:**P-loop containing nucleoside triphosphate hydrolases<br>**Superfamily:**P-loop containing nucleoside triphosphate hydrolases<br>**Family:**RecA protein-like (ATPase-domain) |
| 109 | c3do6B_ | Alignment | not modelled | 97.6 | 25 | **PDB header:**ligase<br>**Chain:** B: **PDB Molecule:**formate--tetrahydrofolate ligase;<br>**PDBTitle:** crystal structure of putative formyltetrahydrofolate synthetase2 (tm1766) from thermotoga maritima at 1.85 a resolution |
| 110 | c4zc0A_ | Alignment | not modelled | 97.6 | 12 | **PDB header:**hydrolase<br>**Chain:** A: **PDB Molecule:**replicative dna helicase;<br>**PDBTitle:** structure of a dodecameric bacterial helicase |
| 111 | c2q6tB_ | Alignment | not modelled | 97.6 | 14 | **PDB header:**hydrolase<br>**Chain:** B: **PDB Molecule:**dnab replication fork helicase;<br>**PDBTitle:** crystal structure of the thermus aquaticus dnab monomer |
| 112 | c2vyeA_ | Alignment | not modelled | 97.6 | 17 | **PDB header:**hydrolase/dna<br>**Chain:** A: **PDB Molecule:**replicative dna helicase;<br>**PDBTitle:** crystal structure of the dnac-ssdna complex |
| 113 | d2vo1a1 | Alignment | not modelled | 97.6 | 21 | **Fold:**P-loop containing nucleoside triphosphate hydrolases<br>**Superfamily:**P-loop containing nucleoside triphosphate hydrolases<br>**Family:**Nitrogenase iron protein-like |
| 114 | c5x06G_ | Alignment | not modelled | 97.6 | 18 | **PDB header:**replication<br>**Chain:** G: **PDB Molecule:**dnaa regulatory inactivator hda;<br>**PDBTitle:** dna replication regulation protein |
| 115 | c4a1fB_ | Alignment | not modelled | 97.5 | 12 | **PDB header:**hydrolase<br>**Chain:** B: **PDB Molecule:**replicative dna helicase;<br>**PDBTitle:** crystal structure of c-terminal domain of helicobacter2 pylori dnab helicase |
| 116 | c2f1rA_ | Alignment | not modelled | 97.5 | 13 | **PDB header:**biosynthetic protein<br>**Chain:** A: **PDB Molecule:**molybdopterin-guanine dinucleotide biosynthesis<br>**PDBTitle:** crystal structure of molybdopterin-guanine biosynthesis2 protein b (mobb) |
| 117 | c2ztsB_ | Alignment | not modelled | 97.5 | 15 | **PDB header:**atp-binding protein<br>**Chain:** B: **PDB Molecule:**putative uncharacterized protein ph0186;<br>**PDBTitle:** crystal structure of kaic-like protein ph0186 from hyperthermophilic2 archaea pyrococcus horikoshii ot3 |
| 118 | d1nija1 | Alignment | not modelled | 97.5 | 12 | **Fold:**P-loop containing nucleoside triphosphate hydrolases<br>**Superfamily:**P-loop containing nucleoside triphosphate hydrolases<br>**Family:**Nitrogenase iron protein-like |
| 119 | c5a4jC_ | Alignment | not modelled | 97.5 | 19 | **PDB header:**ligase<br>**Chain:** C: **PDB Molecule:**formate--tetrahydrofolate ligase;<br>**PDBTitle:** crystal structure of fthfs1 from t.acetoxydans re1 |
| 120 | c5hcnA_ | Alignment | not modelled | 97.5 | 32 | **PDB header:**hydrolase<br>**Chain:** A: **PDB Molecule:**gpn-loop gtpase 1;<br>**PDBTitle:** gpn-loop gtpase npa3 in complex with gmppcp |
