## Supplementary Figure 7 for "A conserved Guided Entry of Tail-anchored pathway is involved in the trafficking of tail-anchored membrane proteins in *Plasmodium falciparum*"

**A**

| Model No. | Oligomer template | No. of subunits | Interface area | Sequence identity | Structure similarity (TM-score) |
|---|---|---|---|---|---|
| 1 | 2WOO | 2-mer | 4199.1 | 40.6 | 0.8660 |
| 2 | 3IQW | 2-mer | 4019.8 | 39.8 | 0.7945 |
| 3 | 3IBG | 2-mer | 2514.3 | 40.6 | 0.7831 |
| 4 | 3SJA | 2-mer | 2408.1 | 39.1 | 0.7857 |
| 5 | 3UG7 | 4-mer | 10787.4 | 26.9 | 0.7099 |

**B**

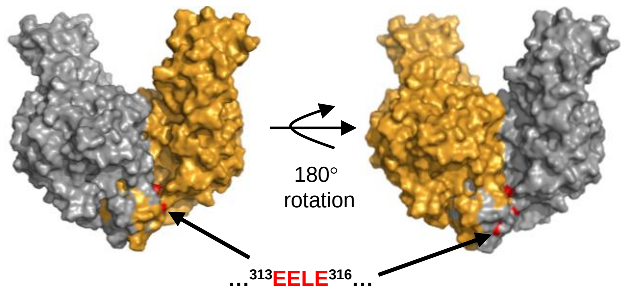


**C**

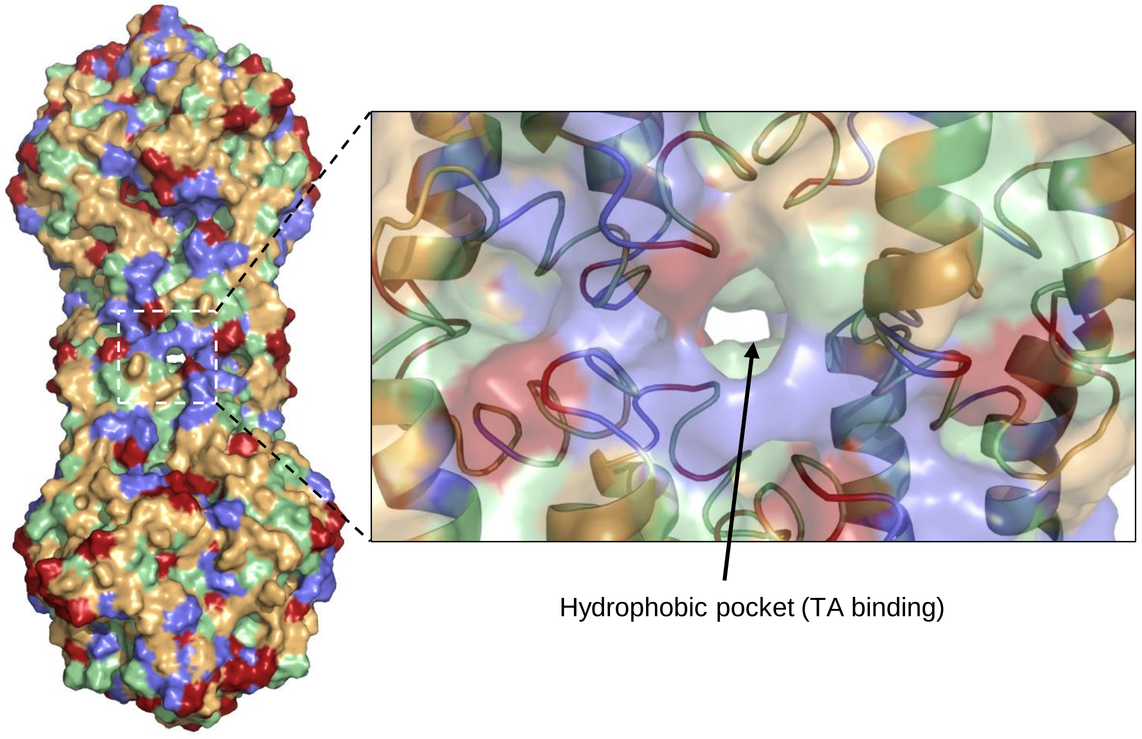
