## Supplementary Figure 8 for "A conserved Guided Entry of Tail-anchored pathway is involved in the trafficking of tail-anchored membrane proteins in *Plasmodium falciparum*"

**A**

| ID | *P. falciparum* 3D7 Get4 Homolog (Pf3D7_1438600) | Identity | Similarity |
|---|---|---|---|
| **Yor164C** | *S. cerevisiae* S288C | 17.6% | 36.3% |
| **TRC35** | *H. sapiens* | 15.6% | 37.8% |
| **At5g63220** | *A. thaliana* | 17.8% | 36.7% |
| **BN1205_070760** | *T. gondii* | 28.1% | 47.5% |
| **CTHT_0028730** | *C. thermophilum* | 17.2% | 30.5% |

```
PF3D7_1438600      1 ----------------MKKFKFSKEKLAKEKIENGLHYEAHQLIKSIINR     34
                                     :::|        :.||:.|.:|||||.:::|.||
Sc_Yor164c         1 MVPAESNAVQAKLAKTLQRF--------ENKIKAGDYYEAHQTLRTIANR     42

PF3D7_1438600     35 KLQKNQISNCISVIFYFSLKFATAEQYVLLCDLMYQCCLILTQHDIEFNL     84
                     .::.....:.|.:|...:|.|..|:|.....||::....:....:::.:.
Sc_Yor164c        43 YVRSKSYEHAIELISQGALSFLKAKQGGSGTDLIFYLLEVYDLAEVKVDD     92

PF3D7_1438600     85 EYANKIIEIFQKCPPN--STEEKYKFMNKSILWS-KSEENVYGYLEFHKV    131
                     ....:::.:..:..|:  :.::....||.   || |..|..:|....|..
Sc_Yor164c        93 ISVARLVRLIAELDPSEPNLKDVITGMNN---WSIKFSEYKFGDPYLHNT    139

PF3D7_1438600    132 IG-------YAYFEDKQYSLAHNHFIYLDDNDILYEIICLWRKEAYPSES    174
                     ||       :.| |.::|.:...|...:...|:|::.:|........:.:
Sc_Yor164c       140 IGSKLLEGDFVY-EAERYFMLGTHDSMIKYVDLLWDWLCQVDDIEDSTVA    188

PF3D7_1438600    175 HFFVLRTILCLLV-----------------LDKFDQALNLIN-------M    200
                     .||.......|.:                 ::||......|:       .
Sc_Yor164c       189 EFFSRLVFNYLFISNISFAHESKDIFLERFIEKFHPKYEKIDKNGYEIVF    238

PF3D7_1438600    201 FE--TNLNRQDVPLPIQLAYLITSSCIYKSLALYEDVKYKYRLILNYDPD    248
                     ||  ::||         ...|:..:|..|..:.:.::|..|   |::...
Sc_Yor164c       239 FEDYSDLN---------FLQLLLITCQTKDKSYFLNLKNHY---LDFSQA    276

PF3D7_1438600    249 FQKYINIIDAAVFN----KKKHNLFSMFQNIFA---    277
                     ::..:..:....||    |:.:.|..|......
Sc_Yor164c       277 YKSELEFLGQEYFNIVAPKQTNFLQDMMSGFLGGSK    312
```

### Supplementary Figure 8 (contd.)

```
PF3D7_1438600     1 MKKFKFSKEKLAKE-------------KIENGLHYEAHQLIKSIINRKLQ     37
                         .|:|::.:|             .||.|.:|.|.|:.|||
At5g63220         1 -----MSRERIKRELPPVQEHIDKLRKVIEEGNYYGALQMYKSI------     39

PF3D7_1438600    38 KNQISNCISVIFYFSLKFATAEQYVLLCDLMYQCCLILTQHDI-------     80
                                  |.::.||:::....|:::....|..:|.:
At5g63220        40 --------------SARYVTAQRFSEALDILFSGACIELEHGLVNCGADL     75

PF3D7_1438600    81 ----------------EFNLEYANKIIEIFQK--CPPN----STEEKYKF    108
                                    :..|:....|.::|.:  .||:    |.:|..:.
At5g63220        76 AILFVDTLVKAKSPCNDETLDRIRCIFKLFPRVPVPPHLVDVSDDEDVQN    125

PF3D7_1438600   109 MNKSILWSKSE-ENV------------------YGYLEFHKVIG-YAYFE    138
                    :.:|:..::|. ||:                  .||.|.|.::| |.|.|
At5g63220       126 LQESLGEARSRVENLTSFLRAAIKWSAEFGGPRTGYPELHAMLGDYLYTE    175

PF3D7_1438600   139 DKQYSLAH--NHFIYLDDNDILYEIICLWRKEAYPSESHFFVLRTILCLL    186
                    ..:..:..  .||:..:|.:....::..:....||.|....:.|.:|..|
At5g63220       176 CPELDMVRISRHFVRAEDPEKFASMLVNFMGRCYPGEDDLAIARAVLMYL    225

PF3D7_1438600   187 VLDKFDQALNLINMFETNLNRQDVPLP----IQLAYLITSSCIYKSLALY    232
                    .:.....|..:::..:.....::..|.    ||....:..:....:|.|:
At5g63220       226 SMGNMKDANFMMDEIKKQAETKNPELSESDLIQFISYLLETLQRDALPLF    275

PF3D7_1438600   233 EDVKYKYRLILNYDPDFQKYINIIDAAVFN-KKKHNLFSMFQNIFA---    277
                    ..::.||:..::.|....:.::.|....:. ::|:.|..||.:||.
At5g63220       276 NMLRVKYKSSIDRDQLLNELLDEIAERFYGVQRKNPLQGMFGDIFKMMG    324


PF3D7_1438600     1 ----------------------MKKFKFSKEKLAKEKIENGLHYEAHQLI     28
                                          :::.    |...:..:|.|.:|||||:.
Hs_TRC35          1 MAAAAAMAEQESARNGGRNRGGVQRV----EGKLRASVEKGDYYEAHQMY     46

PF3D7_1438600    29 KSIINRKLQKNQISNCISVIFYFSLKFATAEQYVLLCDLMYQCCLILTQH     78
                    :::..|.:.:::.:....:::..:|.|.:..|.....||.......|.:.
Hs_TRC35         47 RTLFFRYMSQSKHTEARELMYSGALLFFSHGQQNSAADLSMLVLESLEKA     96

PF3D7_1438600    79 DIEFNLEYANKIIEIFQKCPPNSTEEKYKFMNKSILWSKSEENVYGYLEF    128
                    ::|...|....:.::|....||| .|:..|:::::.||.......|:...
Hs_TRC35         97 EVEVADELLENLAKVFSLMDPNS-PERVTFVSRALKWSSGGSGKLGHPRL    145

PF3D7_1438600   129 HKVIGYAYFEDKQYSLAHNHFIYLDDNDILYEIICLW-RKEAYPSESHFF    177
                    |:::....::::.|..:..||::..|.:....::..: ....:.||...|
Hs_TRC35        146 HQLLALTLWKEQNYCESRYHFLHSADGEGCANMLVEYSTSRGFRSEVDMF    195

PF3D7_1438600   178 VLRTILCLLVL-DKFDQALNLINMFETNLNRQDVPLP-----IQLAYLIT    221
                    |.:.:|..|.| :|...::......:.:.:.:|.| |     :...:.:.
Hs_TRC35        196 VAQAVLQFLCLKNKSSASVVFTTYTQKHPSIEDGP-PFVEPLLNFIWFLL    244

PF3D7_1438600   222 SSCIYKSLALYEDVKYKYRLILNYDPDFQKYINIIDAAVFN---KKKHNL    268
                    .:.....|.::..:..:|:..|..||.:.:|::.|....|.   |:..:.
Hs_TRC35        245 LAVDGGKLTVFTVLCEQYQPSLRRDPMYNEYLDRIGQLFFGVPPKQTSSY    294

PF3D7_1438600   269 FSMFQNIFA------------------------    277
                    ..:..|:..
Hs_TRC35        295 GGLLGNLLTSLMGSSEQEDGEESPSDGSPIELD    327
```

#### Supplementary Figure 8 (contd.)

```
PF3D7_1438600      1 MKKFKFSKEKLAKE-----------------------KIENGLHYEAHQL     27
                           .||::..:                       ||:||..|:||||
Tg_BN1205_070      1 ------MKERVQPQAPAGTDISGGSVLPLRVQASVDRKIDNGDLYDAHQL     44

PF3D7_1438600     28 IKSIINRKLQKNQISNCISVIFYFSLKFATAEQYVLLCDLMYQCCLILTQ     77
                     ::::..|.:.|.:....:.:...:.|:||..:|..|..||.......|..
Tg_BN1205_070     45 VRTLFFRFMAKRETLQAVELCRLYGLRFAGLDQEALAVDLGMNMLTALEA     94

PF3D7_1438600     78 HDIEFNLEYA------NKIIEIFQKCPPNSTE---EKYKFMNKSILWSKS    118
                     ...| ..|.|      ::|||:|..|.....:   :||||:|:::.||::
Tg_BN1205_070     95 SGTE-EEETAPSEPQLDQIIELFNACAAAGDKKGVDKYKFINRALKWSRT    143

PF3D7_1438600    119 EENVYGYLEFHKVIGYAYFEDKQYSLAHNHFIYLDDNDILYEIICLWRKE    168
                     .:..:|::..|:....||:::::|.|...|.||..|.:.|.:::..|:..
Tg_BN1205_070    144 PQAPFGHVRLHRAAADAYWKERRYGLCQGHLIYCRDPEALSQMLKEWQAT    193

PF3D7_1438600    169 AYPSESHFFVLRTILCLLVLDKFDQALNLINMFETNLNRQDVPLPIQLAY    218
                     .|.||..||.||.:|.||.|...:.|..|::....|....:||.|:||||
Tg_BN1205_070    194 GYLSERPFFWLRLVLILLCLRDTETAEKLLDSSGENWTSCEVPAPLQLAY    243

PF3D7_1438600    219 LITSSCIYKSLALYEDVKYKYRLILNYDPDFQKYINIIDAAVFNKKKH--    266
                     |:..:|.|||..|::.:|.||.|:|..||.|.||::.|:.....:.:.
Tg_BN1205_070    244 LLVCACKYKSGKLFDLLKQKYHLVLRRDPTFAKYMDEIEKRAIGRIQRPS    293

PF3D7_1438600    267 -NLFSMFQNIFA--------    277
                      .|.|:|.::.|
Tg_BN1205_070    294 TGLASLFSSLMAGLTADDEA    313


PF3D7_1438600      1 MKKFKFSKEKLAKEKIENGL--------------HYEAHQLIKSIINRKL     36
                                ...|||..:              .|||.|..:.:..|..
CTHT_0028730       1 -----------MSNKIERIIARLQRRIAEGQPEEQYEAAQETRLVAARYS     39

PF3D7_1438600     37 QKNQISNCISVIFYFSLKFATAEQ-------YVLLCDLMYQC--------     71
                     ::...:..:.::...|.....:.|       .|||.|...|.
CTHT_0028730      40 KQGNWAAAVDILASVSQTLLRSGQGGSGGDLAVLLVDTFRQAGQRVDGAS     89

PF3D7_1438600     72 ------CLILTQHDIEFNLEYANKIIEIFQKCPPNSTEEKYKFMNKSILW    115
                           ||                  .:||...|    .:.:|:.:.|.|
CTHT_0028730      90 RGKLLGCL------------------RLFQPGEP----VRKRFVKEMIDW    117

PF3D7_1438600    116 SKS-EENVYGYLEFHKVIGYAYFEDKQYSLAHNHFIY--LDDNDILYEII    162
                     ||. .:...|..|.|.|:|..|.|:.::..|..|.:.  .:..::|..:.
CTHT_0028730     118 SKKFGDYPAGDPELHHVVGTLYVEEGEFEAAEKHLVLGTKESPEVLARME    167

PF3D7_1438600    163 CLWRKEAYPSESH---FFVLRTILCLLVLDKFDQALNLINMFETNLNRQD    209
                     ..|.|:   .|||   .:..|.:|..|::.....|.....:|.:.|...:
CTHT_0028730     168 YEWYKQ---DESHTAPLYCARAVLPYLLVANVRAANTAYRIFTSALVEDN    214

PF3D7_1438600    210 VPLPIQ--------------------LAYLITSSCIYK-SLALYEDVKYK    238
                     ..|.:|                    ::.|:.|  :.| |..|:..:|.|
CTHT_0028730     215 KGLTVQNIGSQSAELRIFPSLPLLNFISMLLLS--VQKGSPDLFRQLKSK    262

PF3D7_1438600    239 YRLILN-YDPDFQKYINIIDAAVFN----KKKHNLFSMFQNIFA------    277
                     |...|| .:..:...:.:|....|.    ::.:.|..|..::|.
CTHT_0028730     263 YEANLNELNGIWDTALELIAEMYFGIQRPRQSNPLLDMMGSLFGGGGGAP    312

PF3D7_1438600    278 -----------------    277

CTHT_0028730     313 SKAALRRIDTPAAEGLD    329
```

#### Supplementary Figure 8 (contd.)

B

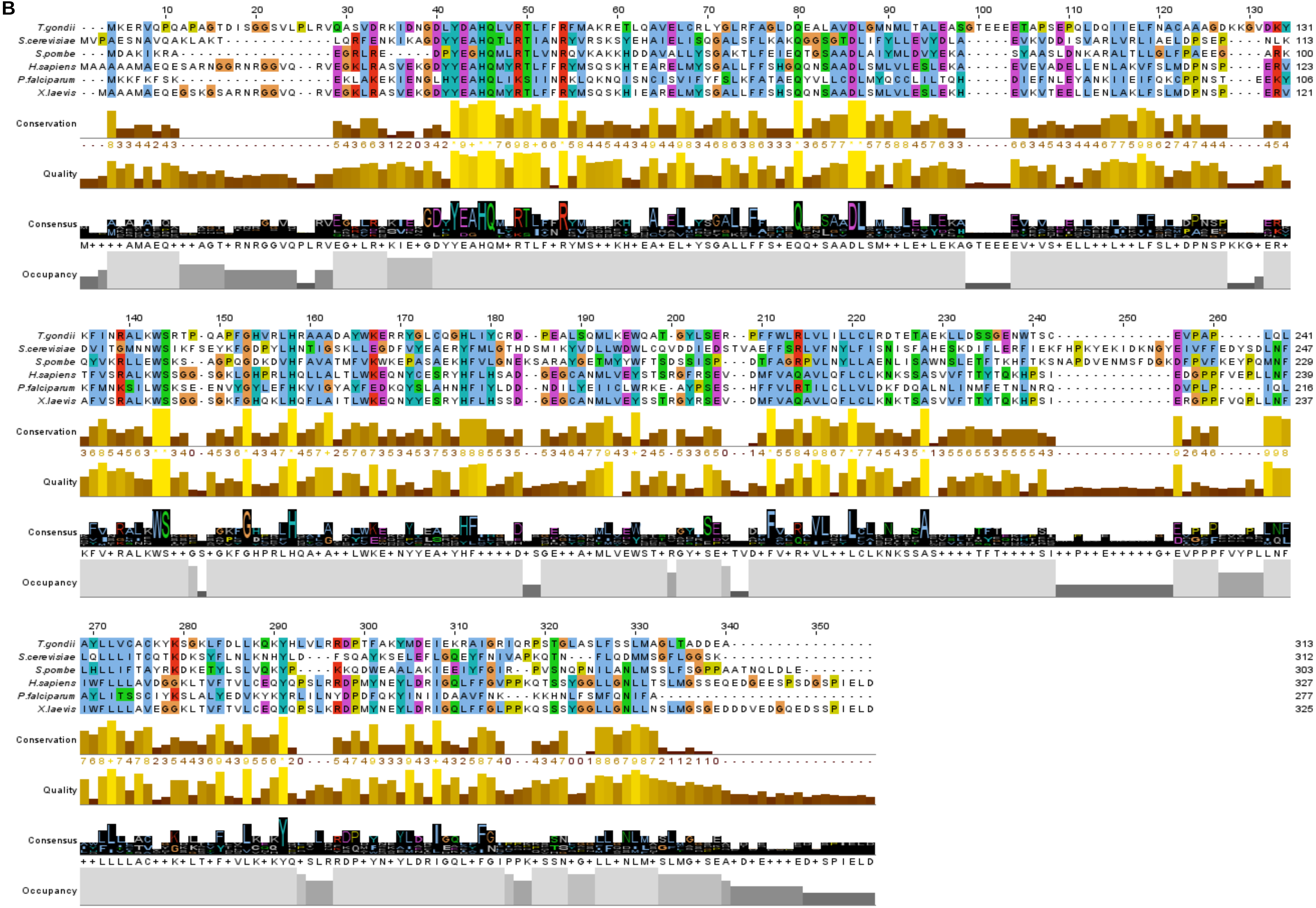
