## Supplementary Figure 11 for "A conserved Guided Entry of Tail-anchored pathway is involved in the trafficking of tail-anchored membrane proteins in *Plasmodium falciparum*"

Supplementary Figure 11. Hit report of PfGet3 predicted by Phyre2 server (www.sbg.bio.ic.ac.uk/phyre2). Schematic tabular representation showing the % confidence and identity of PfGet3 with the retrieved hits from Phyre2 server.

| # | Template | Alignment Coverage | 3D Model | Confidence | % i.d. | Template Information |
|---|---|---|---|---|---|---|
| 1 | c6au8A_ | Alignment | 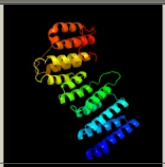 | 100.0 | 18 | **PDB header:**chaperone<br>**Chain:** A: **PDB Molecule:**golgi to er traffic protein 4 homolog;<br>**PDBTitle:** 1.8 angstrom crystal structure of the human bag6-nls & trc35 complex |
| 2 | c3lpzA_ | Alignment | 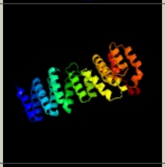 | 100.0 | 19 | **PDB header:**protein transport<br>**Chain:** A: **PDB Molecule:**uncharacterized protein;<br>**PDBTitle:** crystal structure of c. therm. get4 |
| 3 | c2wpvG_ | Alignment | 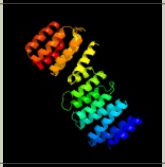 | 100.0 | 19 | **PDB header:**protein binding<br>**Chain:** G: **PDB Molecule:**upf0363 protein yor164c;<br>**PDBTitle:** crystal structure of s. cerevisiae get4-get5 complex |
| 4 | c3j98J_ | Alignment | 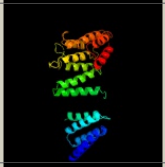 | 98.7 | 11 | **PDB header:**hydrolase<br>**Chain:** J: **PDB Molecule:**alpha-soluble nsf attachment protein;<br>**PDBTitle:** structure of 20s supercomplex determined by single particle2 cryoelectron microscopy (state iiia) |
| 5 | c4b4tQ_ | Alignment | 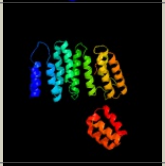 | 98.5 | 16 | **PDB header:**hydrolase<br>**Chain:** Q: **PDB Molecule:**26s proteasome regulatory subunit rpn6;<br>**PDBTitle:** near-atomic resolution structural model of the yeast 26s proteasome |
| 6 | c4cr3Q_ | Alignment | 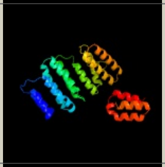 | 97.6 | 16 | **PDB header:**hydrolase<br>**Chain:** Q: **PDB Molecule:**26s proteasome regulatory subunit rpn6;<br>**PDBTitle:** deep classification of a large cryo-em dataset defines the2 conformational landscape of the 26s proteasome |
| 7 | d1qqea_ | Alignment | 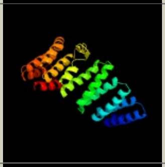 | 97.5 | 10 | **Fold:**alpha-alpha superhelix<br>**Superfamily:**TPR-like<br>**Family:**Tetratricopeptide repeat (TPR) |
| 8 | c3txmA_ | Alignment | 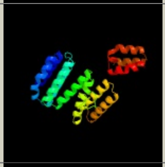 | 97.4 | 15 | **PDB header:**hydrolase, protein binding<br>**Chain:** A: **PDB Molecule:**26s proteasome regulatory complex subunit p42b;<br>**PDBTitle:** crystal structure of rpn6 from drosophila melanogaster, gd(3+) complex |
| 9 | c2ifuA_ | Alignment | 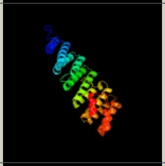 | 96.7 | 7 | **PDB header:**endocytosis/exocytosis<br>**Chain:** A: **PDB Molecule:**gamma-snap;<br>**PDBTitle:** crystal structure of a gamma-snap from danio rerio |
| 10 | c6mfvC_ | Alignment | 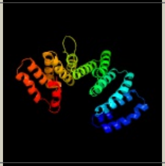 | 96.3 | 10 | **PDB header:**signaling protein<br>**Chain:** C: **PDB Molecule:**tetratricopeptide repeat sensor ph0952;<br>**PDBTitle:** crystal structure of the signal transduction atpase with numerous2 domains (stand) protein with a tetratricopeptide repeat sensor ph09523 from pyrococcus horikoshii |
| 11 | c4yvqC_ | Alignment | 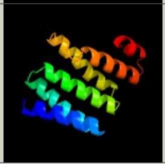 | 96.0 | 12 | **PDB header:**oxidoreductase/fluorescent protein<br>**Chain:** C: **PDB Molecule:**protein fluorescent in blue light, chloroplastic;<br>**PDBTitle:** crystal structure of flu-tpr in complex with the c-terminal region of2 glutr |

| | | | | | | |
|---|---|---|---|---|---|---|
| 12 | c5khrO_ | Alignment | 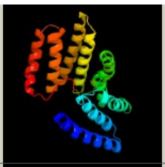 | 95.5 | 8 | **PDB header:**cell cycle<br>**Chain:** O: **PDB Molecule:**anaphase-promoting complex subunit 5;<br>**PDBTitle:** model of human anaphase-promoting complex/cyclosome complex (apc152 deletion mutant) in complex with the e2 ube2c/ubch10 poised for3 ubiquitin ligation to substrate (apc/c-cdc20-substrate-ube2c) |
| 13 | c5t0gV_ | Alignment | 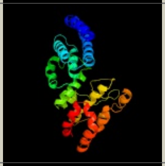 | 95.4 | 12 | **PDB header:**hydrolase<br>**Chain:** V: **PDB Molecule:**26s proteasome non-atpase regulatory subunit 3;<br>**PDBTitle:** structural basis for dynamic regulation of the human 26s proteasome |
| 14 | c5l4kS_ | Alignment | 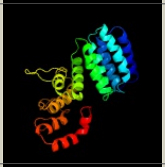 | 94.9 | 12 | **PDB header:**structural protein<br>**Chain:** S: **PDB Molecule:**26s proteasome non-atpase regulatory subunit 3;<br>**PDBTitle:** the human 26s proteasome lid |
| 15 | c4ui9O_ | Alignment | 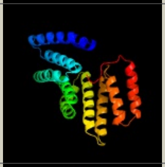 | 94.4 | 9 | **PDB header:**cell cycle<br>**Chain:** O: **PDB Molecule:**anaphase-promoting complex subunit 5;<br>**PDBTitle:** atomic structure of the human anaphase-promoting complex |
| 16 | c5gjqS_ | Alignment | 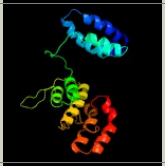 | 94.2 | 11 | **PDB header:**hydrolase<br>**Chain:** S: **PDB Molecule:**26s proteasome non-atpase regulatory subunit 3;<br>**PDBTitle:** structure of the human 26s proteasome bound to usp14-ubal |
| 17 | c4cr2P_ | Alignment | 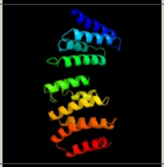 | 93.8 | 15 | **PDB header:**hydrolase<br>**Chain:** P: **PDB Molecule:**26s proteasome regulatory subunit rpn5;<br>**PDBTitle:** deep classification of a large cryo-em dataset defines the2 conformational landscape of the 26s proteasome |
| 18 | c3gw4B_ | Alignment | 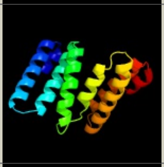 | 93.6 | 9 | **PDB header:**structural genomics, unknown function<br>**Chain:** B: **PDB Molecule:**uncharacterized protein;<br>**PDBTitle:** crystal structure of uncharacterized protein from deinococcus2 radiodurans. northeast structural genomics consortium target drr162b. |
| 19 | c3ro3A_ | Alignment | 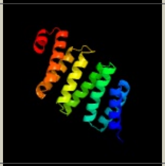 | 93.6 | 8 | **PDB header:**protein binding<br>**Chain:** A: **PDB Molecule:**g-protein-signaling modulator 2;<br>**PDBTitle:** crystal structure of lgn/minscuteable complex |
| 20 | c6r7nB_ | Alignment | 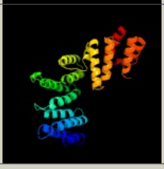 | 92.8 | 12 | **PDB header:**ligase<br>**Chain:** B: **PDB Molecule:**cop9 signalosome complex subunit 2;<br>**PDBTitle:** structural basis of cullin-2 ring e3 ligase regulation by the cop92 signalosome |
| 21 | c6c95A_ | Alignment | not modelled | 91.8 | 14 | **PDB header:**transferase<br>**Chain:** A: **PDB Molecule:**n-alpha-acetyltransferase 15, nata auxiliary subunit;<br>**PDBTitle:** the human nata (naa10/naa15) amino-terminal acetyltransferase complex2 bound to hypk |
| 22 | c5mpeP_ | Alignment | not modelled | 91.5 | 13 | **PDB header:**hydrolase<br>**Chain:** P: **PDB Molecule:**26s proteasome regulatory subunit rpn5;<br>**PDBTitle:** 26s proteasome in presence of atp (s2) |
| 23 | c1xi4D_ | Alignment | not modelled | 90.6 | 22 | **PDB header:**endocytosis/exocytosis<br>**Chain:** D: **PDB Molecule:**clathrin heavy chain;<br>**PDBTitle:** clathrin d6 coat |
| 24 | c6tedQ_ | Alignment | not modelled | 89.0 | 11 | **PDB header:**transcription<br>**Chain:** Q: **PDB Molecule:**rna polymerase-associated protein ctr9 homolog;<br>**PDBTitle:** structure of complete, activated transcription complex pol ii-dsif-2 paf-spt6 uncovers allosteric elongation activation by rtf1 |
| 25 | c5jqyA_ | Alignment | not modelled | 88.1 | 8 | **PDB header:**oxidoreductase<br>**Chain:** A: **PDB Molecule:**aspartyl/asparaginyl beta-hydroxylase;<br>**PDBTitle:** aspartyl/asparaginyl beta-hydroxylase (asph)oxygenase and tpr domains2 in complex with manganese, n-oxalylglycine and factor x substrate3 peptide fragment(39mer-4ser) |
| 26 | c4b4tP_ | Alignment | not modelled | 86.3 | 15 | **PDB header:**hydrolase<br>**Chain:** P: **PDB Molecule:**26s proteasome regulatory subunit rpn5;<br>**PDBTitle:** near-atomic resolution structural model of the yeast 26s proteasome |
| 27 | c6wb92_ | Alignment | not modelled | 86.1 | 14 | **PDB header:**membrane protein<br>**Chain:** 2: **PDB Molecule:**er membrane protein complex subunit 2;<br>**PDBTitle:** structure of the s. cerevisiae er membrane complex |
| 28 | c6af0A_ | Alignment | not modelled | 85.9 | 7 | **PDB header:**transcription<br>**Chain:** A: **PDB Molecule:**ctr9 protein; |

| | | | | | | |
|---|---|---|---|---|---|---|
| 28 | [illegible] | Alignment | not modelled | [illegible] | 7 | **PDBTitle:** structure of ctr9, paf1 and cdc73 ternary complex from myceliophthora2 thermophila |
| 29 | c4cr2R_ | Alignment | not modelled | 84.3 | 7 | **PDB header:**hydrolase<br>**Chain:** R: **PDB Molecule:**26s proteasome regulatory subunit rpn7;<br>**PDBTitle:** deep classification of a large cryo-em dataset defines the2 conformational landscape of the 26s proteasome |
| 30 | c5a5tM_ | Alignment | not modelled | 82.5 | 10 | **PDB header:**hydrolase<br>**Chain:** M: **PDB Molecule:**eukaryotic translation initiation factor 3 subunit m;<br>**PDBTitle:** structure of mammalian eif3 in the context of the 43s preinitiation2 complex |
| 31 | c5nnrD_ | Alignment | not modelled | 81.9 | 12 | **PDB header:**transferase<br>**Chain:** D: **PDB Molecule:**n-terminal acetyltransferase-like protein;<br>**PDBTitle:** structure of naa15/naa10 bound to hypk-thb |
| 32 | c6ez8B_ | Alignment | not modelled | 81.8 | 8 | **PDB header:**protein binding<br>**Chain:** B: **PDB Molecule:**factor viii intron 22 protein;<br>**PDBTitle:** human huntingtin-hap40 complex structure |
| 33 | c4b4tR_ | Alignment | not modelled | 81.3 | 8 | **PDB header:**hydrolase<br>**Chain:** R: **PDB Molecule:**26s proteasome regulatory subunit rpn7;<br>**PDBTitle:** near-atomic resolution structural model of the yeast 26s proteasome |
| 34 | c2qfcB_ | Alignment | not modelled | 80.0 | 12 | **PDB header:**transcription regulation<br>**Chain:** B: **PDB Molecule:**plcr protein;<br>**PDBTitle:** crystal structure of bacillus thuringiensis plcr complexed with papr |
| 35 | c5gjqQ_ | Alignment | not modelled | 79.3 | 13 | **PDB header:**hydrolase<br>**Chain:** Q: **PDB Molecule:**26s proteasome non-atpase regulatory subunit 11;<br>**PDBTitle:** structure of the human 26s proteasome bound to usp14-ubal |
| 36 | d1elra_ | Alignment | not modelled | 78.3 | 19 | **Fold:**alpha-alpha superhelix<br>**Superfamily:**TPR-like<br>**Family:**Tetratricopeptide repeat (TPR) |
| 37 | c4b4tS_ | Alignment | not modelled | 77.3 | 13 | **PDB header:**hydrolase<br>**Chain:** S: **PDB Molecule:**26s proteasome regulatory subunit rpn3;<br>**PDBTitle:** near-atomic resolution structural model of the yeast 26s proteasome |
| 38 | c5an3B_ | Alignment | not modelled | 75.7 | 10 | **PDB header:**transcription<br>**Chain:** B: **PDB Molecule:**sgt1;<br>**PDBTitle:** structure of an sgt1-skp1 complex |
| 39 | c4uzyA_ | Alignment | not modelled | 75.3 | 10 | **PDB header:**motor protein<br>**Chain:** A: **PDB Molecule:**flagellar associated protein;<br>**PDBTitle:** crystal structure of the chlamydomonas ift70 and ift52 complex |
| 40 | c3mv3B_ | Alignment | not modelled | 74.6 | 8 | **PDB header:**protein transport<br>**Chain:** B: **PDB Molecule:**coatomer subunit epsilon;<br>**PDBTitle:** crystal structure of a-cop in complex with e-cop |
| 41 | c6hftA_ | Alignment | not modelled | 73.8 | 9 | **PDB header:**chaperone<br>**Chain:** A: **PDB Molecule:**hsp70/hsp90 co-chaperone cns1;<br>**PDBTitle:** hsp90 co-chaperone cns1 from saccharomyces cerevisiae (delta69) |
| 42 | c4kvmA_ | Alignment | not modelled | 73.3 | 15 | **PDB header:**transferase/transferase inhibitor<br>**Chain:** A: **PDB Molecule:**n-terminal acetyltransferase a complex subunit nat1;<br>**PDBTitle:** the nata (naa10p/naa15p) amino-terminal acetyltransferase complex2 bound to a bisubstrate analog |
| 43 | c1na3A_ | Alignment | not modelled | 73.2 | 19 | **PDB header:**de novo protein<br>**Chain:** A: **PDB Molecule:**designed protein ctpr2;<br>**PDBTitle:** design of stable alpha-helical arrays from an idealized tpr motif |
| 44 | c5m72A_ | Alignment | not modelled | 72.9 | 11 | **PDB header:**protein transport<br>**Chain:** A: **PDB Molecule:**signal recognition particle subunit srp72;<br>**PDBTitle:** structure of the human srp68-72 protein-binding domain complex |
| 45 | c4kbmB_ | Alignment | not modelled | 72.1 | 15 | **PDB header:**transferase/transcription<br>**Chain:** B: **PDB Molecule:**rna polymerase-binding transcription factor card;<br>**PDBTitle:** structure of the mtb card/rnap beta subunit b1-b2 domains complex |
| 46 | c3rkvA_ | Alignment | not modelled | 71.8 | 14 | **PDB header:**isomerase<br>**Chain:** A: **PDB Molecule:**putative peptidylprolyl isomerase;<br>**PDBTitle:** c-terminal domain of protein c56c10.10, a putative peptidylprolyl2 isomerase, from caenorhabditis elegans |
| 47 | c2xcbA_ | Alignment | not modelled | 71.8 | 9 | **PDB header:**protein binding<br>**Chain:** A: **PDB Molecule:**regulatory protein pcrh;<br>**PDBTitle:** crystal structure of pcrh in complex with the chaperone2 binding region of popd |
| 48 | c5wftA_ | Alignment | not modelled | 69.8 | 7 | **PDB header:**structural protein<br>**Chain:** A: **PDB Molecule:**pelb;<br>**PDBTitle:** pelb 319-436 from pseudomonas aeruginosa pao1 |
| 49 | c5mpdS_ | Alignment | not modelled | 69.5 | 14 | **PDB header:**hydrolase<br>**Chain:** S: **PDB Molecule:**26s proteasome regulatory subunit rpn3;<br>**PDBTitle:** 26s proteasome in presence of atp (s1) |
| 50 | c4i1aA_ | Alignment | not modelled | 68.5 | 12 | **PDB header:**hydrolase<br>**Chain:** A: **PDB Molecule:**response regulator aspartate phosphatase i;<br>**PDBTitle:** crystal structure of the apo form of rapi |
| 51 | c6ww7B_ | Alignment | not modelled | 68.1 | 10 | **PDB header:**membrane protein<br>**Chain:** B: **PDB Molecule:**er membrane protein complex subunit 2;<br>**PDBTitle:** structure of the human er membrane protein complex in a lipid nanodisc |
| 52 | c1wao4_ | Alignment | not modelled | 67.8 | 16 | **PDB header:**hydrolase<br>**Chain:** 4: **PDB Molecule:**serine/threonine protein phosphatase 5;<br>**PDBTitle:** pp5 structure |
| 53 | c4hnxA_ | Alignment | not modelled | 67.5 | 14 | **PDB header:**transferase<br>**Chain:** A: **PDB Molecule:**n-terminal acetyltransferase a complex subunit nat1; |

| | | | | | | |
|---|---|---|---|---|---|---|
| | | | | | | **PDBTitle:** the nata acetyltransferase complex bound to ppgpp |
| 54 | c6q8jA_ | Alignment | not modelled | 67.5 | 11 | **PDB header:**splicing **Chain:** A: **PDB Molecule:**wd40 repeat-containing protein smu1; **PDBTitle:** nterminal domain of human smu1 in complex with lsp641 |
| 55 | c3n71A_ | Alignment | not modelled | 67.3 | 8 | **PDB header:**transcription **Chain:** A: **PDB Molecule:**histone lysine methyltransferase smyd1; **PDBTitle:** crystal structure of cardiac specific histone methyltransferase smyd1 |
| 56 | c5ganJ_ | Alignment | not modelled | 66.6 | 14 | **PDB header:**transcription **Chain:** J: **PDB Molecule:**pre-mrna-splicing factor 6; **PDBTitle:** the overall structure of the yeast spliceosomal u4/u6.u5 tri-snrnp at2 3.7 angstrom |
| 57 | c4yczB_ | Alignment | not modelled | 66.4 | 9 | **PDB header:**structural protein **Chain:** B: **PDB Molecule:**nup85; **PDBTitle:** y-complex hub (nup85-nup120-nup145c-sec13 complex) from m. thermophila2 (a.k.a. t. heterothallica) |
| 58 | c3jckA_ | Alignment | not modelled | 64.0 | 17 | **PDB header:**hydrolase **Chain:** A: **PDB Molecule:**26s proteasome regulatory subunit rpn3; **PDBTitle:** structure of the yeast 26s proteasome lid sub-complex |
| 59 | c5jjtA_ | Alignment | not modelled | 63.2 | 24 | **PDB header:**hydrolase **Chain:** A: **PDB Molecule:**serine/threonine-protein phosphatase 5; **PDBTitle:** crystal structure of a type 5 serine/threonine protein phosphatase2 from arabidopsis thaliana |
| 60 | c2vq2A_ | Alignment | not modelled | 62.4 | 8 | **PDB header:**structural protein **Chain:** A: **PDB Molecule:**putative fimbrial biogenesis and twitching motility **PDBTitle:** crystal structure of pilw, widely conserved type iv pilus biogenesis2 factor |
| 61 | c5l0wB_ | Alignment | not modelled | 62.2 | 11 | **PDB header:**membrane protein **Chain:** B: **PDB Molecule:**sec72; **PDBTitle:** structure of post-translational translocation sec71/sec72 complex |
| 62 | c5l0yE_ | Alignment | not modelled | 60.9 | 12 | **PDB header:**protein transport **Chain:** E: **PDB Molecule:**sec72-ssa1 c-terminal peptide fusion protein; **PDBTitle:** crystal structure of a sec72-ssa1 c-terminal peptide fusion protein |
| 63 | c3gyzB_ | Alignment | not modelled | 59.8 | 12 | **PDB header:**chaperone **Chain:** B: **PDB Molecule:**chaperone protein ipgc; **PDBTitle:** crystal structure of ipgc from shigella flexneri |
| 64 | c3t5xA_ | Alignment | not modelled | 59.2 | 9 | **PDB header:**transcription **Chain:** A: **PDB Molecule:**pci domain-containing protein 2; **PDBTitle:** pcid2:dss1 structure |
| 65 | c2lt3A_ | Alignment | not modelled | 58.0 | 15 | **PDB header:**transcription **Chain:** A: **PDB Molecule:**transcriptional regulator, card family; **PDBTitle:** solution nmr structure of the c-terminal domain of cdnl from2 myxococcus xanthus |
| 66 | c3upvA_ | Alignment | not modelled | 57.3 | 14 | **PDB header:**peptide binding protein **Chain:** A: **PDB Molecule:**heat shock protein sti1; **PDBTitle:** tpr2b-domain:phsp70-complex of yeast sti1 |
| 67 | c6vl6U_ | Alignment | not modelled | 56.3 | 13 | **PDB header:**de novo protein **Chain:** U: **PDB Molecule:**t33_dn2b; **PDBTitle:** de novo designed tetrahedral nanoparticle t33_dn2 presenting bg5052 sosip trimers |
| 68 | c4nrhB_ | Alignment | not modelled | 54.0 | 6 | **PDB header:**chaperone/protein binding **Chain:** B: **PDB Molecule:**chaperone sycd; **PDBTitle:** copn-scc3 complex |
| 69 | c5ulmB_ | Alignment | not modelled | 50.9 | 15 | **PDB header:**transferase **Chain:** B: **PDB Molecule:**mitogen-activated protein kinase kinase kinase 5; **PDBTitle:** structure of the ask1 central regulatory region |
| 70 | c3zpjA_ | Alignment | not modelled | 50.4 | 14 | **PDB header:**unknown function **Chain:** A: **PDB Molecule:**ton_1535; **PDBTitle:** crystal structure of ton1535 from thermococcus onnurineus na1 |
| 71 | d2c2la1 | Alignment | not modelled | 50.1 | 15 | **Fold:**alpha-alpha superhelix **Superfamily:**TPR-like **Family:**Tetratricopeptide repeat (TPR) |
| 72 | c4cr2Z_ | Alignment | not modelled | 50.0 | 16 | **PDB header:**hydrolase **Chain:** Z: **PDB Molecule:**26s proteasome regulatory subunit rpn1; **PDBTitle:** deep classification of a large cryo-em dataset defines the2 conformational landscape of the 26s proteasome |
| 73 | d1a17a_ | Alignment | not modelled | 49.9 | 11 | **Fold:**alpha-alpha superhelix **Superfamily:**TPR-like **Family:**Tetratricopeptide repeat (TPR) |
| 74 | c5gjqR_ | Alignment | not modelled | 49.5 | 7 | **PDB header:**hydrolase **Chain:** R: **PDB Molecule:**26s proteasome non-atpase regulatory subunit 6; **PDBTitle:** structure of the human 26s proteasome bound to usp14-ubal |
| 75 | c5gjqZ_ | Alignment | not modelled | 49.2 | 9 | **PDB header:**hydrolase **Chain:** Z: **PDB Molecule:**26s proteasome non-atpase regulatory subunit 2; **PDBTitle:** structure of the human 26s proteasome bound to usp14-ubal |
| 76 | c3ro2A_ | Alignment | not modelled | 46.6 | 8 | **PDB header:**protein binding **Chain:** A: **PDB Molecule:**g-protein-signaling modulator 2; **PDBTitle:** structures of the lgn/numa complex |
| 77 | c5l4kZ_ | Alignment | not modelled | 46.2 | 13 | **PDB header:**structural protein **Chain:** Z: **PDB Molecule:**26s proteasome non-atpase regulatory subunit 2; **PDBTitle:** the human 26s proteasome lid |
| 78 | c4cr3S_ | Alignment | not modelled | 46.2 | 11 | **PDB header:**hydrolase **Chain:** S: **PDB Molecule:**26s proteasome regulatory subunit rpn3; **PDBTitle:** deep classification of a large cryo-em dataset defines the2 conformational landscape of the 26s proteasome |

| | | | | | | |
|---|---|---|---|---|---|---|
| 79 | d2crba1 | Alignment | not modelled | 45.6 | 24 | **Fold:**Spectrin repeat-like<br>**Superfamily:**MIT domain-like<br>**Family:**MIT domain |
| 80 | c4gyoB_ | Alignment | not modelled | 45.6 | 10 | **PDB header:**hydrolase<br>**Chain:** B: **PDB Molecule:**response regulator aspartate phosphatase j;<br>**PDBTitle:** crystal structure of rap protein complexed with competence and2 sporulation factor |
| 81 | c5cwnA_ | Alignment | not modelled | 44.0 | 12 | **PDB header:**de novo protein<br>**Chain:** A: **PDB Molecule:**designed helical repeat protein;<br>**PDBTitle:** crystal structure of de novo designed helical repeat protein dhr71 |
| 82 | c5fzqB_ | Alignment | not modelled | 43.6 | 12 | **PDB header:**unknown function<br>**Chain:** B: **PDB Molecule:**designed tpr protein;<br>**PDBTitle:** designed tpr protein m4n |
| 83 | c3vtxB_ | Alignment | not modelled | 43.5 | 12 | **PDB header:**protein binding<br>**Chain:** B: **PDB Molecule:**mama;<br>**PDBTitle:** crystal structure of mama protein |
| 84 | c5cqsC_ | Alignment | not modelled | 42.7 | 8 | **PDB header:**protein binding<br>**Chain:** C: **PDB Molecule:**elongator complex protein 1;<br>**PDBTitle:** dimerization of elp1 is essential for elongator complex assembly |
| 85 | c4a1sB_ | Alignment | not modelled | 42.7 | 6 | **PDB header:**cell cycle<br>**Chain:** B: **PDB Molecule:**partner of inscuteable;<br>**PDBTitle:** crystallographic structure of the pins:insc complex |
| 86 | c5mpeZ_ | Alignment | not modelled | 37.0 | 9 | **PDB header:**hydrolase<br>**Chain:** Z: **PDB Molecule:**26s proteasome regulatory subunit rpn1;<br>**PDBTitle:** 26s proteasome in presence of atp (s2) |
| 87 | d2fbna1 | Alignment | not modelled | 35.8 | 15 | **Fold:**alpha-alpha superhelix<br>**Superfamily:**TPR-like<br>**Family:**Tetratricopeptide repeat (TPR) |
| 88 | c2fbnA_ | Alignment | not modelled | 35.8 | 15 | **PDB header:**structural genomics, unknown function<br>**Chain:** A: **PDB Molecule:**70 kda peptidylprolyl isomerase, putative;<br>**PDBTitle:** plasmodium falciparum putative fk506-binding protein2 pfl2275c, c-terminal tpr-containing domain |
| 89 | c4cgwA_ | Alignment | not modelled | 35.6 | 18 | **PDB header:**chaperone<br>**Chain:** A: **PDB Molecule:**rna polymerase ii-associated protein 3;<br>**PDBTitle:** second tpr of spaghetti (rpap3) bound to hsp90 peptide srmeevd |
| 90 | c3urzB_ | Alignment | not modelled | 34.9 | 11 | **PDB header:**protein binding<br>**Chain:** B: **PDB Molecule:**uncharacterized protein;<br>**PDBTitle:** crystal structure of a putative protein binding protein (bacova_03105)2 from bacteroides ovatus atcc 8483 at 2.19 a resolution |
| 91 | c1ihgA_ | Alignment | not modelled | 34.4 | 12 | **PDB header:**isomerase<br>**Chain:** A: **PDB Molecule:**cyclophilin 40;<br>**PDBTitle:** bovine cyclophilin 40, monoclinic form |
| 92 | c5efrA_ | Alignment | not modelled | 34.2 | 16 | **PDB header:**cell adhesion<br>**Chain:** A: **PDB Molecule:**bama-bamd fusion protein;<br>**PDBTitle:** crystal structure of a bama-bamd fusion |
| 93 | c5gjqP_ | Alignment | not modelled | 33.4 | 11 | **PDB header:**hydrolase<br>**Chain:** P: **PDB Molecule:**26s proteasome non-atpase regulatory subunit 12;<br>**PDBTitle:** structure of the human 26s proteasome bound to usp14-ubal |
| 94 | d2buga1 | Alignment | not modelled | 33.2 | 16 | **Fold:**alpha-alpha superhelix<br>**Superfamily:**TPR-like<br>**Family:**Tetratricopeptide repeat (TPR) |
| 95 | c3fgaB_ | Alignment | not modelled | 32.6 | 13 | **PDB header:**hydrolase/hydrolase inhibitor<br>**Chain:** B: **PDB Molecule:**serine/threonine-protein phosphatase 2a 56 kda regulatory<br>**PDBTitle:** structural basis of pp2a and sgo interaction |
| 96 | c2lwjA_ | Alignment | not modelled | 32.1 | 14 | **PDB header:**transcription<br>**Chain:** A: **PDB Molecule:**transcriptional regulator, card family;<br>**PDBTitle:** nmr solution structure myxoccoccus xanthus cdnl |
| 97 | c2yinB_ | Alignment | not modelled | 31.7 | 9 | **PDB header:**apoptosis<br>**Chain:** B: **PDB Molecule:**dedicator of cytokinesis protein 2;<br>**PDBTitle:** structure of the complex between dock2 and rac1. |
| 98 | d2nppb1 | Alignment | not modelled | 31.0 | 11 | **Fold:**alpha-alpha superhelix<br>**Superfamily:**ARM repeat<br>**Family:**B56-like |
| 99 | c2kztA_ | Alignment | not modelled | 29.6 | 13 | **PDB header:**apoptosis<br>**Chain:** A: **PDB Molecule:**programmed cell death protein 4;<br>**PDBTitle:** structure of the tandem ma-3 region of pdcd4 |
| 100 | c5v8kA_ | Alignment | not modelled | 29.2 | 22 | **PDB header:**photosynthesis<br>**Chain:** A: **PDB Molecule:**p800 reaction center core protein;<br>**PDBTitle:** homodimeric reaction center of h. modesticaldum |
| 101 | c6gmhQ_ | Alignment | not modelled | 28.9 | 11 | **PDB header:**transcription<br>**Chain:** Q: **PDB Molecule:**ctr9,rna polymerase-associated protein ctr9 homolog,rna<br>**PDBTitle:** structure of activated transcription complex pol ii-dsif-paf-spt6 |
| 102 | c6i57A_ | Alignment | not modelled | 28.8 | 13 | **PDB header:**chaperone<br>**Chain:** A: **PDB Molecule:**sperm-associated antigen 1;<br>**PDBTitle:** nmr structure of the third tpr domain of the human spag1 protein |
| 103 | c2e2eA_ | Alignment | not modelled | 28.7 | 17 | **PDB header:**lyase<br>**Chain:** A: **PDB Molecule:**formate-dependent nitrite reductase complex nrfg subunit;<br>**PDBTitle:** tpr domain of nrfg mediates the complex formation between heme lyase2 and formate-dependent nitrite reductase in escherichia coli o157:h7 |
| | | | | | | **PDB header:**signaling protein<br>[illegible] |

| | | | | | | |
|---|---|---|---|---|---|---|
| 104 | c4jhrA_ | Alignment | not modelled | 28.5 | 9 | **Chain:** A: **PDB Molecule:**g-protein-signaling modulator 2;<br>**PDBTitle:** an auto-inhibited conformation of lgn reveals a distinct interaction2 mode between goloco motifs and tpr motifs |
| 105 | c5a7dB_ | Alignment | not modelled | 28.5 | 6 | **PDB header:**cell cycle<br>**Chain:** B: **PDB Molecule:**pins;<br>**PDBTitle:** tetrameric assembly of lgn with inscuteable |
| 106 | c4d10C_ | Alignment | not modelled | 28.2 | 10 | **PDB header:**signaling protein<br>**Chain:** C: **PDB Molecule:**cop9 signalosome complex subunit 3;<br>**PDBTitle:** crystal structure of the cop9 signalosome |
| 107 | c5lynA_ | Alignment | not modelled | 27.9 | 14 | **PDB header:**chaperone<br>**Chain:** A: **PDB Molecule:**small glutamine-rich tetratricopeptide repeat-containing<br>**PDBTitle:** structure of the tpr domain of sgt2 in complex with yeast ssa1 peptide2 fragment |
| 108 | c2nppE_ | Alignment | not modelled | 27.9 | 14 | **PDB header:**hydrolase/hydrolase inhibitor<br>**Chain:** E: **PDB Molecule:**serine/threonine-protein phosphatase 2a 56 kda regulatory<br>**PDBTitle:** structure of the protein phosphatase 2a holoenzyme |
| 109 | c4d0pA_ | Alignment | not modelled | 27.4 | 9 | **PDB header:**signaling protein<br>**Chain:** A: **PDB Molecule:**cop9 signalosome complex subunit 4;<br>**PDBTitle:** crystal structure of human csn4 |
| 110 | c6rnsA_ | Alignment | not modelled | 27.1 | 11 | **PDB header:**rna binding protein<br>**Chain:** A: **PDB Molecule:**gem-associated protein 5;<br>**PDBTitle:** crystal structure of the dimerization domain of gemin5 at 2.7 a |
| 111 | c5aioA_ | Alignment | not modelled | 26.6 | 14 | **PDB header:**transcription<br>**Chain:** A: **PDB Molecule:**transcription factor tau 131 kda subunit;<br>**PDBTitle:** crystal structure of t131 n-terminal tpr array |
| 112 | c3as5A_ | Alignment | not modelled | 26.6 | 13 | **PDB header:**protein binding<br>**Chain:** A: **PDB Molecule:**mama;<br>**PDBTitle:** mama amb-1 p212121 |
| 113 | c4ga0A_ | Alignment | not modelled | 26.4 | 17 | **PDB header:**transport protein<br>**Chain:** A: **PDB Molecule:**e3 sumo-protein ligase ranbp2;<br>**PDBTitle:** structure of the n-terminal domain of nup358 |
| 114 | c3rmrA_ | Alignment | not modelled | 26.1 | 16 | **PDB header:**protein binding<br>**Chain:** A: **PDB Molecule:**avirulence protein;<br>**PDBTitle:** crystal structure of hyaloperonospora arabidopsidis atr1 effector2 domain |
| 115 | c2wmoA_ | Alignment | not modelled | 24.9 | 19 | **PDB header:**cell cycle<br>**Chain:** A: **PDB Molecule:**dedicator of cytokinesis protein 9;<br>**PDBTitle:** structure of the complex between dock9 and cdc42. |
| 116 | c2katA_ | Alignment | not modelled | 23.7 | 11 | **PDB header:**structural genomics, unknown function<br>**Chain:** A: **PDB Molecule:**uncharacterized protein;<br>**PDBTitle:** solution structure of protein bpp2914 from bordetella parapertussis.2 northeast structural genomics consortium target bpr206 |
| 117 | c6px0A_ | Alignment | not modelled | 23.4 | 10 | **PDB header:**isomerase<br>**Chain:** A: **PDB Molecule:**aryl-hydrocarbon-interacting protein-like 1;<br>**PDBTitle:** crystal structure of the tpr domain of human aryl hydrocarbon2 receptor-interacting protein-like 1 (aipl1) |
| 118 | c2r5sB_ | Alignment | not modelled | 23.4 | 14 | **PDB header:**structural genomics, unknown function<br>**Chain:** B: **PDB Molecule:**uncharacterized protein vp0806;<br>**PDBTitle:** the crystal structure of a domain of protein vp0806 (unknown function)2 from vibrio parahaemolyticus rimd 2210633 |
| 119 | c5a1uC_ | Alignment | not modelled | 23.4 | 17 | **PDB header:**transport protein<br>**Chain:** C: **PDB Molecule:**coatomer subunit alpha;<br>**PDBTitle:** the structure of the copi coat triad |
| 120 | c4cgvA_ | Alignment | not modelled | 22.9 | 12 | **PDB header:**chaperone<br>**Chain:** A: **PDB Molecule:**rna polymerase ii-associated protein 3;<br>**PDBTitle:** first tpr of spaghetti (rpap3) bound to hsp90 peptide srmeevd |
