## Supplementary Figure 13 for "A conserved Guided Entry of Tail-anchored pathway is involved in the trafficking of tail-anchored membrane proteins in *Plasmodium falciparum*"

```
CLUSTAL O(1.2.4) multiple sequence alignment


PF3D7_0215600      ------------------------MDKNTLKRRSRLLQKNEARMSMLLGRNLDDEISKEK      36
HsCAML             MESMAVATDGGERPGVPAGSGLSASQRRAELRRRKLLMNSEQRINRIMGFHRPGSGAEEE      60
                                           ::.:   ** :** :.* *:. ::* :  .. ::*:

PF3D7_0215600      KESNKNDKHKKNDQHKKNDKSNQNGEDNQNGEDNQNDESNQNDEHKKNDEHKKNDEHNQN      96
HsCAML             ----------------------------------SQTKSKQQDSDKLNSLS---------      77
                                                     .: :*:*:*..* *.

PF3D7_0215600      DEHNQNDEHNQNDEHNQNDEHNQNDEHNQNDEHNQNDEHNQNDESNQNDKNRKEIPPKEE      156
HsCAML             -------------VPSVSKRVVLGDSVSTGT-TDQQGGVAEVKGTQLGDKLDSFIKPPEC      123
                                  . ...   .*. . .   :*:.   : . :: .**  . * * *

PF3D7_0215600      KDKENNPSVVMENNNNIRKDQDNKTSEHKSSTNIYNKDKNNDYNK-LLDKDDNNNNNKNI      215
HsCAML             SSD-VNLELRQRNRGDLTADSVQRGSRHGLEQYLSRFEEAMKLRKQLISEKPSQ-----E      177
                   ...  * .:  .*..::  *. :: *.*  .  : . ::  . .* *:.:. .:

PF3D7_0215600      NKNDENDFTSINNDIPNKNKITSQFIITKHEKLHFIILIILCIFISIFKVYYNNKNNLIY      275
HsCAML             DGNTTEEFDS-----------FRIFRLVGCALLALGVRAFVCKYLSIFAPFLTLQLAY--      224
                   : *  ::* *              * :.    * : :  ::* ::***  : . :

PF3D7_0215600      KKKKKGNNNLNIVQMIFFNFINSPNFFFSFSVFYNILFLLIIMLMYIKKNNITRKRIQDF      335
HsCAML             --------------MGLYKYFPKSEKKIKTTVLTAALL-----LSGIPAE----------      255
                                 * ::::: . :  :. :*:   *:     *  *  :

PF3D7_0215600      FVNMKNKLNNQNEHVFYFINNAVLCILFMGRIFKSYIISMFLINLFHDILHNYLIGVSML      395
HsCAML             -----------------VINRSMDTYSKMGEVFTDLCVYFFTFIFCHELLDYWGSEVP--      296
                                    .**.::     **.:*..  : :* : : *::*. :   *

PF3D7_0215600      QPQKVVLL       403
HsCAML             --------       296
```
